# Selection strengthens the relationship between plant diversity and the metabolic profile of *Plantago lanceolata*

**DOI:** 10.1101/2024.11.29.626026

**Authors:** Pamela Medina-van Berkum, Francesca De Giorgi, Beate Rothe, Walter Durka, Jonathan Gershenzon, Christiane Roscher, Sybille B. Unsicker

**Author notes:** Corresponding author:* Pamela Medina-van Berkum.

## Abstract

- Plants growing in biodiverse communities often increase productivity, but how plant diversity impacts the metabolome and the underlying ecological and evolutionary processes remains unclear. This study investigated how plant species diversity and selection for growing in different diversity environments affects the leaf metabolome of *Plantago lanceolata*.
- We compared the metabolites of plants derived from those that had been *selected* in the “Jena Experiment” for 17 years in plant communities with differing plant diversity with the metabolites of *naïve* plants not subjected to this selection. The metabolic profiles of *selected P. lanceolata* phytometers were also compared after growing in experimental environments varying in plant species richness, soil history, and community plant history.
- Results showed volatile compound diversity in *P. lanceolata* decreased with plant species richness, primarily due to phenotypic plasticity rather than selection. Soil history further strengthened this relationship. Conversely, non-volatile compound diversity increased with plant species richness, but only in phytometers subjected diversity-driven selection. These effects were more pronounced when plants shared soil-plant history with their community.
- In summary, our study revealed that both plastic and adaptative responses shape the metabolome of *P. lanceolata* in relation to plant diversity with these effects becoming stronger as plant and soil communities mature.

## Introduction

Concerns about the loss of global biodiversity in recent decades have intensified efforts to understand the mechanisms that mediate the relationships between biodiversity and ecosystem functioning. Experimental studies on grassland biodiversity have shown that high biodiversity promotes plant community productivity and stability (Cardinale *et al*., 2007; Allan *et al*., 2013; Wagg *et al*., 2022). These effects strengthen over time as complementary interactions between species become more important (Cardinale *et al*., 2007), leading to more pronounced relationships between biodiversity and ecosystem functioning (Reich *et al*., 2012). Although several studies on species-level responses to increased plant community diversity exist, they mostly focused on plant biomass production or plant morphological traits (Tilman *et al*., 1996; Lipowsky *et al*., 2011; van Moorsel *et al*., 2018). Only few have investigated other important plant traits, such as specialized plant metabolites (e.g. Scherling *et al*., 2010; Mraja *et al*., 2011; Zuppinger-Dingley *et al*., 2015; Ristok *et al*., 2023). This is particularly important because plant phenotypes are influenced by the synthesis and accumulation of specialized metabolites in specific organs, at various developmental stages, and in response to environmental cues.

Plant specialized (secondary) metabolites play essential roles in species interactions within communities as deterrents, toxins, attractants or signals for other organisms and in resistance to abiotic stresses (Erb & Kliebenstein, 2020). By mediating biotic interactions, they can impact plant performance and survival (Hartmann, 2007; Kessler & Kalske, 2018; Sosenski & Parra-Tabla, 2019; Erb & Kliebenstein, 2020). Specialized metabolites can be constitutive or induced, directly affecting plant antagonists or indirectly by attracting their natural enemies, thus providing a versatile defense strategy. This flexibility highlights the importance of plant chemodiversity, encompassing both the richness and composition of these chemicals, which can vary not only due to genetic differences but also in response to environmental pressures, such as herbivory and pathogens attacks or resource availability (Endara *et al*., 2023).

Previous studies have demonstrated that varying plant community diversity can induce changes in both primary and specialized metabolites in grassland species (Scherling *et al*., 2010; Mraja *et al*., 2011; Ristok *et al*., 2019). The variation in chemodiversity observed among plants of the same species may result from phenotypic plasticity or genotype selection in response to the surrounded environment (Zuppinger-Dingley *et al*., 2015). Phenotypic plasticity can take place within an organism’s lifespan in response to its environment, while evolutionary adaptations occur over a time span of a few (Rauschkolb *et al*., 2022) to many generations (Nicotra *et al*., 2010). Previous research has reported both plastic and adaptative responses at the chemical level. For instance, Zuppinger-Dingley *et al*. (2015) found that for several grassland species the different selection pressures in low or high diversity communities led to adaptation in plant chemical traits over several generations. On the other hand, Miehe-Steier *et al*. (2015) showed that for *Plantago lanceolata L.* (ribwort plantain), the production of iridoid glycosides is a plastic response to the surrounding plant community. This means that individuals of the same species growing in communities of varying diversity might show differences in their chemical traits due to their different environments and associated (a)biotic selective pressures.

Soil communities modify the biotic and abiotic environment of plants, while plants create belowground legacies by altering the soil’s biotic and abiotic properties. This mutual interaction, known as plant-soil feedback (van der Putten *et al*., 2013), can influence plant defense by modulating the plant chemodiversity (Huberty *et al*., 2020; Ristok *et al*., 2023). Moreover, plant-soil feedback is an important selective driver in plant communities, hence influencing the micro-evolutionary processes in plants (Dietrich *et al*., 2021; De Giorgi *et al*., 2024).

Despite the importance of plant metabolites in the establishment, development, and survival of plants, there is a lack of knowledge on how different environments can shape metabolic profiles that are heritable and adapted. This scarcity of studies hinders our understanding of how chemical traits respond to selective pressures. Long-term biodiversity studies enable the examination of whether plant diversity effects on the plant metabolome are due to adaptation of plant species to different environments of origin or phenotypic plasticity to the actual growth environment.

Using the short-lived perennial plant *Plantago lanceolata* L. as a model species, we designed a phytometer experiment in a long-term grassland biodiversity experiment (The Jena Experiment; Roscher *et al*., 2004) to study the impact of selection and community history on metabolic responses to plant diversity. We performed two experiments: (1) *Selection Experiment* in which we compared *selected* phytometers (offspring of plants that underwent the selection pressures in the biodiversity experiment for 17 years) planted in their environment of origin with *naïve* phytometers (offspring of plants that did not experience this biodiversity selection). (2) *Community History Experiment*, where phytometers with selection history were transplanted back into their environment of origin as well as in two other experimental environments, specifically communities with the same species composition but differing in the community history, with one having only soil history and the other lacking both soil and plant history. After one year, we analysed the leaf metabolomes of these individuals. We hypothesize that (1) *naïve* phytometers will experience more leaf damage than *selected* phytometers and thus, exhibit greater antagonist-induced metabolic diversity irrespective of the surrounding plant diversity. In addition, *naïve* phytometers will have a weaker response to plant diversity compared to *selected* phytometers. (2) Community soil-plant history will strengthen the relationship between plant diversity and the plant metabolome.

## Material & Methods

### Field site

The study was conducted in the Jena Experiment (Jena, Germany; 50°55 ’ N, 11°35 ’ E; 130 m a. s. l.), a long-term grassland biodiversity experiment established in 2002 in Jena, Germany (Roscher *et al*., 2004). We performed the study in the ΔBEF Experiment (Determinants of Long-Term Biodiversity Effects on Ecosystem Functioning), established in 2016 (see Vogel *et al*., 2019 for more details). We selected 12 communities (12 plots) where *Plantago lanceolata* L. (ribwort plantain) belonged to the sown species combinations, covering a gradient in species richness from a *P. lanceolata* monoculture to a 60 plant species-mixture (1, 2, 4, 8, 16, and 60 plant species).

### Selection Experiment

To explore the effects of experimental selection on *P. lanceolata* metabolic profiles, we used *P. lanceolata* phytometers with two seed origins. (1) *Selected seeds*: Seeds collected from *P. lanceolata* individuals growing in experimental communities that had experienced differing biodiversity conditions for 17 years ranging from monoculture to 60 plant species-mixture. (2) *Naïve seeds*: phytometers obtained by growing plants from the initial seed batches (Rieger-Hofmann) used to establish the Jena Experiment 2002, whose ancestors did not experience the environment of the biodiversity experiment (Fig. 1). Both types of phytometers, *naïve* and *selected*, were transplanted into the 17 years-old plant communities when they were ten weeks-old (details below).

**Fig. 1.**
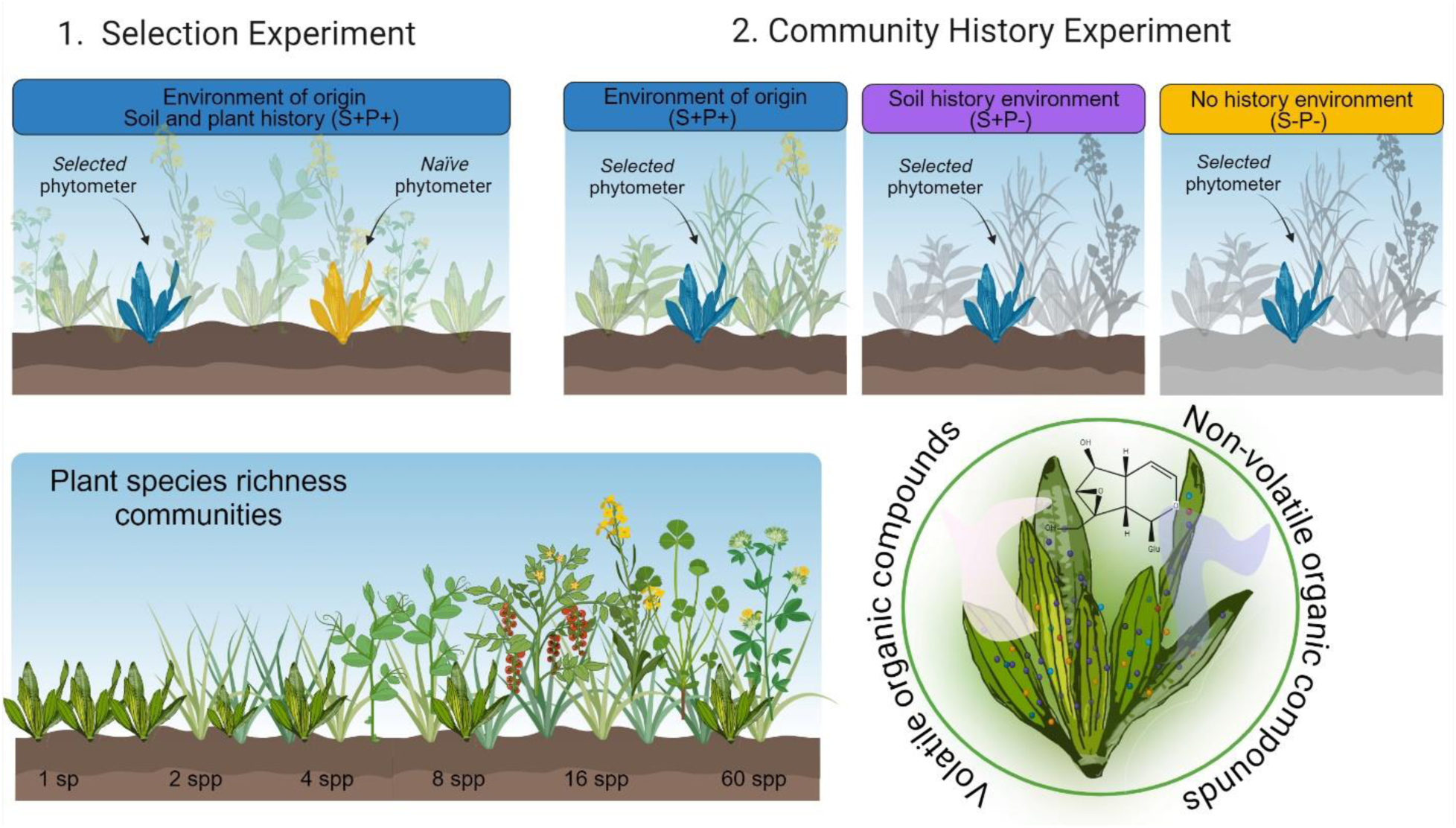
Graphical illustration of the *Selection Experiment* and the *Community History Experiment* with *Plantago lanceolata* phytometers across the plant diversity gradient of the Jena Experiment, Germany. The *Selection Experiment* compared the metabolic profiles between *selected* and *naïve* plants. *Selected* plants (offspring of plants that underwent the selection pressures of varying biodiversity environments in the Jena Experiment) were planted in their environment of origin with the *naïve* plants (offspring of plants that did not experience biodiversity selection of the Jena Experiment). The *Community History Experiment* compared the metabolic profiles of *selected* plants grown in different experimental environments based on the ΔBEF experiment established in 2016 (Vogel *et al*., 2019). *No history* environment (S-P-): soil layer and plant community removed. *Soil history* environment (S+P-): only plant community removed. In both treatments, new plot-specific plant mixtures were sown in 2016. *Soil-Plant history* environment (S+P+): environment of origin, same as core area established in 2002 (long-term control). Headspace analysis of volatile organic compounds and metabolic profiles of leaf extracts were analyzed after one year of transplantation in communities with a plant diversity gradient from *P. lanceolata* monoculture to a 60 plant species-mixture (1, 2, 4, 8, 16, and 60 plant species). Figure was created with Biorender.

### Community History Experiment

To explore the effects of community history on *P. lanceolata* metabolomic profiles, we used *P. lanceolata* phytometers from seeds collected in the 17-year-old communities (same *selected* seeds used for the *Selection Experiment*) and transplanted into the ΔBEF treatments of their original community (Fig. 1). The ΔBEF experiment consisted of three subplots (1.5 m x 3 m) inside the main experimental plots, with different degrees of community history. (1) *Soil and plant history* (S+P+): 17-year-old plant communities, long-term control (from where the seeds were collected, their environment of origin). (2) *Soil history* (S+P-): experimental environment in which plant species were removed while keeping the soil and resowing plot-specific plant species. (3) *No history* (S-P-): experimental environment in which soil and plant layer were removed and replaced with arable field soil resown with plot-specific plant species (see Vogel *et al*., 2019 for more details).

### Preparation and establishment of phytometer plants

During summer 2018, *P. lanceolata* plants were obtained from the germination of seeds originally used to establish the Jena Experiment, which had been stored at –20°C since 2002. These seedlings were grown in a greenhouse, then transplanted to a seed bed outdoors in autumn at the Experimental Field Station Bad Lauchstädt (Germany). A year later, seeds from these plants were collected and called *naïve* seeds. *Selected* seeds were collected in 2019 from four mothers of *P. lanceolata* growing in the 17-year-old communities (with soil and plant history), cleaned and stored at room temperature until the start of the experiment. In January 2020, single seeds (*selected* and *naïve*) were germinated in cells of QuickPot trays (Hermann Meyer KG, Rellingen, Germany) filled with autoclaved soil from the field site mixed with sterile mineral sand (25 vol%) in a greenhouse (temperature of 18°C: 12°C with 14 hours of day light). After eight weeks, the trays were moved into an open greenhouse with outside light and temperature conditions for two weeks to harden the plants before being planted in the field. In early April 2020, when the phytometers were ten weeks old, *selected* phytometers were transplanted into the same plot where the seeds were initially collected, while the *naïve* ones were transplanted into all of the plots (see (see De Giorgi *et al*., 2024 for more details).

### Sampling and measurements

One year after the transplantation (August 2021), we simultaneously measured morphological and chemical traits of the phytometer plants of both experiments in the field. These included the collection of headspace volatile emissions, assessment of phenotypic traits, analysis of non-volatile leaf metabolites, and calculation of percentage of leaf damage. Six individuals per treatment were designated for the measurements of phenotypic traits and non-volatile leaf metabolites. From these, four individuals where designated to collect the headspace volatile compounds and assess leaf damage.

### Phenotypic traits and leaf damage

For phenotypic traits, we recorded the reproductive status, leaf biomass, plant height, and leaf greenness according to De Giorgi *et al*. (2024). Leaf damage by herbivores and pathogens was assessed using the method described by Unsicker and Mody (2005). This involved reconstructing the original leaf area in digital photographs taken of both sides from the leaves after the harvest using Adobe Photoshop 2020 (Adobe, California, USA). Damage was quantified as percentage of the total leaf area (cm^2^).

### Headspace volatile collection

Volatile organic compounds (VOC) emitted by *P. lanceolata* phytometers were collected and measured using the protocol described in Medina-van Berkum et al. (2024) with few modifications. In brief, VOC emission of individual plants was captured using a closed push-pull system over a two-hour period. The plants were enclosed in PET bags (Bratschlauch, Toppits, Germany), and charcoal-filtered air was continuously pumped into these bags at a flow rate of 1 L/min. VOC traps, consisting of 25 mg of Porapak absorbent (ARS, Grainville, FL, USA) inserted in Teflon tubes, was attached to the bags and air was pumped out through them at a flow rate of 0.6 L/min. All volatile collections were performed between 9:00 am and 1 pm. After collection, the traps were eluted with 200 µl of dichloromethane containing nonyl acetate (Sigma-Aldrich, 10 ng/µl) as an internal standard. The eluted VOCs were analyzed using an Agilent (Santa Clara, CA, USA) 6890 series gas chromatograph (GC) coupled to either an Agilent 5973 series mass spectrometer (MS) for identification or to a flame ionization detector (FID) for quantification (see Medina-van Berkum *et al*., 2024 for more details). VOC identification was achieved by comparing GC-MS spectra with reference spectra from the Wiley and National Institute of Standards and Technology libraries, as well as by comparing retention times and mass spectra to those of standards from our collection. VOC quantification was determined from GC-FID data based on the peak area in relation to the peak area of the internal standard. The relative response factor was computed with authentic standards or estimated with the effective carbon number concept, and normalized to leaf fresh weight and duration of collection (nanogram per gram FW per hour).

### Metabolite extraction from leaves

Leaf samples were flash frozen in liquid nitrogen after the harvesting, lyophilized and ground to fine powder by agitating them together with a mix of stainless-steel balls (2-4mm in diameter) in a paint shaker. Then, 10 mg of leaf powder was extracted with 100% methanol (0.1 mL per mg) containing D6-salicylic acid (SA), D6-jasmonic acid (JA) and D6-abscisic acid (ABA) as internal standards (Sigma-Aldrich). Aliquots of raw extracts were used for (1) untargeted metabolite profiling and (2) targeted analysis of phytohormones, iridoid glycosides and phenylpropanoid glycosides.

### Metabolome profiling

Untargeted metabolic profiles of leaves were obtained by ultra-high performance liquid chromatography coupled *via* electrospray ionization (ESI) to a qTOF mass spectrometer (UHPLC-ESI-HRMS) in negative ionization mode. The mobile phase consisted of 0.1% v/v formic acid in water and in acetonitrile. Raw data files from UHPLC-HRMS were transferred to the Metaboscape® (Bruker) software to perform the bucketing based on MS1 spectra. Quality control (QC) samples were prepared by pipetting equal volumes of all the samples in a designated LC-MS vial for analysis and run every 40 samples together with the blanks. Raw data acquisition was carried out as previously described by Medina-van Berkum *et al*. (2024). The processed LC-MS/MS data were then used for the *in-silico* prediction of chemical taxonomic classification using the CANOPUS package (Dührkop *et al*., 2021) from the SIRIUS software (Dührkop *et al*., 2019), considering only classified features with a probability of at least 70% at pathway level.

### Quantification of targeted compounds

Quantification of targeted compounds was conducted using an HPLC-MS/MS system (HPLC 1260 Infinity II [Agilent Technologies, Santa Clara, USA]—QTrap® 6500+ [AB Sciex, Waltham, Massachusetts, USA]) in multiple reaction monitoring (MRM) mode, following Medina-van Berkum *et al*. (2024) with few modifications. Phytohormones were quantified with authentic standards (D6-JA, D6-ABA, D6-SA). For iridoid glycosides and verbascoside, quantification was based on comparison to external authentic standards curves (aucubin: Carl Roth, Germany; catalpol: Wako, Japan; verbascoside: Extrasynthese, France).

### Data analysis

We performed mixed-model analysis and linear discriminate analysis to test the effect of both selection history and community history on leaf traits, leaf damage and leaf metabolome of *P. lanceolata*. The data of the two experiments, *Selection Experiment* and *Community History Experiment*, were analyzed separately. Leaf metabolome diversity for both volatile and non-volatile compounds was calculated based on Hill numbers using VOC concentration and peak intensity of features as abundance. To test the effect of biodiversity selection in the *Selection Experiment*, we fitted “species richness” (SR; log2 transformed sown diversity), “selection” (S; factor with two levels: *selected* vs. *naïve*) and their interactions (SR *x* S) as fixed effects. Plot identity nested in block was fitted as random effect. To test the effect of community history in the *Community History Experiment*, we performed a similar model with environment instead of selection as fixed effect (E; factor with three levels; S-P-no history, S+P-with soil history only, S+P+ soil and plant history). We started with a null model with the random effects only, and successively added the fixed effects with species richness first, followed by treatment (selection or environment) and interactions. To investigate the presence of legumes in shaping the selection and community history effect, we created another model by fitting legumes (presence/absence) before or after species richness. Since previous analyses have shown that vegetation height varies either with sown species richness or depending on environment history (De Giorgi *et al*., 2024), this could be a potentially explain the effects of both factors. Therefore, we performed another model in which we included mean height of the surrounding vegetation as a co-variable fitted before the experimental factors. All models were fitted with maximum likelihood (ML), and Wald tests were used to decide on the significance of the fixed effects. When needed, data were transformed to meet the assumptions of normality and homogeneity of variances. To identify non-volatile metabolic features significantly affected by the treatments, we used generalized linear mixed models (Gaussian log link) based on the previously mentioned model structures. The significance of fixed effects (p < 0.05) was assessed by Wald tests, followed by false discovery rate (FDR) adjustment for p-values.

The analyses and visualization were conducted in R version 4.3.3 (R Core Team, 2024) using the following packages: rBExIS, dplyr (Wickham *et al*., 2023), tidyverse (Wickham *et al*., 2019), tibble (Müller & Wickham, 2023) and janitor (Firke, 2023) for data retrieve, cleaning and formatting; lme4 (Bates *et al*., 2015), lmerTest (Kuznetsova *et al*., 2017), glmmTMB (Brooks *et al*., 2017), performance (Lüdecke *et al*., 2021), mixOmics (Rohart *et al*., 2017) and hillR (Li, 2018) for statistical and diversity analysis; notame (Klavus *et al*., 2020) for filtering false positive signals of untargeted metabolites; and ggplot2 (Wickham, 2016), ggeffects (Lüdecke, 2018) and pheatmap (Kolde, 2019) for graphical visualization. Graphics were enhanced with Adobe Illustrator CC 2021.

## Results

### Effects of selection and community history on plant performance and leaf damage

We found that only *selected* phytometers (*P. lanceolata* that were offspring of plants that underwent the selection pressures of varying biodiversity environments in the Jena Experiment) showed an increase of shoot biomass with increasing species richness of the community, while *naïve* phytometers (offspring of plants that did not experience these biodiversity selection pressures) had similar shoot biomass across diversity gradient (SR *x* S: *x^2^* = 4.95, *p* = 0.026; Fig. **2a**; Table S1). Moreover, this pattern was only true when *selected* phytometers grew in their environment of origin (SR *x* E: *x^2^* = 9.86, *p* = 0.007; Fig. **2e**; Table S2) and not when community history had been eliminated by removing surrounding soil and plants or plants alone. Leaf greenness and leaf length did not differ between *naïve* and *selected* phytometers (Fig. **2b, c**). However, phytometers increased their leaf greenness when they grew in communities with legumes compared to non-legumes communities (*x^2^* = 9.11, *p* = 0.003; Table S1). In the *Community History Experiment*, we found that leaf length increased with increasing species richness regardless of the environment treatment (*x^2^* = 4.71, *p* = 0.03, Fig. **2g**). Similar to the *Selection Experiment* results, the presence of legumes in the plot enhanced leaf greenness in *P. lanceolata* (*x^2^* = 13.68, *p* < 0.001; Table S2) while it decreased with increasing vegetation height in their surroundings (*x^2^* = 7.56, *p* = 0.006; Table S2).

**Fig. 2.**
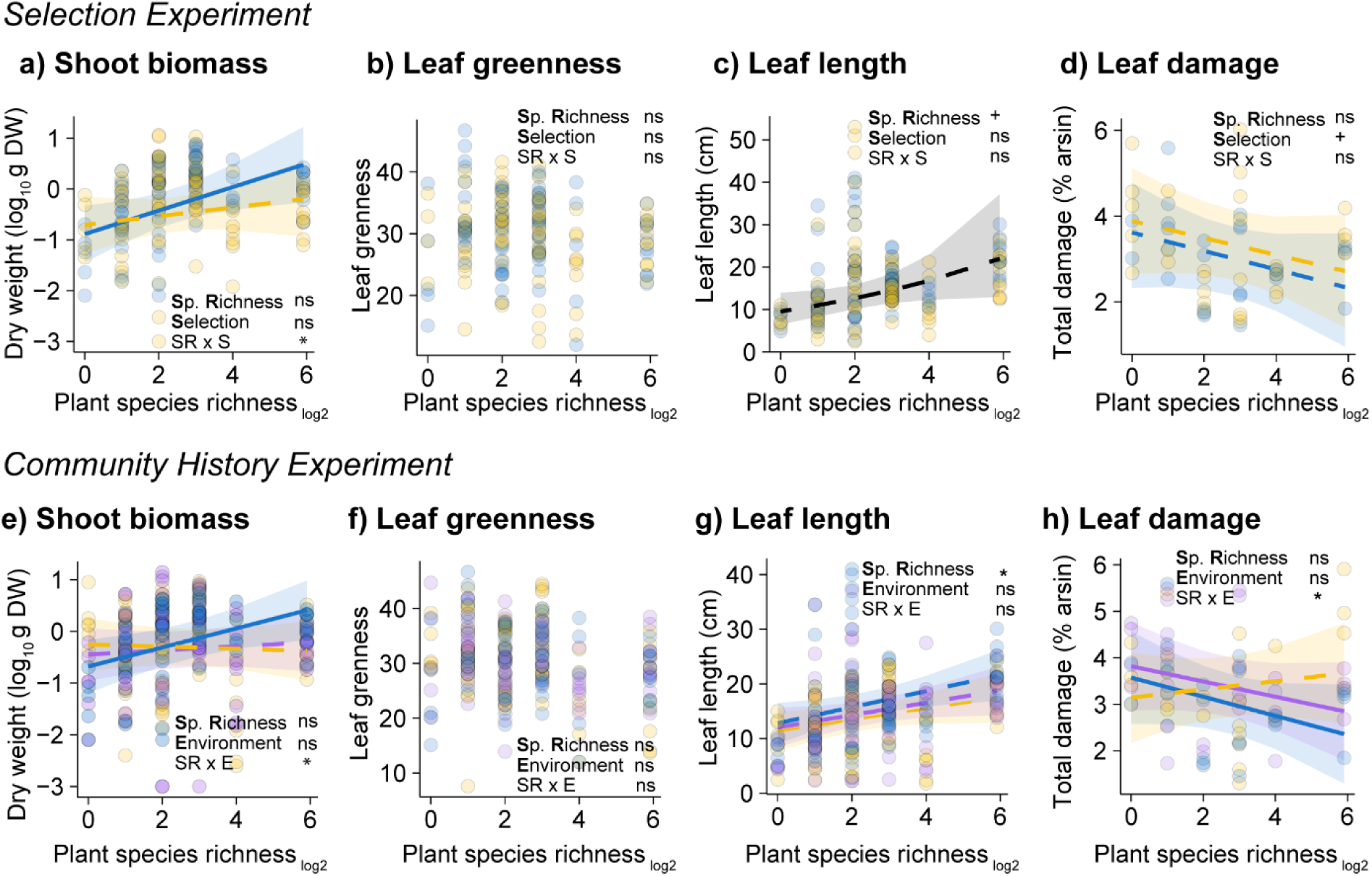
Effects of selection history and community history on leaf traits and leaf damage of *Plantago lanceolata* across a plant diversity gradient. *Selection Experiment* (top section): shoot biomass, leaf greenness, leaf length and total leaf damage (herbivore + pathogen damage) of *naïve* (yellow dots; offspring of the original seed material used in establishment of the Jena Experiment) and *selected* (blue dots; offspring of plants that experienced selection pressures in the biodiversity experiment) phytometers across a plant diversity gradient. *Community History Experiment* (bottom section): shoot biomass, leaf greenness, leaf length and total leaf damage of *selected* phytometers grown in three different environments: their environment of origin (blue dots; soil and plant history, S+P+), in the environment with soil history only (purple dots; S+P-) or no history environment (yellow dots; S-P-) across a plant diversity gradient. The different environment treatments are based on the ΔBEF experiment established in 2016 (Vogel *et al*., 2019). Lines represent predictions from linear mixed-effects models. Solid lines denote significant species richness relationship (p < 0.05) and dashed lines show non-significant relationship. Points represent each phytometer. Asterisks indicate significant effects (ns = no significant, +p < 0.1; *p < 0.05; **p < 0.01; ***p < 0.001) on species richness (SR), selection history (S), community history (E) or their interactions (SR x S or SR x E). Selection Experiment: N = 112 and 56, for leaf traits and leaf damage respectively. Community History Experiment: N = 169 and 80, for leaf traits and leaf damage respectively.

*Naïve* phytometers had a tendency to experience higher leaf damage compared to the *selected* ones (*x^2^* = 2.77, p = 0.096; Fig. **2d**). When phytometers grew in a no history environment (S-P-), leaf damage increased with increasing species richness (SR *x* E: *x^2^* = 6.19, *p* = 0.045; post hoc S-P-vs S-P+ and S+P+: < 0.05; Fig. **2h**). These patterns were primarily driven by changes in pathogen damage across species richness rather than herbivory damage (Table S2). Additionally, an increase in vegetation height in the surroundings reduced pathogen leaf damage of *P. lanceolata* phytometers (Table S1).

### Volatile and non-volatile leaf metabolome profiles

A total of 31 volatile organic compounds (VOC) were identified from the headspace volatile collection of *P. lanceolata* in the field (Table S3). These VOCs were categorized into green leaf volatiles (GLVs) (5), aromatics (4), homoterpenes (1), monoterpenes (6), sesquiterpenes (8), and nine other compounds not classified into these groups. Sesquiterpenes represented the most diverse group, while monoterpenes and GLVs were the most abundant.

Overall, we detected 2,263 features in leaf extracts of non-volatile compounds from *P. lanceolata* analyzed by untargeted LC-MS measurements in the negative ionization mode with 49% of the features putatively annotated by CANOPUS. Based on these *in-*silico classification, the terpenoid and shikimate-phenylpropanoid pathways were the most dominant pathways in the leaf metabolome of *P. lanceolata* (Fig. **3a, b**). Iridoid monoterpenoids constituted the most abundant class (72%) within terpenoids, largely due to the high number of iridoid glycosides, such as aucubin and catalpol, which are two of the most abundant examples reaching up to 30 mg per g DW based on targeted analyses (Fig. **3c, d**). Phenylpropanoids comprised with 42% to the metabolic features and were thus the most diverse class within the shikimates and phenylpropanoids pathway. Verbascoside and plantamajoside are two of the most abundant phenylpropanoids in *P. lanceolata* leaves reaching up to 50 mg per g DW (Fig. **3e, f**).

**Fig. 3.**
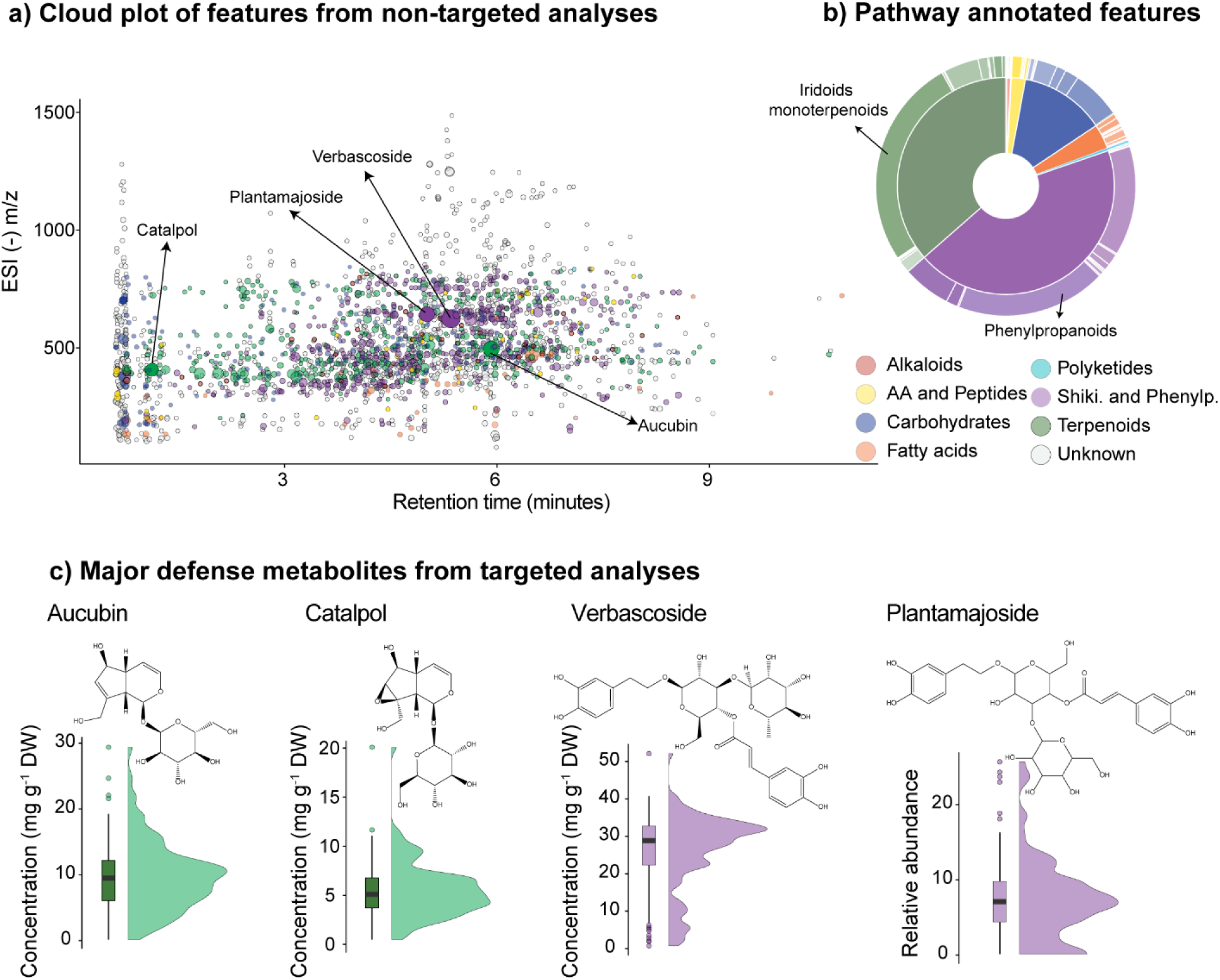
Profiles of non-volatile leaf metabolites of *Plantago lanceolata* phytometers after one year of transplantation in the Jena Experiment. a) Cloud plot of metabolic features from untargeted LC-MS measurements made in the negative ionization mode. Features are color-coded based on putative biosynthetic pathway classification; size of the circles represent their intensity; b) Pie chart with the number of features classified by biosynthetic pathway. A total of 2,263 were detected with 49% of the features putatively annotated. Biosynthetic pathway classification was performed with CANOPUS on Sirius platform. c) Boxplot and violin plots of concentrations of aucubin, catalpol, verbascoside and relative abundance of plantamajoside in the leaves of all phytometers measured by targeted LC-MS analysis.

### Plant metabolome responses to selection history

Overall, VOC profiles overlapped between *naïve* and *selected P. lanceolata* phytometers (Fig. **4a**). Total emission was unaffected by plant species richness or selection history (Table S4). Nevertheless, when considering the vegetation height of the surroundings, sesquiterpene emission decreased as plant species richness in the community increased (*x*^2^ = 6.93, *p* = 0.008; Table S4). Accounting for vegetation height also revealed that both VOC richness (Hill q0) and diversity (Hill q1) decreased with increasing plant species richness in the community (richness: *x^2^* = 4.72, *p* = 0.030; diversity: *x^2^* = 3.97, *p* = 0.048; Fig. **4b, c**). *Selected* phytometers had a tendency to exhibit higher numbers of VOCs (hill q0) compared to *naïve* phytometers (*x^2^* = 3.20, *p* = 0.074; Fig. **4b**).

**Fig. 4.**
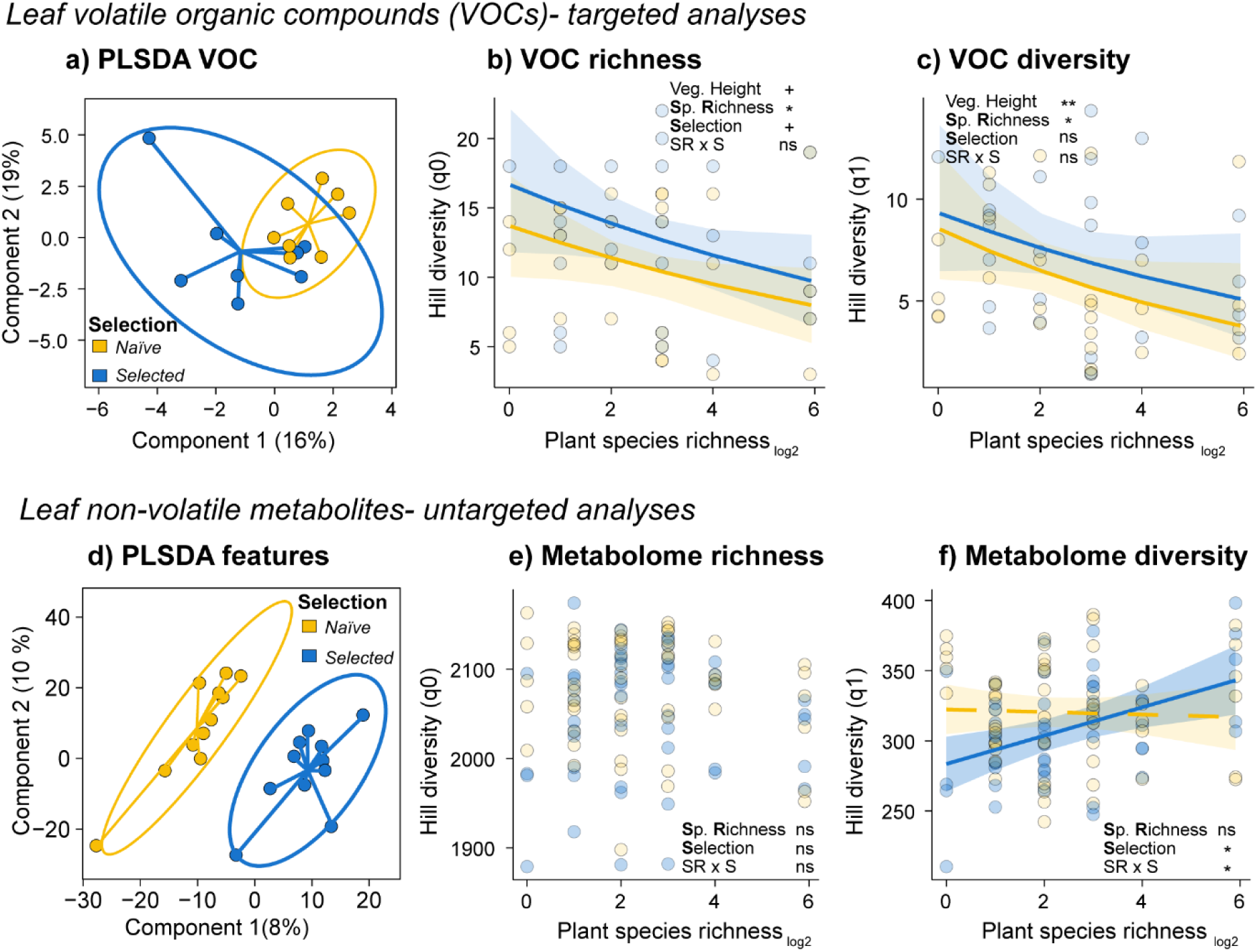
*Selection Experiment:* Effects of selection history on leaf metabolite profiles of *Plantago lanceolata* across a plant diversity gradient. *Leaf VOCs* (top section): Overall, a total of 31 VOCs was identified. a) Partial least square discriminant analysis (PLS-DA) of the VOC profile in *naïve* (offspring of the original seed material used for the establishment of the Jena Experiment) and *selected* (offspring of plants that underwent the selection pressures in the biodiversity experiment) phytometers. The results are presented as principal component score plots, with each point in the plot representing a mean value of phytometers in a community; b) VOC richness (hill richness-q0) and c) VOC diversity (Hill Shannon- q1) across a plant richness gradient. *Leaf non-volatile metabolites* (bottom section): There were 2263 metabolic features in the non-targeted analysis in the negative ionization mode after bucketing and filtering. d) Partial least square discriminant analysis (PLS-DA) of metabolic features, b) Metabolome richness (hill richness-q0), c) Metabolome diversity (hill Shannon-q1) of *naïve* and *selected* phytometers across a plant diversity gradient. Lines represent predictions from linear mixed-effects models. Solid lines denote significant species richness relationship (p < 0.05) and dashed lines show non-significant relationship. Points represent each phytometer. Asterisks indicate significant effects (ns= no significant, +p < 0.1; *p < 0.05; **p < 0.01; ***p < 0.001) on species richness (SR), selection history (S) or their interaction (SR *x* S). N = 60 and 108, for VOC and metabolic feature profiles, respectively.

In terms of non-volatile compounds, we observed that *naïve* and *selected P. lanceolata* phytometers displayed distinctly different metabolic profiles (Fig. **4d**). The number of metabolic features was not influenced by species richness (*x^2^* = 0.19, *p* = 0.662; Fig. **4e**); however, considering feature intensity (Hill q1 and q2), we found that metabolite diversity increased with increasing species richness in *selected* phytometers, while the metabolite diversity of *naïve* phytometers was not affected by plant species richness (SR *x* S: Hill q1: *x^2^* = 6.35, *p* = 0.012; Hill q2: *x^2^* = 7.90, *p* = 0.005; Fig. **4f**). Vegetation height in their surrounding did not influence the overall metabolite richness and diversity of *P. lanceolata* individuals (Table S5).

Overall, 689 metabolic features in *P. lanceolata* leaf extracts differed significantly in their intensity across diversity gradient, selection history or their interaction (34% of the whole metabolome) including a mixture of features from different metabolic classes, mainly terpenoids and shikimates/phenylpropanoids (Fig. S1). Species richness had an impact on 224 unique features (10% of whole metabolome), of which 65% decreased in intensity with increasing species richness (Fig. S1). Selection history affected 18% of the metabolome, where the majority of features had higher intensity in *naïve* phytometers compared to *selected* phytometers. Furthermore, 9% of the features were affected by the interaction between species richness and selection history, suggesting that these features in *selected* phytometers reacted differently to species richness than they did in *naïve* phytometers. The increased vegetation height of the surrounding plant community in high diversity mixtures influenced 11% of the metabolome. Additionally, the presence of legumes influenced the feature intensity in response to species richness and selection, both positively and negatively (Fig. S1).

Based on these results, we quantified some of the main defense hormones in *P. lanceolata* and the best-known anti-herbivore defense compounds, the iridoid glycosides aucubin and catalpol and the phenylpropanoid glycosides verbascoside and plantamajoside (Table 1, Table S6). Considering the vegetation height of the surrounding plant community, we found that the concentration of aucubin, verbascoside and plantamajoside decreased with increasing species richness regardless of the selection history (aucubin: *x^2^* = 9.34, *p* = 0.002, verbascoside: *x^2^* = 4.65, *p* = 0.031, plantamajoside: *x^2^* = 4.96, *p* = 0.026). The negative relationship of verbascoside foliar concentration with species richness was stronger in communities with legumes compared to the ones without legumes. Aucubin concentrations increased with increasing vegetation height of the surrounding plant community (*x^2^* = 8.12, *p* = 0.004, Table 1). For defensive hormones, species richness influenced salicylic acid (SA) and abscisic acid (ABA) concentrations, though they did not affect the levels of jasmonates overall (Table 1, but see Table S6 for specific patterns of each jasmonates). The concentration of SA and ABA decreased with increasing species richness (SA: *x^2^* = 7.60, *p* = 0.005; ABA: *x^2^* = 3.54, *p* = 0.048). However, the decrease of ABA concentrations was driven by the increasing vegetation height of the surrounding plant community (*x^2^* = 9.35, *p* = 0.002, Table 1).

**Table 1.**
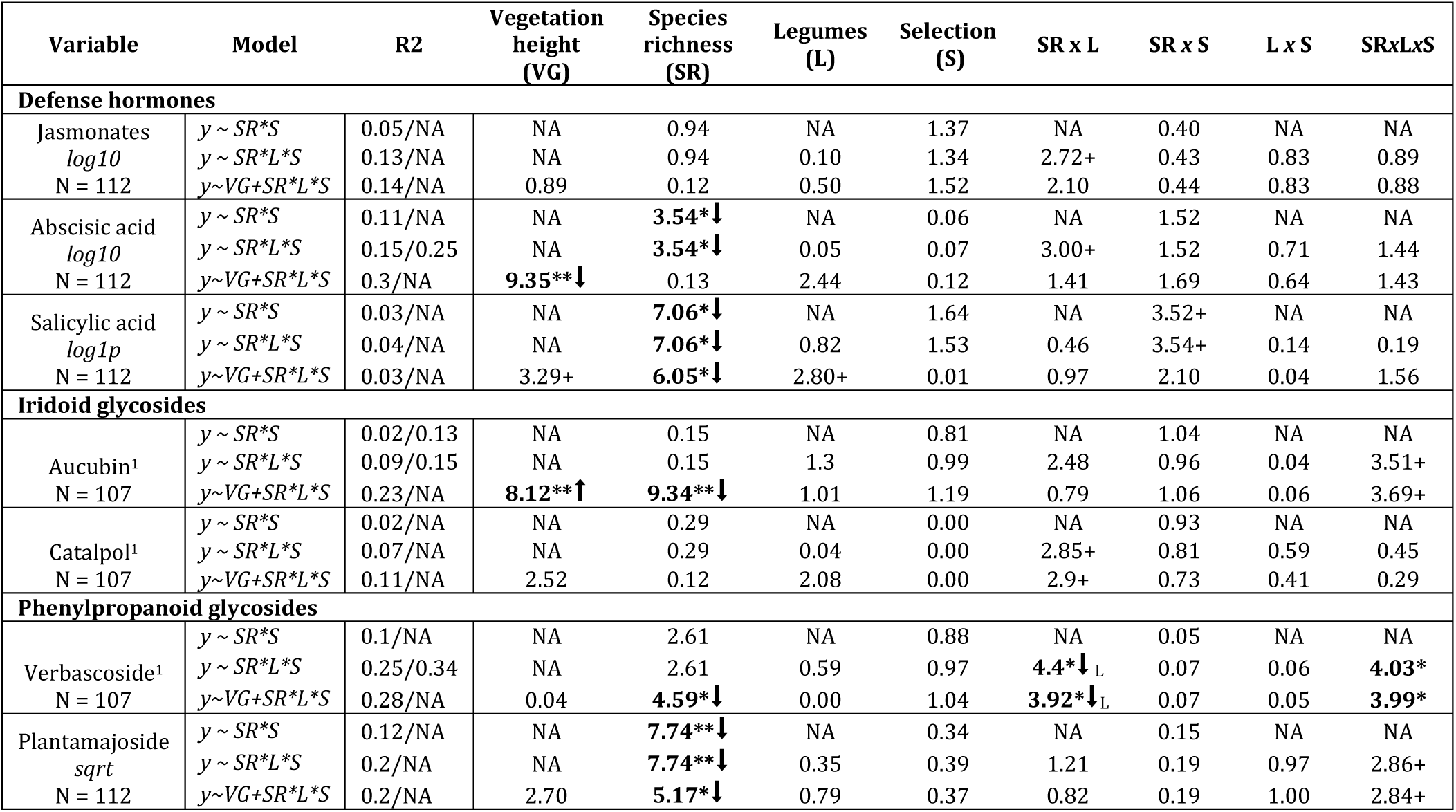
*Selection Experiment:* Wald chi-squared analysis of variance (ANOVA) results for the linear mixed models of *P. lanceolata* hormones and other non-volatile defense compounds quantified by targeted analyses. The table reports marginal and conditional R^2^ (marginal before slash and conditional after), the number of samples, and x^2^-values for each model (rows). Level of significances is based on p-values and reported with asterisks and dots: *** p < 0.001; ** p < 0.01; * p < 0.05, [x^2^] p<0.1. Arrows next to the x^2^-values indicate the patterns: increase (⬆) or decrease (⬇) in relation to the fixed factor (column).^1^Five samples were lost during the LC-MS quantification of the compounds.

### Plant metabolite profile responses to community history

VOC profiles of *selected P. lanceolata* did not vary among the different environments (Fig. **5a**). Nevertheless, phytometers in environments without history (S-P-) displayed greater similarity in VOC profiles across the species richness gradient compared to those in environments with soil history (S+P+, S+P-). While total emission was not influenced by environment, monoterpene emission decreased with increasing species richness only in environments with soil history (S+P+, S+P-: SR *x* E: *x*^2^ = 8.53, *p* = 0.014; post-hoc < 0.05; Table S7). Emission of sesquiterpene and aromatic compounds decreased with species richness when considering the surrounding vegetation height (sesquiterpene: *x*^2^ = 6.84, *p* = 0.008; aromatic: *x*^2^ = 4.68, *p* = 0.031; Table S7). Neither species richness nor community history influenced directly the number of VOC (Hill q0) in *P. lanceolata* (Fig. **5b**). However, accounting for vegetation height revealed that VOC diversity (Hill q1) decreased with increasing species richness only in phytometers that grew in environments with soil history (S+P+, S+P-), but remained similar across the species richness gradient in environments without history (S-P-; SR *x* E: *x^2^* = 5.13, *p* = 0.048, Fig. **5c**).

**Fig. 5.**
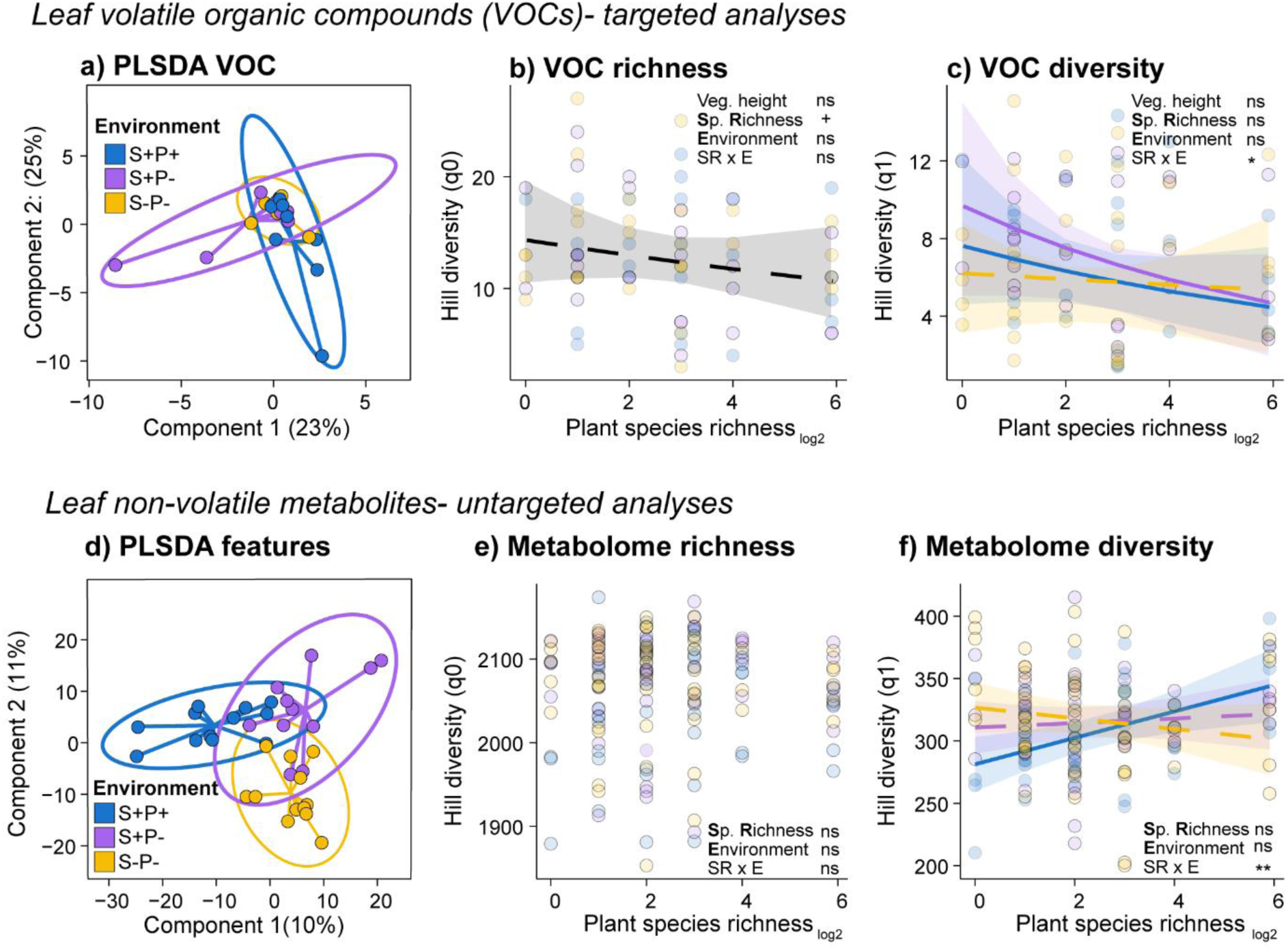
*Community History Experiment:* Effects of community history on volatile and non-volatile leaf metabolites of *Plantago lanceolata* across a plant diversity gradient. *Leaf VOCs* (top section): Overall, a total of 32 VOCs were identified. a) Partial least square discriminant analysis (PLS-DA) of VOCs from *selected* phytometers that grew in their environment of origin (soil-plant history, S+P+), in the environment with soil history (S+P-) or no history environment (S-P-). The results are presented as principal component score plots, with each point in the plot representing a mean value of phytometers in a community; b) VOC richness (hill richness-q0) and c) VOC diversity (Hill Shannon-q1) across a plant richness gradient. *Leaf non-volatile metabolites* (bottom section): There were 2263 metabolic features in the negative ionization mode after bucketing and filtering. d) Partial least square discriminant analysis (PLS-DA) of the metabolites of *selected* phytometers that grew in their environment of origin, in the environment with soil history or no history environment. The results are presented as principal component score plots, with each point in the plot representing a plot, e) Metabolome richness (hill richness-q0), and f) Metabolome diversity (Hill Shannon-q1) across a plant richness gradient. Lines represent predictions from linear mixed-effect models. Solid lines denote significant species richness relationship (p < 0.05) and dashed lines show a non-significant relationship. Points represent each phytometer. Asterisks indicate significant effects (ns= no significant, +p < 0.1; *p < 0.05; **p < 0.01; ***p < 0.001) on species richness (SR), community history (E) or their interaction (SR x E). N = 87 and 156, for VOC and metabolic feature profiles, respectively.

The non-volatile leaf metabolic composition of *selected P. lanceolata* phytometers was influenced by the experimental environment in which they grew (Fig. **5d**). The number of non-volatile metabolic features did not differ among environment treatments (*x^2^* = 0.42, *p* = 0.809; Fig. **5e**). However, when considering the intensity of these features, we found a significant interaction between species richness and community history. Specifically, metabolic diversity increased with increasing species richness only in phytometers growing in environments of origin (S+P+), whereas for phytometers in the other experimental environments (S+P-, S-P-) the response to species richness was weaker (Hill q1: *x^2^* = 10.28, *p* = 0.006; Hill q2: *x^2^* = 8.19, *p* = 0.017; post hoc < 0.01; Fig. **5f**, Table S8). Additionally, the surrounding vegetation height did not influence the overall metabolome richness and diversity of *P. lanceolata* phytometers (Table S8).

Overall, 830 features significantly differed in intensity among the experimental environments (37% of the whole metabolome). Phytometers growing in the environment of origin had lower intensity for most of their features compared to phytometers growing in the other experimental environments (Fig. S3). Species richness had an impact on 374 features (16% of whole metabolome), of which 52% decreased in intensity with increasing species richness (Fig S4). Regarding the impact of community history (16% of the features), we discovered that the majority of features intensified when phytometers were in experimental environments without history (S-P-, S+P-). Moreover, we found that 12% of the features had an interaction effect between species richness and experimental environment, primarily due to different strength in responses of phytometers in the environment with soil and plant history (S+P+) and no history (S-P-; Fig. S4).

While foliar concentration of catalpol did not change across the species richness gradient, aucubin concentration decreased as plant species richness increased when vegetation height in the surrounding was taken into consideration (*x^2^* = 6.32, *p* = 0.012). Catalpol concentration was higher in environments with soil and plant history (*x^2^* = 6.27, *p* = 0.043), however this effect became a tendency when we considered the vegetation height of the surrounding (*x^2^* = 9.77, *p* = 0.002). Considering the presence of legumes, verbascoside concentration increased with species richness in the presence of legumes but decreased in communities without legumes, regardless of the experimental environment (*x^2^* = 9.47, *p* = 0.002, Table 2). On the other hand, the relative abundance of plantamajoside decreased with species richness only in communities with legumes (*x^2^* = 9.07, *p* = 0.002, Table 2, Table S9).

**Table 2.** *Community History Experiment:* Wald Chi-Squared Analysis of Variance (ANOVA) results for the linear mixed models of *P. lanceolata* plant defense hormones and other non-volatile defense compounds quantified by targeted analyses. The table reports marginal and conditional R^2^ (marginal before slash and conditional after), the number of samples, and x^2^-values for each model (rows). Level of significances is based on p-values and reported with asterisks and dots: *** p < 0.001; ** p < 0.01; * p < 0.05, + p<0.1. Arrows next to the x^2^-values indicate the patterns: increase (⬆) or decrease (⬇) in relation to the fixed factor (column).

| Compounds | Model | R<sup>2</sup><sub>m/c</sub> | Vegetation height (VG) | Species richness (SR) | Legumes (L) | Environment (E) | SR x L | SR x E | L x E | SR x L x E |
| --- | --- | --- | --- | --- | --- | --- | --- | --- | --- | --- |
| **Defence hormones** |  |  |  |  |  |  |  |  |  |  |
| Jasmonates<br>*log10*<br>N = 163 | $y \sim SR^*E$ | 0.05/0.28 | NA | 0.83 | NA | 1.87 | NA | 4.00 | NA | NA |
| | $y \sim SR^*L^*E$ | 0.14/0.27 | NA | 0.83 | 0.11 | 1.84 | **4.45\***↓ | 4.03 | 1.75 | 2.09 |
| | $y \sim VG+SR^*L^*E$ | 0.19/NA | 2.27 | 2.32 | 0.37 | 0.93 | 3.71+ | 3.45 | 1.18 | 2.36 |
| Absciscic acid<br>*log10*<br>N = 163 | $y \sim SR^*E$ | 0.07/NA | NA | 1.36 | NA | 1.12 | NA | 5.94+ | NA | NA |
| | $y \sim SR^*L^*E$ | 0.1/NA | NA | 1.36 | 0.03 | 1.12 | 2.20 | 5.99+ | 0.38 | 0.27 |
| | $y \sim VG+SR^*L^*E$ | 0.11/NA | 0.58 | 0.85 | 0.01 | 1.28 | 2.44 | **6.38\***↓s | 0.35 | 0.53 |
| Salicylic acid<br>*log10*<br>N = 163 | $y \sim SR^*E$ | 0.02/NA | NA | **5.59\***↓ | NA | 1.81 | NA | 0.79 | NA | NA |
| | $y \sim SR^*L^*E$ | 0.02/NA | NA | **5.59\***↓ | 0.01 | 1.82 | **8.82\*\***↓ | 0.91 | 1.21 | **10.62\*\***↓ |
| | $y \sim VG+SR^*L^*E$ | 0.02/NA | 3.43+↑ | 3.56+ | 0.23 | 1.3 | **10.46\*\***↓ | 1.17 | 1.46 | **10.89\*\***↓ |
| **Iridoid glycosides** |  |  |  |  |  |  |  |  |  |  |
| Aucubin<br>N = 163 | $y \sim SR^*E$ | 0.03/0.12 | NA | 0.94 | NA | 1.94 | NA | 0.55 | NA | NA |
| | $y \sim SR^*L^*E$ | 0.06/0.14 | NA | 0.94 | 1.98 | 1.84 | 0.04 | 0.50 | 0.12 | 3.90 |
| | $y \sim VG+SR^*L^*E$ | 0.13/NA | 3.56+↑ | **6.32\***↓ | 0.11 | 2.63 | 0.00 | 0.31 | 0.30 | 3.75 |
| Catalpol<br>N = 163 | $y \sim SR^*E$ | 0.07/0.12 | NA | 1.19 | NA | **6.27\*** | NA | 2.54 | NA | NA |
| | $y \sim SR^*L^*E$ | 0.1/0.16 | NA | 1.19 | 0.43 | **6.26\*** | 0.54 | 2.51 | 0.46 | 4.07 |
| | $y \sim VG+SR^*L^*E$ | 0.17/NA | **9.77\*\***↑ | 0.58 | **4.14\*** | 4.84+ | 0.60 | 1.82 | 0.77 | 2.76 |
| **Phenylpropanoid glycosides** |  |  |  |  |  |  |  |  |  |  |
| Verbascoside<br>N = 163 | $y \sim SR^*E$ | 0.06/NA | NA | 0.61 | NA | 2.77 | NA | 3.85 | NA | NA |
| | $y \sim SR^*L^*E$ | 0.25/0.33 | NA | 0.61 | 0.47 | 2.73 | **8.22\*\***↓ | 3.72 | 1.03 | 5.24+ |
| | $y \sim VG+SR^*L^*E$ | 0.28/NA | 0.05 | 0.97 | 0.28 | 2.00 | **10.04\*\***↓ | 3.30 | 0.90 | **6.47\***↓ |
| Plantamajoside<br>N = 163 | $y \sim SR^*E$ | 0.05/NA | NA | 0.67 | NA | 2.99 | NA | 3.43 | NA | NA |
| | $y \sim SR^*L^*E$ | 0.28/NA | NA | 0.67 | 0.00 | 2.99 | **9.07\*\***↓ | 3.54 | 1.91 | **11.04\*\***↓ |
| | $y \sim VG+SR^*L^*E$ | 0.28/NA | 1.11 | 0.17 | 0.08 | 2.58 | **9.07\*\***↓ | 3.69 | 1.89 | **10.65\*\***↓ |

Considering the defense hormones, we observed that jasmonates concentration, in most cases, remained unaffected by species richness or experimental environment (Table 2, Table S9). On the other hand, SA decreased with increasing species richness (*x^2^* = 5.59, *p* = 0.018, Table 2). ABA decreased with increasing species richness only in *P. lanceolata* phytometers in environments with soil and plant history when considering the surrounding vegetation height (S+P+*; x^2^* = 6.38, *p* = 0.041, Table 2). Considering the presence of legumes in the model, SA concentration decreased with increasing species richness in non-legume communities. In plots with legumes, SA only increased with species richness in the environment with no history environment (S-P-), whereas in other environments there was no effect of species richness (Table 2).

## Discussion

Selection pressures from interspecific interactions between plants and other organisms may select for plants with traits that promote coexistence. In a recent study, De Giorgi *et al*. (2024) demonstrated that selection history plays a crucial role in enhancing the performance of several grassland species within high-diversity plant communities. To explore whether these selection effects extend to metabolic diversity, we focused on *P. lanceolata* phytometers and examined their metabolic changes, both volatile and non-volatile compounds, across a plant diversity gradient. We found that volatile diversity of *P. lanceolata* decreased with increasing plant species richness in the surrounding community, while their non-volatile metabolic diversity increased. However, volatile diversity responded to the increase of plant species richness diversity through plasticity rather than selection. Moreover, soil history enhanced the decrease of VOC diversity with species richness. In contrast, non-volatile diversity only increased with increasing plant species richness in individuals that underwent plant diversity-driven selection. The effects were more pronounced when plants shared soil-plant history with their community. In summary, our findings indicate that 17 years of selection history in the biodiversity experiment induced both plastic and adaptative responses in the metabolome of *P. lanceolata* in relation to plant species diversity with these effects strengthening over time as the soil and plant community aged.

### Plant diversity affects the composition and diversity of leaf metabolome

Earlier studies have shown that grassland species displayed metabolic variation in response to the diversity or plant identity of their surrounding community (Scherling *et al*., 2010; Kigathi *et al*., 2013; Kigathi *et al*., 2019). Similar results have been observed in *P. lanceolata,* where species richness directly and indirectly influenced the foliar concentration of major defense compounds (Mraja *et al*., 2011). Our results align with these studies, demonstrating that both volatile and non-volatile compounds in leaves responded to increasing plant diversity in their surroundings. Interestingly, the response of these compounds exhibited opposing patterns: volatile diversity decreased with increasing species richness in the community while non-volatile diversity increased.

An increase in community plant species richness leads to alterations in light and nutrient availability, competition with neighbors, and herbivore or pathogen load (Roscher *et al*., 2008; Ebeling *et al*., 2014; Rottstock *et al*., 2014; Bachmann *et al*., 2018; Kigathi *et al*., 2019). Light availability can strongly modulate the biosynthesis of compounds in plants, through a range of mechanisms, such as photosynthesis, light wavelength, photoperiod, and carbon and nitrogen allocations (Liu *et al*., 2023). Moreover, the presence of N_2_-fixing legumes in the community enhances soil nitrogen availability (Hartwig, 1998; Roscher *et al*., 2010). In *P. lanceolata,* light and nutrient availability have been identified as crucial players driving the variation in main defense compounds (Mraja *et al*., 2011; Miehe-Steier *et al*., 2015). We observed that VOC diversity decreased with increasing plant diversity, when considering the vegetation height of the surrounding community; i.e., sesquiterpene emissions were reduced in tall vegetation. Additionally, communities with legumes showed an overall increase in VOC emissions and richness compared to those without legumes. While the overall diversity of the non-volatile metabolome was not affected by vegetation height, some specific metabolic features were influenced by both vegetation height and the presence of legumes (affecting 10% and 7% of features detected in negative ionization mode, respectively). These findings align with previous research (Scherling *et al*., 2010), indicating that certain metabolic features are sensitive to light and nutrient availability, thereby confirming these factors as key drivers of plant metabolome profiles.

Another key driver of the plantś metabolome is antagonistic pressure, which can vary depending on the surrounding plant community. Low-diversity plant communities tend to accumulate and be dominated by plant antagonists above- and below-ground (Thakur *et al*., 2021). Previous studies have shown that *P. lanceolata* plants experienced higher leaf damage in low-diversity plant communities compared to high-diversity plant communities (Lipowsky *et al*., 2011), although some studies found no effect of plant diversity (Mraja *et al*., 2011). In our study, we did not observe a reduction of herbivory damage in plants growing in high-diversity plant communities; instead, with increasing diversity, we observed a reduction of leaf pathogen damage, which followed a similar pattern to salicylic acid (SA) concentration, though there was no significant correlation between them. Plant pathogens are typically categorized into biotrophs and necrotrophs based on their lifestyles. SA is usually induced upon biotrophic leaf pathogens, while jasmonic acid (JA) and ethylene depended responses are triggered by necrotrophic pathogens (Glazebrook, 2005). The lack of significant correlation between SA or JA levels and leaf pathogen damage was expected. This is likely because phytohormone levels reflect short-term responses, while pathogen damage accumulated over the survey period. Furthermore, the fungi’s life history is unknown, and leaf damage may not accurately reflect the actual pathogen load in the leaves.

While we found a positive relationship between metabolome diversity and plant species richness, we found that this relationship was primarily driven by chemical evenness rather than chemical richness. More specifically, *P. lanceolata* individuals in low-diversity plant communities showed a decreased emission of VOCs with increasing plant diversity. This led to reduced diversity in both number of compounds and emission of VOCs. Conversely, in low-diversity plant communities, plants exhibited increased intensity of several non-volatile metabolic features, leading to a reduction in chemical evenness. Dominant defense compounds, such as aucubin and verbascoside, decreased in concentration as plant diversity increased. This observation is consistent with the resource dilution hypothesis (Otway *et al*., 2005), which suggest that plants in high-diversity plant communities experience reduced herbivore damage and pathogen pressure, leading to decreased investment in defense compounds. This might be also explained by associational effects, in which *P. lanceolata* might benefit from being surrounded by plants with different chemical profiles (Hambäck *et al*., 2014).

Overall, in low-diversity plant communities, plants emit highly diverse VOC bouquets but display low non-volatile metabolic diversity, which is driven by high concentrations of major defense compounds. In contrast, in high-diversity plant communities, plants decrease VOC diversity, but increase non-volatile diversity due to having a more even composition. This suggests a shift in defense strategies between low-diversity and high-diversity plant communities. As community diversity increases, plants interact with a broader range of organisms, both within and across species. These interactions involve diverse metabolites that play active roles. The variation in defense strategies and responses to species diversity observed in our experiments align with the Interaction Diversity Hypothesis (Wetzel & Whitehead, 2020) or the Common-Sense Scenario (Berenbaum & Zangerl, 1996). Both perspectives propose that plant chemodiversity is influenced by intricate multi-species interactions, which simultaneously drive and reflect their chemical complexity. Further research is needed to determine if these differences represent a diversity-mediated transition from direct to indirect defense strategies.

### Plantago lanceolata exhibits both phenotypic plasticity and adaptations at metabolic level

Plant diversity can create differential selection pressures between low-diversity and high-diversity plant communities (Zuppinger-Dingley *et al*., 2015). These pressures influence plant phenotypes through both plasticity and genetic processes, affecting traits at morphological and chemical level, which in the end leads to better performance (Defossez *et al*., 2021; Thon *et al*., 2024). *Plantago lanceolata* performance, measured by total aboveground biomass, was better in *selected* phytometers compared to the *naïve* ones in their environment of origin. As a consequence, *naïve* and *selected* phytometers in low-diversity plant communities had similar biomass, but in high-diversity plant communities, *selected* phytometers had higher biomass compared to *naïve* ones. In other words, *selected* phytometers benefit from species rich communities.

We hypothesized that if there is a diversity-inflicted selection pressure at the plant metabolome level, we would observe a relationship between species richness and metabolic diversity only in *selected* phytometers, while *naïve* phytometers would display a similar metabolic diversity along a plant diversity gradient. Our results showed metabolic differences between *naïve* and *selected* phytometers growing in the same environment, but these changes were primarily evident in non-volatile profiles, while volatile profiles were similar between *naïve* and *selected* phytometers. Specifically, volatile diversity decreased with increasing plant species richness, regardless of the selection history of the phytometers. On the other hand, non-volatile diversity showed a positive relationship with plant diversity only in *selected* phytometers. These results suggest that plants exhibits both phenotypic plasticity and adaptations in their leaf metabolome.

Plants emit VOCs constitutively, but most of their VOC diversity originates from induced responses to biotic and abiotic stress. This phenotypic plasticity allows plants to communicate with beneficial organisms (like predators and parasitoids of insect herbivores) and detrimental ones and to send messages to other conspecifics and parts of the same plant. To archive this, plants must be able to recognize and differentiate between different neighbors and adjust their phenotype accordingly (Dicke, 2016). Therefore, it is more likely that plants exhibit a plastic response in VOC emissions rather than an adaptive response, especially since they are essential for intra- and interspecific communication within their surroundings. On the other hand, our study revealed that non-volatile metabolic diversity showed a positive relationship with plant diversity only in *selected* phytometers. Interestingly, when examining specific compounds, we found that the variation in iridoid glycosides and verbascoside in *P. lanceolata* across a diversity gradient appears to be driven primarily by phenotypic plasticity rather than by the selection of genotypes better fitting to specific plant diversity community, pattern previously reported (Miehe-Steier *et al*., 2015). Although previous studies have shown that the production of iridoid glycosides is heritable (Marak *et al*., 2000), we did not observe significant differences between *naïve* and *selected* phytometers. This suggests that diversity-driven selection pressures may not significantly affect the production of these compounds. Instead, the changes in their concentrations likely reflect a plastic response to the surrounding diversity.

Natural selection at the metabolome level occurs when metabolites which provide benefits become more abundant, while those that impose fitness costs become less abundant (Thon *et al*., 2024). Despite the targeted metabolome showing diversity-driven responses, our non-targeted analysis showed that features influenced by either species richness, selection history or environment history, did not belong to a specific class but rather a mix of several classes of compounds. This finding reinforces the idea that changes in plant metabolomes are complex, resulting from interacting responses among metabolic features not confined to a single class of compounds. This underscores the need to be careful in interpreting compound classes always as functional classes, when seeking explanations for plant defenses responses. Additionally, it is important to consider that a plant is simultaneously exposed to (a)-biotic factors, further supporting the concept of multivariate changes at the metabolomic level.

The interplay between phenotypic plasticity and adaptive changes at the metabolome level has been observed in previous studies. Research has highlighted that *P. lanceolata* demonstrates both plastic and adaptive capabilities in response to varying environmental conditions, particularly in its morphological and chemical traits (Bischoff *et al*., 2006; Skinner & Stewart, 2014; Medina-van Berkum *et al*., 2024). However, the degree of these responses varies depending on the specific traits studied.

### Community history enhanced diversity-driven responses at metabolic level

The impact of biodiversity on plant performance increases with ecosystem “age”, as plant and soil processes change over time (Guerrero-Ramírez *et al*., 2017; Meyer *et al*., 2017; Huang *et al*., 2018; Vogel *et al*., 2019). Given our findings that plants experienced selection pressures driven by plant diversity at the metabolome level, we further explored whether these effects were influenced by the history of the soil or plant community, based on the ΔBEF experiment (Vogel *et al*., 2019). Here we provide evidence that plant diversity-driven responses at the metabolic level are enhanced as the communities mature, promoting a greater plasticity and adaptive responses to the increase of plant species richness in the community.

Our findings indicate that emission of VOC of *P. lanceolata* showed stronger plasticity responses when plants shared history with the soil community, either only soil history or both soil and plant history. As a response to differences in plant diversity, the assembly of biotic communities and changes in soil nutrient availability over time create a history (Eisenhauer *et al*., 2024). Previous studies have shown that soil biota can significantly influence the emission and composition of volatiles in plants, involving both beneficial and detrimental ones (Fontana *et al*., 2009; Hammerbacher *et al*., 2019). Therefore, it is likely that the negative relationship between VOC and species richness strengthens over time, by the modulation of microbe-mediated soil history relationships.

A previous study showed soil legacy effects in plant metabolome (Ristok *et al*., 2019). Although the overall metabolome composition was similar among the plants growing in different environments, we found that metabolic diversity in *selected* phytometers growing in their original environment had a stronger positive response to plant diversity compared to those in environments where they did not share community plant history. These results support the idea that the impact of biodiversity on plant performance increases with ecosystem age (Eisenhauer *et al*., 2024), not only at the level of morphological plant traits and plant performance but also at the chemical level.

## Conclusion

In summary, our study has revealed a clear effect of plant diversity on *P. lanceolata* metabolome profiles, revealing contrasting responses between volatile and non-volatile compound diversity. As species richness in the surrounding environment increases, volatile diversity declines, whereas non-volatile diversity takes the opposite trajectory, showing an increased. Moreover, our findings highlight the complex interplay between plasticity and adaptation in plant responses to their environment. While VOC emissions primarily show plasticity in response to species diversity in the surrounding community, non-volatile compound production seems to involve both plastic and adaptative responses. Additionally, we demonstrated that plant and soil histories play critical roles in shaping plant metabolic responses to biodiversity over time. As soil and plant community mature, these effects seem to intensify both the plastic and adaptive responses of plants to their surrounding communities, emphasizing the dynamic of plant interactions at the metabolomic level. Further research is needed to disentangle the contributions of these mechanisms and to understand how they shape plant interactions within diverse ecological communities.

## Acknowledgments

We thank the gardeners, technicians and numerous student helpers for maintaining field site and Anne Ebeling for coordinating the Jena Experiment. We thank Nils Gottschaldt, Klara Beier-Heuchert, Christiana Voy and Melanie Werlich for their help during the field experiments. This work was supported by the Deutsche Forschungsgemeinschaft (DFG) Research Unit (RO2397-10, DU404-15 and UN276/4-1 in1in FOR 5000).

## Author contributions

PMB, FDG, WD, CR and SBU designed the research. PMB and FDG performed the field experiment. PMB and BR performed the chemical analysis. PMB analyzed the data and wrote the first draft of the manuscript. JG, CR, WD, FDG and SBU review and edited the manuscript. All authors discussed the results, contributed substantially to the drafts and gave final approval of the manuscript prior to the submission.

## Data and code availability

The data and R code will be publicly available through the Jena Experiment database (https://jexis.idiv.de). The R codes (ID = 671), leaf traits (ID= 656), leaf damage (ID = 656), volatile organic compounds (ID = 665), defense compounds based on targeted analysis (ID = 666) and processed metabolome data (ID = 668-670) will be available upon acceptance. Raw metabolome data will be available upon acceptance from MetaboLights (Yurekten et al. 2024; www.ebi.ac.uk/metabolights/MTBLS11792); Study Identifier: MTBLS11792.

