## Supplemental table for "Selection strengthens the relationship between plant diversity and the metabolic profile of *Plantago lanceolata*"

Figure S3. Upset plot of the interactions of 634 features whose intensity were significantly influenced by vegetation height, species richness, community history or their interaction.	5

**Figure S1. Upset plot of the interactions of features whose intensity were significantly influenced by vegetation height, species richness, presence of legumes, selection history or their interactions.**

Features positive correlated with the fixed factor (top section) a) Model 1: species richness x selection; b) Model 2: species richness x legumes x selection; c) Model 3: vegetation height x species richness x legumes x selection.

Features negative correlated with the fixed factor (bottom section) a) Model 1: species richness x selection; b) Model 2: species richness x legumes x selection; c) Model 3: vegetation height x species richness x legumes x selection. The blue or read bars (bottom left) indicate the number of features whose intensity was significantly influenced by each fixed factor. The blue dots (connected with black lines) represent the intersections of features by each factor, and the bars (top) indicate the frequency of these intersections. Features were putatively classified by biosynthetic pathway. Effect of the treatments to the intensity was tested with generalized linear mixed regression model.

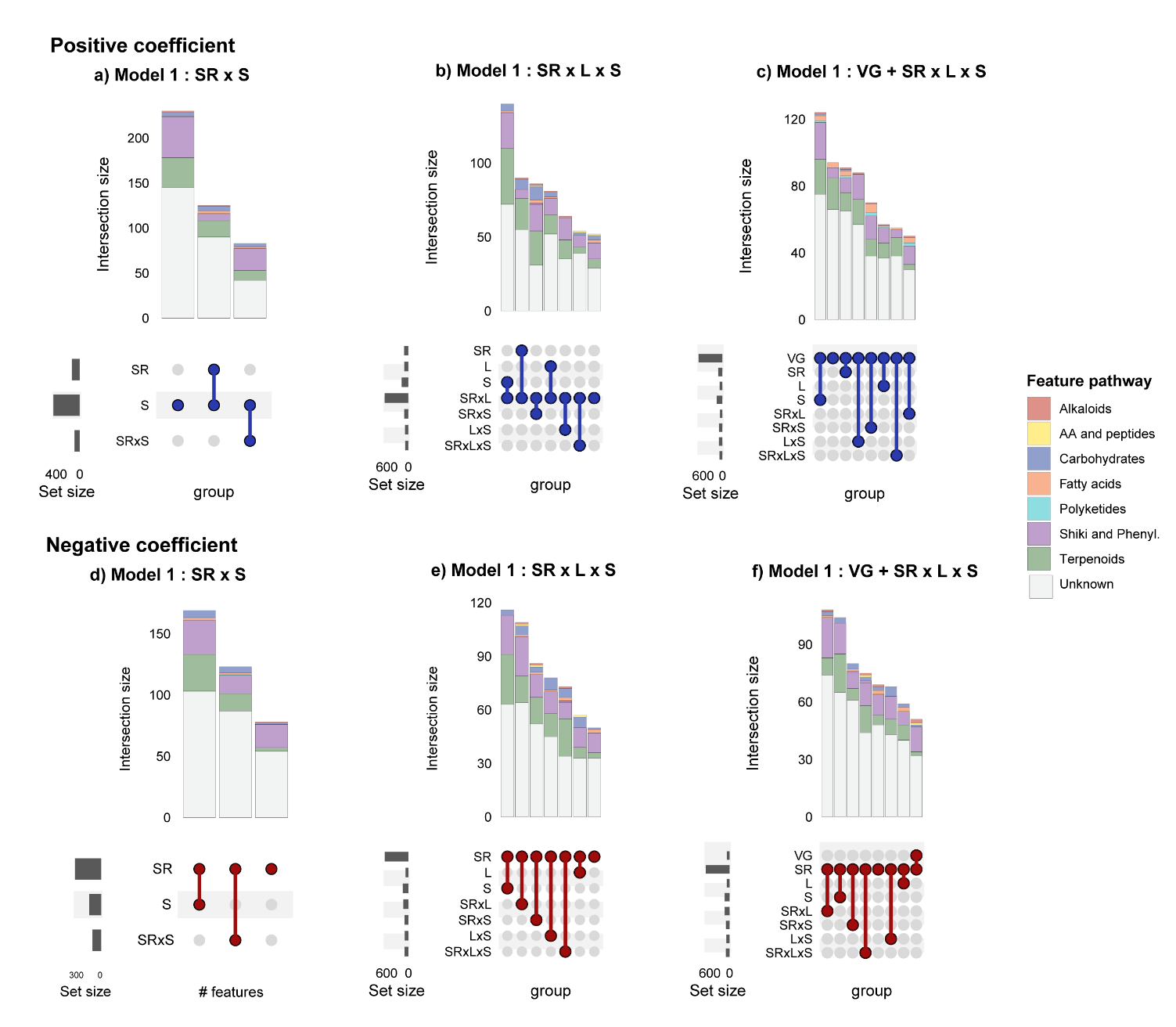

**Figure S2** **Heatmap of leaf metabolic features in *Plantago lanceolata* significantly influenced by species richness and community history.**

Effect of the treatments to the intensity was tested with generalized linear mixed regression model. The community history experiment compared the metabolomic profiles of selected plants grown in different environment treatments based on the δBEF experiment established in 2016(Vogel et al., 2019). No history (NH: –SH-PH): soil and plant layer removed, Soil history (SH: +SH-PH): only plant layer removed. In both treatments, new plot-specific plants species-mixtures were sowed. Soil-Plant history (SPH: +SH+PH): as control, same as core area stablished in 2002.

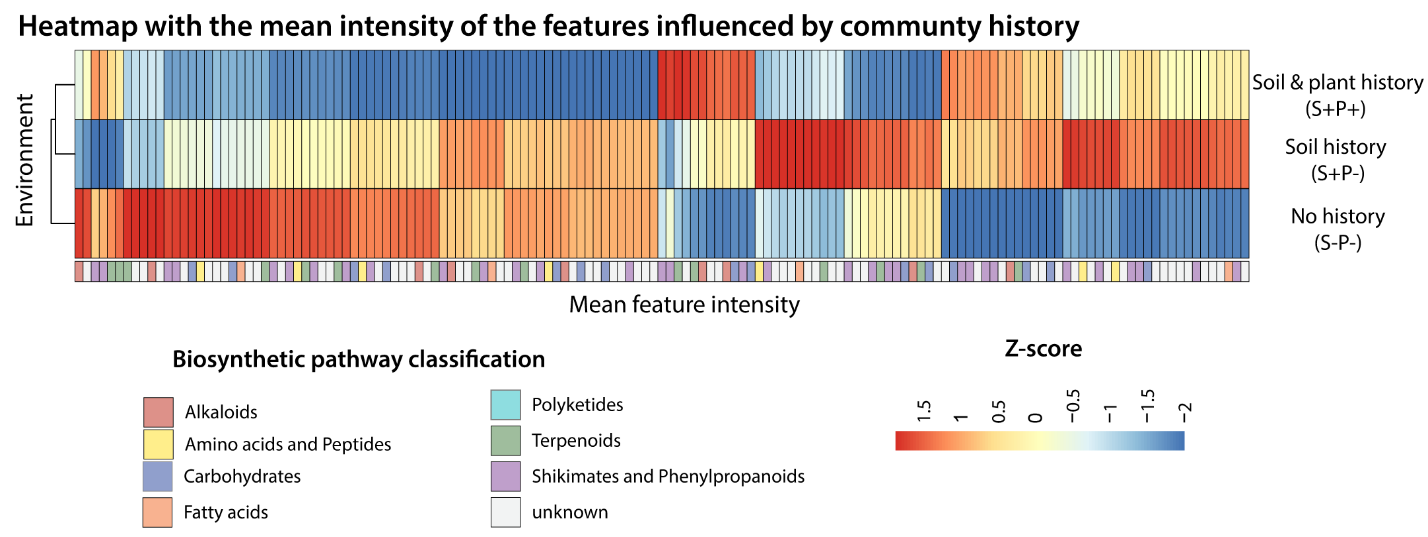

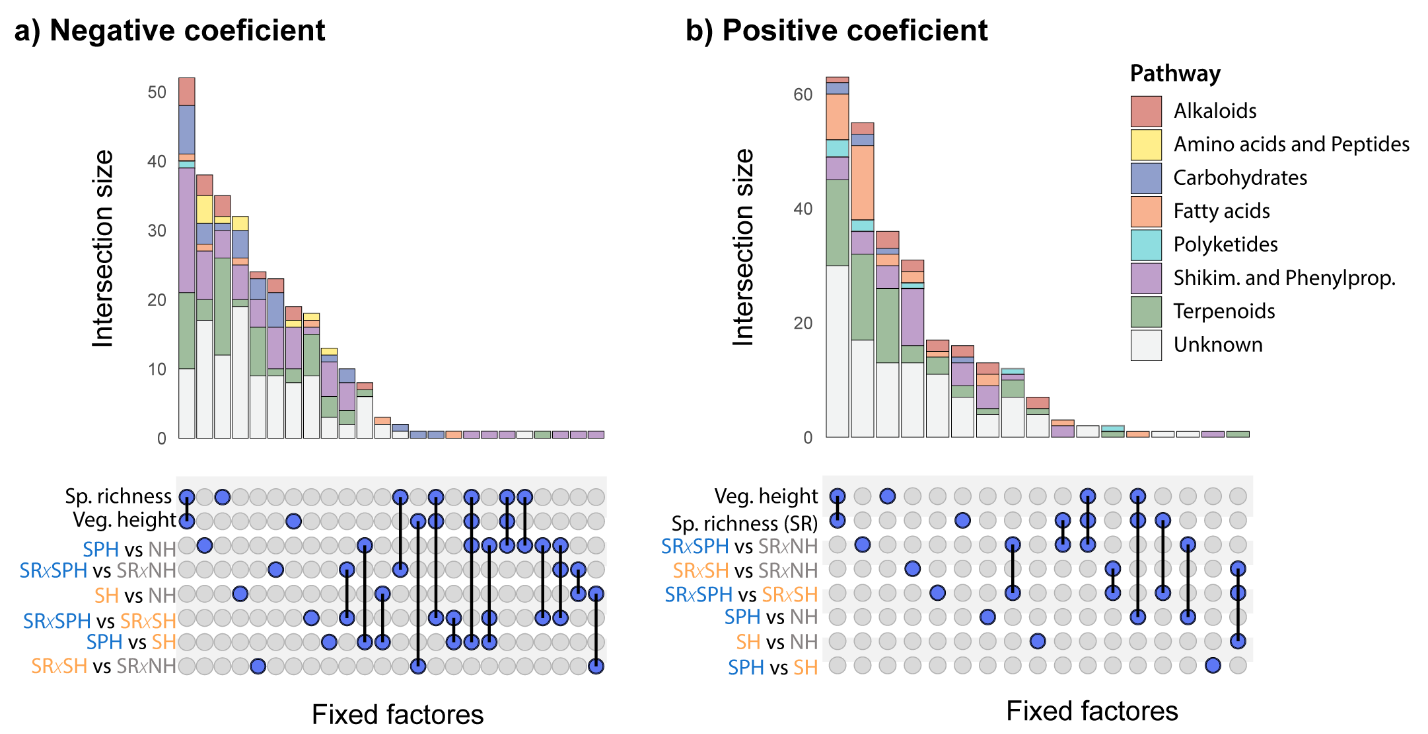

**Figure S3. Upset plot of the interactions of 634 features whose intensity were significantly influenced by vegetation height, species richness, community history or their interaction.**

a) Features that were negatively correlated with the treatment; b) Features that were positively correlated with the treatment. The blue bars (bottom left) indicate the number of metabolites whose intensity was significantly influenced by each fixed factor. The blue dots (connected with black lines) represent the intersections of metabolites by each factor, and the bars (top) indicate the frequency of these intersections. Features were putatively classified by biosynthetic pathway. Effect of the treatments to the feature intensity was tested with linear mixed regression model (intensity~ vegetation height + species richness * environment+ [1| block/plot]). The community history experiment compared the metabolomic profiles of selected plants grown in different environment treatments based on the δBEF experiment established in 2016(Vogel et al., 2019). No history (NH: –SH-PH): soil and plant layer removed, Soil history (SH: +SH-PH): only plant layer removed. In both treatments, new plot-specific plants species-mixtures were sowed. Soil-Plant history (SPH: +SH+PH): as control, same as core area stablished in 2002.

**Table S1. Selection *Experiment:* Wald-chi-squared analysis of variance (ANOVA) results for the linear mixed models of *naïve* and *selected* *Plantago lanceolata* phytometers across a diversity gradient based on leaf traits and leaf damage.**

The effects of vegetation height, plant species richness, selection history (*naïve* or *selected*) and legumes (presence or absence) on leaf traits and leaf damage were tested using mixed-effects models. Six models were run to disentangle the confounding effects: Model 1 examined species richness, selection treatment, and their interaction. Models 2 and 3 assessed legume presence, either before or after species richness. Models 4-6 tested vegetation height, including it as a covariate. All models used plot nested within block as random effects. The table shows Chi-square (X²) and p-values for fixed effects, with significant effects in bold (P< 0.05) and tendencies within brackets (P < 0.1). Data were transformed as needed to meet assumptions.

| **Trait** | **Explanatory variable** | **Model 1** | | **Model 2** | | **Model 3** | | **Model 4** | | **Model 5** | | **Model 6** | |  |
| --- | --- | --- | --- | --- | --- | --- | --- | --- | --- | --- | --- | --- | --- | --- |
|  |  | *y~ SR* S+  (1\| block/ plot)* | | *y~ SR*L* S+  (1\| block/ plot)* | | *y~ L*SR*S+  (1\| block/ plot)* | | *y~ VG+SR*S+  (1\| block/ plot)* | | *y~ VG+SR*L*S+  (1\| block/ plot)* | | *y~ VG+L*SR* S+  (1\| block/ plot)* | |  |
|  |  | **X2** | **P** | **X2** | **P** | **X2** | **P** | **X2** | **P** | **X2** | **P** | **X2** | **P** |  |
| **Morphological leaf traits** | | | | | | | | | | | | | | |
| Shoot biomass  g DW glmer  N= 169 | Veg. height | NA | NA | NA | NA | NA | NA | **5.22** | **0.022** | **5.22** | **0.022** | **5.22** | **0.022** | Shoot biomass increased with increasing species richness only in *selected* phytometers. |
|  | Species richness (SR) | 1.48 | 0.224 | 1.48 | 0.224 | 5.24 | **0.022** | 0.06 | 0.810 | 0.06 | 0.810 | [2.99] | *0.084* |  |
|  | Legume (L) | NA | NA | **7.23** | **0.007** | [3.47] | *0.062* | NA | NA | [3.47] | *0.063* | 0.54 | 0.464 |  |
|  | Selection (S) | 0.00 | 0.969 | 0.02 | 0.896 | 0.02 | 0.896 | 0.02 | 0.892 | 0.02 | 0.895 | 0.02 | 0.895 |  |
|  | SR x L | NA | NA | 0.00 | 1.000 | 0.00 | 1.000 | NA | NA | 1.00 | 0.318 | 1.00 | 0.318 |  |
|  | SR x S | **4.95** | **0.026** | **5.36** | **0.021** | [3.52] | *0.061* | **5.11** | **0.024** | **5.30** | **0.021** | 2.72 | *0.099* |  |
|  | L x S | NA | NA | **4.60** | **0.032** | **6.45** | **0.011** | NA | NA | [3.16] | *0.076* | **5.74** | **0.017** |  |
|  | SR x L x S | NA | NA | 0.01 | 0.919 | 0.01 | 0.919 | NA | NA | 0.02 | 0.885 | 0.01 | 0.907 |  |
|  | R^2^ | 0.04/0.38 | | 0.19/0.44 | | 0.19/0.44 | | 0.1/0.35 | | 0.16/0.39 | | 0.16/0.4 | |  |
| Leaf length  cm glmer  N= 169 | Veg. height | NA | NA | NA | NA | NA | NA | **29.02** | **0.000** | **29.02** | **<0.001** | **29.02** | **<0.001** |  |
|  | Species richness (SR) | [2.98] | *0.084* | [2.98] | *0.084* | 2.12 | 0.146 | 2.03 | 0.154 | 2.03 | 0.154 | 0.03 | 0.864 |  |
|  | Legume (L) | NA | NA | 2.68 | 0.102 | [3.54] | *0.060* | NA | NA | **6.45** | **0.011** | **8.45** | **0.004** |  |
|  | Selection (S) | 0.03 | 0.859 | 0.03 | 0.856 | 0.03 | 0.855 | 0.01 | 0.920 | 0.01 | 0.918 | 0.01 | 0.918 |  |
|  | SR x L | NA | NA | 0.18 | 0.671 | 0.18 | 0.671 | NA | NA | 0.00 | 0.994 | 0.00 | 0.994 |  |
|  | SR x S | 1.90 | 0.168 | 1.82 | 0.178 | 2.39 | 0.122 | 2.45 | 0.117 | 1.93 | 0.165 | 2.41 | 0.121 |  |
|  | L x S | NA | NA | **4.96** | **0.026** | **4.39** | **0.036** | NA | NA | 3.86 | *0.050* | 3.38 | *0.066* |  |
|  | SR x L x S | NA | NA | 1.18 | 0.278 | 1.19 | 0.275 | NA | NA | 0.79 | 0.375 | 0.79 | 0.375 |  |
|  | R^2^ |  | |  | |  | |  | |  | |  | |  |
| Leaf greenness  SPAD glmer | Veg. height | NA | NA | NA | NA | NA | NA | 0.17 | 0.684 | 0.17 | 0.684 | 0.17 | 0.684 | Plots with legumes had higher SPAD. |
|  | Species richness (SR) | 0.64 | 0.423 | 0.64 | 0.423 | [3.44] | *0.064* | 1.42 | 0.234 | 1.42 | 0.234 | 1.36 | 0.243 |  |
|  | Legume (L) | NA | NA | **9.01** | **0.003** | **6.21** | **0.013** | NA | NA | **9.11** | **0.003** | **9.16** | **0.002** |  |
|  | Selection (S) | 0.90 | 0.344 | 0.93 | 0.335 | 0.93 | 0.335 | 0.95 | 0.329 | 0.86 | 0.353 | 0.86 | 0.353 |  |
|  | SR x L | NA | NA | 0.94 | 0.333 | 0.94 | 0.333 | NA | NA | 1.57 | 0.210 | 1.57 | 0.210 |  |
|  | SR x S | 0.07 | 0.793 | 0.11 | 0.738 | 0.06 | 0.808 | 0.07 | 0.792 | 0.17 | 0.684 | 0.10 | 0.751 |  |
|  | L x S | NA | NA | 0.33 | 0.563 | 0.39 | 0.533 | NA | NA | 0.29 | 0.588 | 0.36 | 0.549 |  |
|  | SR x L x S | NA | NA | [3.30] | *0.069* | [3.30] | *0.069* | NA | NA | [3.04] | *0.081* | [3.04] | *0.081* |  |
|  | R^2^ | 0.02/NA | | 0.11/NA | | 0.11/NA | | 0.03/NA | | 0.12/NA | | 0.12/NA | |  |
| Flowering status  (yes/no) binomial/glmer | Veg. height | NA | NA | NA | NA | NA | NA | 2.59 | 0.107 | 2.59 | 0.107 | 2.59 | 0.107 |  |
|  | Species richness (SR) | 2.66 | 0.103 | 2.66 | 0.103 | 1.89 | 0.169 | 0.97 | 0.324 | 0.97 | 0.324 | 1.16 | 0.282 |  |
|  | Legume (L) | NA | NA | 1.73 | 0.189 | 2.49 | 0.114 | NA | NA | 0.93 | 0.336 | 0.74 | 0.389 |  |
|  | Selection (S) | 0.03 | 0.864 | 0.02 | 0.890 | 0.02 | 0.890 | 0.02 | 0.876 | 0.02 | 0.891 | 0.02 | 0.891 |  |
|  | SR x L | NA | NA | 1.32 | 0.251 | 1.32 | 0.251 | NA | NA | 1.22 | 0.270 | 1.22 | 0.270 |  |
|  | SR x S | [3.73] | *0.053* | [3.69] | *0.055* | [3.55] | *0.060* | [3.74] | *0.053* | [3.69] | *0.055* | [3.55] | *0.060* |  |
|  | L x S | NA | NA | 0.04 | 0.844 | 0.18 | 0.674 | NA | NA | 0.04 | 0.846 | 0.18 | 0.675 |  |
|  | SR x L x S | NA | NA | 0.33 | 0.566 | 0.33 | 0.566 | NA | NA | 0.33 | 0.566 | 0.33 | 0.566 |  |
|  | R^2^ | 0.27/NA | | 0.23/0.53 | | 0.23/0.53 | | 0.29/NA | | 0.23/0.53 | | 0.23/0.53 | |  |
| **Percentage of leaf damage** | | | | | | | | | | | | | | |
| Herbivore damage  % arcsin  N= 56 | Veg. height | NA | NA | NA | NA | NA | NA | 0.76 | 0.383 | 0.76 | 0.383 | 0.76 | 0.383 |  |
|  | Species richness (SR) | 1.10 | 0.294 | 1.10 | 0.294 | 1.05 | 0.305 | 0.38 | 0.537 | 0.38 | 0.537 | 0.13 | 0.714 |  |
|  | Legume (L) | NA | NA | 0.05 | 0.821 | 0.10 | 0.752 | NA | NA | 0.16 | 0.691 | 0.41 | 0.524 |  |
|  | Selection (S) | 0.28 | 0.594 | 0.26 | 0.610 | 0.26 | 0.610 | 0.30 | 0.583 | 0.26 | 0.609 | 0.26 | 0.609 |  |
|  | SR x L | NA | NA | 0.64 | 0.422 | 0.64 | 0.422 | NA | NA | 1.11 | 0.293 | 1.11 | 0.293 |  |
|  | SR x S | 0.04 | 0.835 | 0.09 | 0.760 | 0.02 | 0.886 | 0.04 | 0.839 | 0.14 | 0.711 | 0.04 | 0.840 |  |
|  | L x S | NA | NA | 0.81 | 0.368 | 0.88 | 0.348 | NA | NA | 0.88 | 0.348 | 0.98 | 0.323 |  |
|  | SR x L x S | NA | NA | 0.14 | 0.710 | 0.14 | 0.710 | NA | NA | 0.10 | 0.757 | 0.10 | 0.757 |  |
|  | R^2^ | 0.04/NA | | 0.09/NA | | 0.09/NA | | 0.04/NA | | 0.12/NA | | 0.12/NA | |  |
| Pathogen damage  % arcsin  N= 56 | Veg. height | NA | NA | NA | NA | NA | NA | **15.22** | **<0.001** | **15.22** | **<0.001** | **15.22** | **<0.001** | Decrease with increasing vegetation height |
|  | Species richness (SR) | 1.52 | 0.218 | 1.52 | 0.218 | 0.74 | 0.390 | 0.78 | 0.376 | 0.78 | 0.376 | 0.93 | 0.334 |  |
|  | Legume (L) | NA | NA | [3.00] | *0.083* | [3.78] | *0.052* | NA | NA | 0.15 | 0.698 | 0.00 | 0.986 |  |
|  | Selection (S) | 1.56 | 0.211 | 1.30 | 0.253 | 1.30 | 0.253 | 1.21 | 0.271 | 1.27 | 0.260 | 1.27 | 0.260 |  |
|  | SR x L | NA | NA | 1.16 | 0.282 | 1.16 | 0.282 | NA | NA | 0.19 | 0.663 | 0.19 | 0.663 |  |
|  | SR x S | 0.01 | 0.934 | 0.05 | 0.815 | 0.13 | 0.721 | 0.09 | 0.770 | 0.05 | 0.826 | 0.10 | 0.749 |  |
|  | L x S | NA | NA | 0.64 | 0.424 | 0.57 | 0.452 | NA | NA | 0.35 | 0.555 | 0.29 | 0.588 |  |
|  | SR x L x S | NA | NA | 2.02 | 0.155 | 2.02 | 0.155 | NA | NA | 1.43 | 0.232 | 1.43 | 0.232 |  |
|  | R^2^ | 0.14/0.71 | | 0.36/0.73 | | 0.36/0.73 | | 0.59/0.69 | | 0.61/0.72 | | 0.61/0.72 | |  |
| Total leaf damage  % arcsin  N= 56 | Veg. height | NA | NA | NA | NA | NA | NA | **14.83** | **<0.001** | **14.83** | **<0.001** | **14.83** | **<0.001** | Decrease with increasing vegetation height |
|  | Species richness (SR) | 1.16 | 0.281 | 1.16 | 0.281 | 0.46 | 0.500 | 2.26 | 0.133 | 2.26 | 0.133 | 1.96 | 0.162 |  |
|  | Legume (L) | NA | NA | **4.05** | **0.044** | **4.76** | **0.029** | NA | NA | 0.00 | 0.995 | 0.30 | 0.583 |  |
|  | Selection (S) | [2.77] | [0.096] | 2.25 | 0.133 | 2.25 | 0.133 | 2.19 | 0.139 | 2.19 | 0.139 | 2.19 | 0.139 |  |
|  | SR x L | NA | NA | 1.47 | 0.225 | 1.47 | 0.225 | NA | NA | 0.00 | 0.956 | 0.00 | 0.956 |  |
|  | SR x S | 0.03 | 0.865 | 0.14 | 0.707 | 0.20 | 0.654 | 0.16 | 0.686 | 0.16 | 0.690 | 0.20 | 0.658 |  |
|  | L x S | NA | NA | 0.25 | 0.614 | 0.19 | 0.659 | NA | NA | 0.09 | 0.767 | 0.05 | 0.823 |  |
|  | SR x L x S | NA | NA | 1.21 | 0.272 | 1.21 | 0.272 | NA | NA | 0.74 | 0.389 | 0.74 | 0.389 |  |
|  | R^2^ | 0.12/0.67 | | 0.56/NA | | 0.56/NA | | 0.59/0.65 | | 0.6/0.67 | | 0.6/0.67 | |  |

**Table S2** **Community History Experiment: Wald-chi-squared analysis of variance (ANOVA) results for the linear mixed models of selected Plantago lanceolata phytometers across a diversity gradient in different community history environments based on leaf traits and leaf damage**.

The effects of vegetation height, species richness, experimental environment (S+P+, S+P-, S-P-) and legumes (presence or absence) on leaf traits and leaf damage were tested using mixed-effects models. The community history experiment compared the targeted defense compounds of selected phytometers grown in different experimental environments based on the ΔBEF experiment established in 2016 (Vogel et al., 2019). Six models were run to disentangle the confounding effects: Model 1 examined species richness, environement treatment, and their interaction. Models 2 and 3 assessed legume presence, either before or after species richness. Models 4-6 tested vegetation height, including it as a covariate. All models used plot nested within block as random effects. The table shows N, R^2^, Chi-square (X²) and p-values for fixed effects, with significant effects in bold (P< 0.05) and tendencies within brackets (P < 0.1). Data were transformed as needed to meet assumptions.

| **Leaf trait** | **Explanatory variable** | **Model 1** | | **Model 2** | | **Model 3** | | **Model 4** | | **Model 5** | | **Model 6** | | **Pattern** |
| --- | --- | --- | --- | --- | --- | --- | --- | --- | --- | --- | --- | --- | --- | --- |
|  |  | *y~ SR* E+  (1\| block/ plot)* | | *y~ SR*L* E+  (1\| block/ plot)* | | *y~ L*SR*E+  (1\| block/ plot)* | | *y~ VG+SR*E+  (1\| block/ plot)* | | *y~ VG+SR*L* E+  (1\| block/ plot)* | | *y~ VG+L*SR* E+  (1\| block/ plot)* | |  |
|  |  | **X2** | **P** | **X2** | **P** | **X2** | **P** | **X2** | **P** | **X2** | **P** | **X2** | **P** |  |
| **Morphological leaf traits** | | | | | | | | | | | | | | |
| Shoot biomass  g DW glmer  N= 169 | Veg. height | NA | NA | NA | NA | NA | NA | 0.30 | 0.586 | 0.30 | 0.586 | 0.30 | 0.586 | Shoot biomass increased with increasing species richness in *selected* phytometers only when they grew in their environment of origin. Effects are stronger in plots without legumes |
|  | Species richness (SR) | 1.03 | 0.309 | 1.03 | 0.309 | 1.01 | 0.314 | 0.76 | 0.384 | 0.76 | 0.384 | 1.06 | 0.303 |  |
|  | Legume (L) | NA | NA | 2.25 | 0.133 | 2.27 | 0.13 | NA | NA | 2.31 | 0.128 | 2.01 | 0.156 |  |
|  | Environment (E) | 1.61 | 0.456 | 1.55 | 0.461 | 1.55 | 0.461 | 1.59 | 0.451 | 1.66 | 0.436 | 1.66 | 0.436 |  |
|  | SR x L | NA | NA | 0.23 | 0.610 | 0.23 | 0.610 | NA | NA | 0.29 | 0.592 | 0.29 | 0.592 |  |
|  | SR x E | **9.86** | **0.007** | **9.94** | **0.007** | **8.20** | **0.016** | **10.10** | **0.006** | **9.70** | **0.007** | **7.82** | **0.020** |  |
|  | L x E | NA | NA | 0.94 | 0.62 | 2.69 | 0.260 | NA | NA | 0.94 | 0.617 | 2.83 | 0.242 |  |
|  | SR x L x E | NA | NA | **15.52** | **< 0.001** | **15.52** | **< 0.001** | NA | NA | **15.51** | **< 0.001** | **15.51** | **< 0.001** |  |
|  | R^2^ | 0.05/NA | | 0.13/0.39 | | 0.13/0.38 | | 0.06/NA | | 0.4/0.83 | | 0.42/0.84 | |  |
| Leaf length  cm glmer  N= 169 | Veg. height | NA | NA | NA | NA | NA | NA | 0.01 | 0.905 | 0.01 | 0.905 | 0.01 | 0.905 | Increased with increasing species richness regardless of the environment treatment |
|  | Species richness (SR) | **4.71** | **0.030** | **4.71** | **0.030** | **4.88** | **0.027** | **5.16** | **0.023** | **5.16** | **0.023** | **5.77** | **0.016** |  |
|  | Legume (L) | NA | NA | 2.27 | 0.132 | 2.10 | 0.147 | NA | NA | [2.82] | *0.093* | 2.20 | 0.138 |  |
|  | Environment (E) | [5.31] | *0.070* | [4.98] | *0.083* | [4.98] | *0.083* | [5.2] | *0.074* | [4.85] | *0.089* | [4.85] | *0.089* |  |
|  | SR x L | NA | NA | 0.02 | 0.897 | 0.02 | 0.897 | NA | NA | 0.05 | 0.818 | 0.05 | 0.818 |  |
|  | SR x E | 0.73 | 0.695 | 0.72 | 0.699 | 0.50 | 0.777 | 0.59 | 0.745 | 0.50 | 0.779 | 0.33 | 0.850 |  |
|  | L x E | NA | NA | [4.66] | *0.097* | [4.87] | *0.088* | NA | NA | [5.27] | *0.072* | [5.45] | *0.066* |  |
|  | SR x L x E | NA | NA | 3.98 | 0.137 | 3.98 | 0.137 | NA | NA | 3.50 | 0.174 | 3.50 | 0.174 |  |
|  | R^2^ | 0.1/0.21 | | 0.14/0.23 | | 0.14/0.23 | | 0.1/0.22 | | 0.14/0.24 | | 0.14/0.24 | |  |
| Leaf greenness  SPAD glmer  N= 169 | Veg. height | NA | NA | NA | NA | NA | NA | **7.56** | **0.006** | **7.56** | **0.006** | **7.56** | **0.006** | Plots with legumes had higher SPAD, while SPAD decreased with increasing vegetation height in their surroundings |
|  | Species richness (SR) | 2.11 | 0.1463 | 2.11 | 0.1463 | **13.99** | **< 0.001** | [3.65] | *0.056* | [3.65] | *0.056* | **2.13** | **0.144** |  |
|  | Legume (L) | NA | NA | **6.53** | **0.011** | 2.53 | 0.112 | NA | NA | **13.68** | **< 0.001** | **15.19** | **< 0.001** |  |
|  | Environment (E) | 0.42 | 0.810 | 0.29 | 0.863 | 0.29 | 0.863 | 0.14 | 0.932 | 0.46 | 0.793 | 0.46 | 0.793 |  |
|  | SR x L | NA | NA | 1.90 | 0.168 | 1.90 | 0.168 | NA | NA | 2.37 | 0.124 | 2.37 | 0.124 |  |
|  | SR x E | 2-95 | 0.229 | 2.06 | 0.357 | 1.23 | 0.540 | 1.52 | 0.468 | 1.35 | 0.510 | 0.64 | 0.727 |  |
|  | L x E | NA | NA | 2.03 | 0.362 | 2.86 | 0.239 | NA | NA | 2.70 | 0.260 | 3.40 | 0.182 |  |
|  | SR x L x E | NA | NA | 2.17 | 0.338 | 2.17 | 0.338 | NA | NA | 3.39 | 0.183 | 3.39 | 0.183 |  |
|  | R^2^ |  | |  | |  | |  | |  | |  | |  |
| Flowering status  (yes/no) binomial/glmer  N= 169 | Veg. height | NA | NA | NA | NA | NA | NA | 0.11 | 0.735 | 0.11 | 0.735 | 0.11 | 0.735 | Shoot biomass increased with increasing species richness in *selected* phytometers only when they grew in their environment of origin. Effects are stronger in plots without legumes |
|  | Species richness (SR) | 0.38 | 0.539 | 0.38 | 0.539 | 0.18 | 0.674 | 0.81 | 0.368 | 0.81 | 0.368 | 1.12 | 0.289 |  |
|  | Legume (L) | NA | NA | [2.87] | *0.090* | [3.07] | *0.080* | NA | NA | **4.12** | **0.042** | [3.81] | *0.051* |  |
|  | Environment (E) | 2.46 | 0.292 | 2.44 | 0.295 | 2.44 | 0.295 | 2.03 | 0.362 | 1.65 | 0.438 | 1.65 | 0.438 |  |
|  | SR x L | NA | NA | 0.05 | 0.822 | 0.05 | 0.822 | NA | NA | 0.12 | 0.724 | 0.12 | 0.724 |  |
|  | SR x E | **18.12** | **< 0.001** | **18.65** | **< 0.001** | **15.40** | **< 0.001** | 18.00 | **< 0.001** | 18.31 | **< 0.001** | **15.14** | **0.001** |  |
|  | L x E | NA | NA | 3.60 | 0.166 | **6.84** | **0.033** | NA | NA | 4.17 | 0.124 | **7.35** | **0.025** |  |
|  | SR x L x E | NA | NA | [4.76] | *0.092* | [4.76] | *0.092* | NA | NA | [4.74] | *0.093* | [4.74] | *0.093* |  |
|  | R^2^ | 0.15/0.25 | | 0.21/0.28 | | 0.21/0.28 | | 0.15/0.25 | | 0.23/NA | | 0.23/NA | |  |
| **Percentage of leaf damage** | | | | | | | | | | | | | | |
| Herbivore damage  % arcsin  N= 80 | Veg. height | NA | NA | NA | NA | NA | NA | 1.06 | 0.304 | 1.06 | 0.304 | 1.06 | 0.304 |  |
|  | Species richness (SR) | 2.08 | 0.149 | 2.08 | 0.149 | 1.78 | 0.183 | 1.04 | 0.308 | 1.04 | 0.308 | 1.26 | 0.262 |  |
|  | Legume (L) | NA | NA | 0.41 | 0.523 | 0.72 | 0.397 | NA | NA | 0.44 | 0.509 | 0.22 | 0.640 |  |
|  | Environment (E) | 1.23 | 0.539 | 1.45 | 0.483 | 1.45 | 0.483 | 1.46 | 0.483 | 1.44 | 0.488 | 1.44 | 0.488 |  |
|  | SR x L | NA | NA | 0.55 | 0.457 | 0.55 | 0.457 | NA | NA | 0.58 | 0.446 | 0.58 | 0.446 |  |
|  | SR x E | 0.89 | 0.641 | 1.01 | 0.603 | 0.96 | 0.618 | 0.93 | 0.627 | 1.03 | 0.597 | 1.05 | 0.591 |  |
|  | L x E | NA | NA | 2.22 | 0.330 | 2.26 | 0.323 | NA | NA | 2.49 | 0.288 | 2.47 | 0.291 |  |
|  | SR x L x E | NA | NA | 2.37 | 0.306 | 2.37 | 0.306 | NA | NA | 2.54 | 0.281 | 2.54 | 0.281 |  |
|  | R^2^ | 0.06/NA | | 0.13/NA | | 0.13/NA | | 0.06/0.07 | | 0.13/NA | | 0.13/NA | |  |
| Pathogen damage  % arcsin  N= 80 | Veg. height | NA | NA | NA | NA | NA | NA | 1.08 | 0.298 | 1.08 | 0.298 | 1.08 | 0.298 | Decrease with increasing species richness only in phytometers that grew with soil history |
|  | Species richness (SR) | 0.49 | 0.482 | 0.49 | 0.482 | 0.09 | 0.760 | 0.01 | 0.925 | 0.01 | 0.925 | 0.01 | 0.934 |  |
|  | Legume (L) | NA | NA | [3.56] | *0.059* | **3.96** | **0.047** | NA | NA | [3.05] | *0.081* | [3.05] | *0.081* |  |
|  | Environment (E) | 1.79 | 0.408 | 1.71 | 0.425 | 1.71 | 0.425 | 2.09 | 0.351 | 1.82 | 0.403 | 1.82 | 0.403 |  |
|  | SR x L | NA | NA | 1.32 | 0.250 | 1.32 | 0.250 | NA | NA | 1.38 | 0.241 | 1.38 | 0.241 |  |
|  | SR x E | **8.25** | **0.016** | **8.71** | **0.013** | **6.49** | **0.039** | **9.36** | **0.009** | **9.47** | **0.009** | **7.30** | **0.026** |  |
|  | L x E | NA | NA | 1.25 | 0.536 | 3.47 | 0.176 | NA | NA | 1.23 | 0.541 | 3.40 | 0.183 |  |
|  | SR x L x E | NA | NA | 0.05 | 0.974 | 0.05 | 0.974 | NA | NA | 0.56 | 0.754 | 0.56 | 0.754 |  |
|  | R^2^ | 0.1/0.45 | | 0.35/NA | | 0.35/NA | | 0.19/0.35 | | 0.35/NA | | 0.35/NA | |  |
| Total leaf damage  % arcsin  N= 80 | Veg. height | NA | NA | NA | NA | NA | NA | **4.29** | **0.038** | **4.29** | **0.038** | **4.29** | **0.038** | Decreased with increasing vegetation height |
|  | Species richness (SR) | 0.51 | 0.474 | 0.51 | 0.474 | 0.03 | 0.863 | 0.01 | 0.931 | 0.01 | 0.931 | 0.04 | 0.849 |  |
|  | Legume (L) | NA | NA | [2.77] | *0.096* | [3.25] | *0.072* | NA | NA | 0.76 | 0.384 | 0.73 | 0.394 |  |
|  | Environment (E) | 2.97 | 0.226 | 2.76 | 0.252 | 2.76 | 0.252 | 2.77 | 0.251 | 2.63 | 0.268 | 2.63 | 0.268 |  |
|  | SR x L | NA | NA | 1.60 | 0.206 | 1.60 | 0.206 | NA | NA | 1.69 | 0.193 | 1.69 | 0.193 |  |
|  | SR x E | **6.19** | **0.045** | **6.56** | **0.038** | 4.24 | 0.120 | **7.77** | **0.021** | **7.91** | **0.019** | [5.46] | *0.065* |  |
|  | L x E | NA | NA | 2.23 | 0.327 | 4.56 | 0.102 | NA | NA | 1.43 | 0.488 | 3.88 | 0.144 |  |
|  | SR x L x E | NA | NA | 0.16 | 0.921 | 0.16 | 0.921 | NA | NA | 0.39 | 0.823 | 0.39 | 0.823 |  |
|  | R^2^ | 0.1/0.41 | | 0.32/NA | | 0.32/NA | | 0.27/NA | | 0.33/NA | | 0.33/NA | |  |

**Table S3. List of volatile organic compounds (VOC) identified in phytometers of *Plantago lanceolata* transplanted in the Jena Experiment**.

Headspace VOC collection of *P. lanceolata* was performed one year after transplantation using a push-pull system for two hours. Individual plants were enclosed with PET and tied at the bottom using cable binders and sponge to avoid damage. Ambient air entered the system after passing through an activated charcoal filter at a flow rate of 7 mL/min and was pumped out through a trap a rate of 4 mL/min. The trap contained 25 mg Porapak Q adsorbent in a Teflon tube that was inserted in the PET bag. Compounds are sorted by chemical class and retention time (RT in min). † symbol represents compounds identified by comparison to authentic standards, otherwise, they were identified by comparison of mass spectra and retention times to those in Willey or Nist Mass library.

| **Compound** | **Class** | **RT GC-FID** | **RT GC-MS** |
| --- | --- | --- | --- |
| pseudocumene | Aromatic | 8.02 | 7.76 |
| 1-methyl-3-Propylbenzene | Aromatic | 10.18 | 9.89 |
| 3-ethylbenzaldehyde | Aromatic | 12.77 |  |
| 4′-Ethylacetophenone | Aromatic | 15.37 | 15.51 |
| 2-hexanol | Fatty acid derivate | 4.40 | 4.11 |
| 4-methyloctane | Fatty acid derivate | 4.69 | 4.32 |
| *E*-2-hexanal † | Fatty acid derivate (GLV) | 5.23 | 5.05 |
| *Z*-3-hexenol | Fatty acid derivate (GLV) | 5.60 | 5.11 |
| 1-octen-3-ol † | Fatty acid derivate | 8.57 | 8.10 |
| 3-octanone | Fatty acid derivate | 8.82 | 8.27 |
| 3-octanol † | Fatty acid derivate | 8.91 | 8.50 |
| 1-octen-3-yl acetate | Fatty acid derivate | 11.73 | 11.46 |
| *E*-3-hexen-1-ol acetate † | Fatty acid derivate (GLV) | 9.15 | 8.79 |
| hexyl acetate | Fatty acid derivate (GLV) | 9.32 | 8.96 |
| *E*-2-hexenyl acetate | Fatty acid derivate (GLV) | 9.38 | 9.29 |
| tricyclene | Monoterpene | 7.04 | 6.66 |
| α-pinene † | Monoterpene | 7.3 | 6.88 |
| sabinene † | Monoterpene | 7.83 | 7.88 |
| β-pinene † | Monoterpene | 8.27 | 7.94 |
| *E*-β-ocimene † | Monoterpene | 10.08 | 9.84 |
| β -ocimene † | Monoterpene | 10.14 | 9.81 |
| DMNT † | Homoterpene | 11.78 | 11.55 |
| α-copaene † | Sesquiterpene | 17.76 | 17.7 |
| β-elemene † | Sesquiterpene | 18.19 | 18.05 |
| β-caryophyllene † | Sesquiterpene | 18.75 | 18.65 |
| α-bergamotene | Sesquiterpene | 19.04 | 18.96 |
| germacreneD | Sesquiterpene | 20.03 | 19.96 |
| unknown sesquiterpene | Sesquiterpene | 20.08 | 20.02 |
| unknown 1 | Other | 9.23 | 8.88 |
| unknown 2 | Other | 10.25 | 10.03 |
| unknown 3 | Other | 10.47 | 10.26 |

**Table S4. Selection Experiment: Wald-chi-squared analysis of variance (ANOVA) results for the linear mixed models of naïve and selected *Plantago lanceolata* phytometers across a diversity gradient based on** **volatile organic compound profiles*.***

The effects of vegetation height, plant species richness, selection history (*naïve* or *selected*) and legumes (presence or absence) on untargeted metabolome diversity were tested using mixed-effects models. Six models were run to disentangle the confounding effects: Model 1 examined species richness, selection treatment, and their interaction. Models 2 and 3 assessed legume presence, either before or after species richness. Models 4-6 tested vegetation height, including it as a covariate. All models used plot nested within block as random effects. The table shows Chi-square (X²) and p-values for fixed effects, with significant effects in bold (P< 0.05) and tendencies within brackets (P < 0.1). Data were transformed as needed to meet assumptions.

| **Trait** | **Explanatory factor** | **Model 1** | | **Model 2** | | **Model 3** | | **Model 4** | | **Model 5** | | **Model 6** | | **Pattern** |
| --- | --- | --- | --- | --- | --- | --- | --- | --- | --- | --- | --- | --- | --- | --- |
|  |  | y~ SR* S+  (1\| block/ plot) | | y~ SR*L* S+  (1\| block/ plot) | | y~ L*SR*S+  (1\| block/ plot) | | y~ VG+SR*S+  (1\| block/ plot) | | y~ VG+SR*L*S+  (1\| block/ plot) | | y~ VG+L*SR* S+  (1\| block/ plot) | |  |
|  |  | **X2** | **P** | **X2** | **P** | **X2** | **P** | **X2** | **P** | **X2** | **P** | **X2** | **P** |  |
| **Emission of volatile organic compounds** | | | | | | | | | | | | | | |
| Aromatic sqrt N= 60 | Veg. height | NA | NA | NA | NA | NA | NA | **4.57** | **0.033** | **4.57** | **0.033** | **4.57** | **0.033** | Decrease with increasing vegetation height. Effect stronger in communities with legumes |
|  | Species richness (SR) | **5.60** | **0.018** | **5.60** | **0.018** | **5.29** | **0.021** | 1.03 | 0.310 | 1.03 | 0.310 | 0.69 | 0.407 |  |
|  | Legume (L) | NA | NA | 0.01 | 0.916 | 0.32 | 0.572 | NA | NA | 1.56 | 0.211 | 1.91 | 0.167 |  |
|  | Selection (S) | 0.12 | 0.734 | 0.12 | 0.731 | 0.12 | 0.731 | 0.12 | 0.734 | 0.31 | 0.578 | 0.31 | 0.578 |  |
|  | SR x L | NA | NA | **6.74** | **0.009** | **6.74** | **0.009** | NA | NA | **5.00** | **0.025** | **5.00** | **0.025** |  |
|  | SR x S | 1.72 | 0.190 | 2.29 | 0.130 | 1.97 | 0.160 | 1.72 | 0.190 | 2.31 | 0.129 | 2.00 | 0.157 |  |
|  | L x S | NA | NA | 0.39 | 0.533 | 0.71 | 0.400 | NA | NA | 0.40 | 0.528 | 0.70 | 0.402 |  |
|  | SR x L x S | NA | NA | 0.21 | 0.646 | 0.21 | 0.646 | NA | NA | 0.19 | 0.662 | 0.19 | 0.662 |  |
|  | R2 | 0.48/NA | | 0.35/NA | | 0.35/NA | | 0.48/NA | | 0.35/NA | | 0.35/NA | |  |
| Green leaf volatiles log10 N= 60 | Veg. height | NA | NA | NA | NA | NA | NA | 1.16 | 0.282 | 1.16 | 0.282 | 1.16 | 0.282 |  |
|  | Species richness (SR) | 0.39 | 0.533 | 0.39 | 0.533 | 0.13 | 0.718 | 0.02 | 0.893 | 0.02 | 0.893 | 0.13 | 0.718 |  |
|  | Legume (L) | NA | NA | 1.92 | 0.165 | 2.18 | 0.140 | NA | NA | 1.15 | 0.284 | 1.04 | 0.309 |  |
|  | Selection (S) | 0.01 | 0.919 | 0.01 | 0.927 | 0.01 | 0.927 | 0.02 | 0.901 | 0.01 | 0.930 | 0.01 | 0.930 |  |
|  | SR x L | NA | NA | 0.45 | 0.503 | 0.45 | 0.503 | NA | NA | 0.65 | 0.419 | 0.65 | 0.419 |  |
|  | SR x S | 0.49 | 0.485 | 0.70 | 0.402 | 0.50 | 0.481 | 0.52 | 0.473 | 0.76 | 0.383 | 0.55 | 0.460 |  |
|  | L x S | NA | NA | 0.51 | 0.474 | 0.72 | 0.397 | NA | NA | 0.50 | 0.480 | 0.71 | 0.399 |  |
|  | SR x L x S | NA | NA | 0.16 | 0.693 | 0.16 | 0.693 | NA | NA | 0.14 | 0.710 | 0.14 | 0.710 |  |
|  | R2 | 0.01/NA | | 0.07/NA | | 0.07/NA | | 0.03/NA | | 0.07/NA | | 0.07/NA | |  |
| Monoterpene sqrt N= 60 | Veg. height | NA | NA | NA | NA | NA | NA | 0.36 | 0.551 | 0.36 | 0.551 | 1.18 | 0.278 |  |
|  | Species richness (SR) | 1.70 | 0.193 | 1.70 | 0.193 | 1.65 | 0.199 | [3.63] | *0.057* | [3.63] | *0.057* | 1.77 | 0.184 |  |
|  | Legume (L) | NA | NA | 0.43 | 0.514 | 0.47 | 0.491 | NA | NA | 0.15 | 0.697 | 1.54 | 0.215 |  |
|  | Selection (S) | 0.01 | 0.940 | 0.00 | 0.963 | 0.00 | 0.963 | 0.01 | 0.939 | 0.01 | 0.921 | 0.16 | 0.687 |  |
|  | SR x L | NA | NA | 0.04 | 0.837 | 0.04 | 0.837 | NA | NA | 0.23 | 0.633 | 0.02 | 0.887 |  |
|  | SR x S | 1.37 | 0.242 | 1.25 | 0.264 | 1.23 | 0.268 | 1.32 | 0.251 | 1.55 | 0.214 | 1.50 | 0.221 |  |
|  | L x S | NA | NA | 0.00 | 0.972 | 0.02 | 0.886 | NA | NA | 0.00 | 0.966 | 0.08 | 0.775 |  |
|  | SR x L x S | NA | NA | 1.32 | 0.251 | 1.32 | 0.251 | NA | NA | 1.67 | 0.196 | 0.40 | 0.527 |  |
|  | R2 | 0.07/NA | | 0.1/NA | | 0.1/NA | | 0.13/NA | | 0.18/NA | | 0.14/NA | |  |
| Sesquiterpene log10 N= 60 | Veg. height | NA | NA | NA | NA | NA | NA | **6.21** | **0.013** | **6.21** | **0.013** | **6.21** | **0.013** | Decrease with increasing species richness. Effect stronger in communities with legumes |
|  | Species richness (SR) | 0.02 | 0.884 | 0.02 | 0.884 | 1.18 | 0.277 | **6.93** | **0.008** | **6.93** | **0.008** | **5.98** | **0.014** |  |
|  | Legume (L) | NA | NA | **11.59** | **0.001** | **10.43** | **0.001** | NA | NA | **5.19** | **0.023** | **6.13** | **0.013** |  |
|  | Selection (S) | 0.27 | 0.606 | 0.05 | 0.821 | 0.05 | 0.821 | 0.17 | 0.678 | 0.10 | 0.756 | 0.10 | 0.756 |  |
|  | SR x L | NA | NA | 0.80 | 0.371 | 0.80 | 0.371 | NA | NA | 0.11 | 0.743 | 0.11 | 0.743 |  |
|  | SR x S | 0.27 | 0.605 | 0.00 | 0.992 | 0.02 | 0.896 | 0.06 | 0.804 | 0.03 | 0.858 | 0.00 | 0.970 |  |
|  | L x S | NA | NA | 0.88 | 0.347 | 0.87 | 0.352 | NA | NA | 0.90 | 0.342 | 0.93 | 0.334 |  |
|  | SR x L x S | NA | NA | 0.63 | 0.429 | 0.63 | 0.429 | NA | NA | 0.47 | 0.493 | 0.47 | 0.493 |  |
|  | R2 | 0.01/0.42 | | 0.36/NA | | 0.36/NA | | 0.4/NA | | 0.45/NA | | 0.45/NA | |  |
| others sqrt N= 60 | Veg. height | NA | NA | NA | NA | NA | NA | 0.07 | 0.794 | 0.07 | 0.794 | 0.07 | 0.794 | Decrease with increasing species richness. |
|  | Species richness (SR) | **5.45** | **0.020** | **5.45** | **0.020** | **5.78** | **0.016** | **7.36** | **0.007** | **7.36** | **0.007** | **7.60** | **0.006** |  |
|  | Legume (L) | NA | NA | 0.33 | 0.567 | 0.00 | 0.971 | NA | NA | 0.28 | 0.595 | 0.04 | 0.847 |  |
|  | Selection (S) | 0.15 | 0.696 | 0.15 | 0.701 | 0.15 | 0.701 | 0.15 | 0.695 | 0.16 | 0.686 | 0.16 | 0.686 |  |
|  | SR x L | NA | NA | 0.01 | 0.919 | 0.01 | 0.919 | NA | NA | 0.85 | 0.358 | 0.85 | 0.358 |  |
|  | SR x S | 0.09 | 0.770 | 0.08 | 0.774 | 0.18 | 0.670 | 0.06 | 0.802 | 0.13 | 0.718 | 0.25 | 0.615 |  |
|  | L x S | NA | NA | 0.75 | 0.386 | 0.65 | 0.420 | NA | NA | 0.72 | 0.396 | 0.60 | 0.439 |  |
|  | SR x L x S | NA | NA | 0.02 | 0.886 | 0.02 | 0.886 | NA | NA | 0.05 | 0.820 | 0.05 | 0.820 |  |
|  | R2 | 0.18/NA | | 0.2/NA | | 0.2/NA | | 0.22/NA | | 0.25/NA | | 0.25/NA | |  |
| Total emission log10 N= 60 | Veg. height | NA | NA | NA | NA | NA | NA | 1.31 | 0.253 | 1.31 | 0.253 | 1.31 | 0.253 |  |
|  | Species richness (SR) | [2.76] | *0.097* | [2.76] | *0.097* | 2.14 | 0.143 | 1.63 | 0.202 | 1.63 | 0.202 | 2.13 | 0.144 |  |
|  | Legume (L) | NA | NA | 1.00 | 0.318 | 1.61 | 0.204 | NA | NA | 0.96 | 0.328 | 0.45 | 0.502 |  |
|  | Selection (S) | 0.00 | 0.959 | 0.00 | 0.966 | 0.00 | 0.966 | 0.00 | 0.951 | 0.00 | 0.978 | 0.00 | 0.978 |  |
|  | SR x L | NA | NA | 0.17 | 0.679 | 0.17 | 0.679 | NA | NA | 0.45 | 0.504 | 0.45 | 0.504 |  |
|  | SR x S | 0.66 | 0.415 | 0.82 | 0.365 | 0.66 | 0.418 | 0.68 | 0.411 | 0.91 | 0.339 | 0.74 | 0.389 |  |
|  | L x S | NA | NA | 0.24 | 0.625 | 0.40 | 0.525 | NA | NA | 0.22 | 0.635 | 0.40 | 0.528 |  |
|  | SR x L x S | NA | NA | 0.45 | 0.501 | 0.45 | 0.501 | NA | NA | 0.41 | 0.520 | 0.41 | 0.520 |  |
|  | R2 | 0.06/NA | | 0.09/NA | | 0.09/NA | | 0.06/NA | | 0.09/NA | | 0.09/NA | |  |
| **Volatile compound diversity** | | | | | | | | | | | | | | |
| VOC Richness  Hill q0  Negative binomial N= 60 | Veg. height | NA | NA | NA | NA | NA | NA | [3.47] | *0.062* | [3.47] | *0.062* | 3.47 | *0.062* | Decrease with increasing species richness. Plots with legumes has less compounds |
|  | Species richness (SR) | 0.37 | 0.544 | 0.37 | 0.544 | 1.58 | 0.209 | **4.72** | **0.030** | **4.72** | **0.030** | **4.06** | **0.044** |  |
|  | Legume (L) | NA | NA | **6.51** | **0.011** | **5.30** | **0.021** | NA | NA | 1.64 | 0.201 | 2.30 | 0.129 |  |
|  | Selection (S) | [3.20] | *0.074* | [3.26] | *0.071* | [3.26] | *0.071* | [3.43] | *0.064* | [3.48] | *0.062* | [3.48] | *0.062* |  |
|  | SR x L | NA | NA | 0.01 | 0.913 | 0.01 | 0.913 | NA | NA | 1.04 | 0.307 | 1.04 | 0.307 |  |
|  | SR x S | 0.00 | 0.962 | 0.00 | 0.999 | 0.06 | 0.804 | 0.01 | 0.934 | 0.00 | 0.995 | 0.07 | 0.795 |  |
|  | L x S | NA | NA | 2.12 | 0.145 | 2.06 | 0.151 | NA | NA | 2.16 | 0.142 | 2.09 | 0.148 |  |
|  | SR x L x S | NA | NA | 0.74 | 0.390 | 0.74 | 0.390 | NA | NA | 0.85 | 0.358 | 0.85 | 0.358 |  |
|  | R2 | 0.25/1 | | 0.88/1 | | 0.88/1 | | 1/NA | | 1/NA | | 1/NA | |  |
| VOC Shannon  Hill q1 sqrt N= 60 | Veg. height | NA | NA | NA | NA | NA | NA | 0.42 | 0.515 | 0.42 | 0.515 | 0.42 | 0.515 | Decrease with increasing species richness. |
|  | Species richness (SR) | 0.98 | 0.322 | 0.98 | 0.322 | 2.64 | 0.105 | **3.97** | **0.048** | **3.97** | **0.048** | 2.46 | 0.117 |  |
|  | Legume (L) | NA | NA | **4.18** | **0.041** | 2.53 | 0.112 | NA | NA | 1.90 | 0.168 | 2.41 | 0.121 |  |
|  | Selection (S) | 0.69 | 0.405 | 0.78 | 0.376 | 0.78 | 0.376 | 0.81 | 0.368 | 0.81 | 0.368 | 0.81 | 0.368 |  |
|  | SR x L | NA | NA | 0.16 | 0.692 | 0.16 | 0.692 | NA | NA | 0.46 | 0.499 | 0.46 | 0.499 |  |
|  | SR x S | 0.09 | 0.761 | 0.09 | 0.762 | 0.20 | 0.652 | 0.09 | 0.761 | 0.06 | 0.806 | 0.15 | 0.700 |  |
|  | L x S | NA | NA | 1.48 | 0.224 | 1.37 | 0.242 | NA | NA | 1.52 | 0.218 | 1.43 | 0.232 |  |
|  | SR x L x S | NA | NA | 1.18 | 0.277 | 1.18 | 0.277 | NA | NA | 1.34 | 0.247 | 1.34 | 0.247 |  |
|  | R2 | 0.06/NA | | 0.21/0.3 | | 0.21/0.3 | | 0.16/NA | | 0.24/0.31 | | 0.24/0.31 | |  |
| VOC Simpson  Hill q2  log10 N= 60 | Veg. height | NA | NA | NA | NA | NA | NA | 0.85 | 0.357 | 0.85 | 0.357 | 0.85 | 0.357 |  |
|  | Species richness (SR) | 1.82 | 0.177 | 1.82 | 0.177 | 2.99 | *0.084* | [3.66] | *0.056* | [3.66] | *0.056* | 2.57 | 0.109 |  |
|  | Legume (L) | NA | NA | **4.88** | **0.027** | [3.72] | *0.054* | NA | NA | 2.20 | 0.138 | [3.29] | *0.070* |  |
|  | Selection (S) | 0.26 | 0.613 | 0.07 | 0.798 | 0.07 | 0.798 | 0.22 | 0.640 | 0.07 | 0.796 | 0.07 | 0.796 |  |
|  | SR x L | NA | NA | 0.74 | 0.389 | 0.74 | 0.389 | NA | NA | 0.81 | 0.367 | 0.81 | 0.367 |  |
|  | SR x S | 0.22 | 0.641 | 0.23 | 0.632 | 0.90 | 0.344 | 0.48 | 0.490 | 0.22 | 0.640 | 0.89 | 0.346 |  |
|  | L x S | NA | NA | [3.09] | *0.079* | 2.43 | 0.119 | NA | NA | [3.11] | *0.078* | 2.45 | 0.118 |  |
|  | SR x L x S | NA | NA | 0.39 | 0.530 | 0.39 | 0.530 | NA | NA | 0.42 | 0.516 | 0.42 | 0.516 |  |
|  | R2 | 0/NA | | 0.01/NA | | 0.01/NA | | 0/NA | | 0.01/NA | | 0.01/NA | |  |

**Table S5 Selection Experiment: Wald-chi-squared analysis of variance (ANOVA) results for the linear mixed models of *naïve* and *selected Plantago lanceolata* phytometers across a diversity gradient based on non-volatile** **untargeted metabolome diversity.**

The effects of vegetation height, plant species richness, selection history (*naïve* or *selected*) and legumes (presence or absence) on untargeted metabolome diversity were tested using mixed-effects models. Six models were run to disentangle the confounding effects: Model 1 examined species richness, selection treatment, and their interaction. Models 2 and 3 assessed legume presence, either before or after species richness. Models 4-6 tested vegetation height, including it as a covariate. All models used plot nested within block as random effects. The table shows Chi-square (X²) and p-values for fixed effects, with significant effects in bold (P< 0.05) and tendencies within brackets (P < 0.1). Data were transformed as needed to meet assumptions.

| **Variable** | **Factor** | **Model 1** | | **Model 2** | | **Model 3** | | **Model 4** | | **Model 5** | | **Model 6** | | **Pattern** |
| --- | --- | --- | --- | --- | --- | --- | --- | --- | --- | --- | --- | --- | --- | --- |
|  |  | y~ SR* S+  (1\| block/ plot) | | y~ SR*L* S+  (1\| block/ plot) | | y~ L*SR*S+  (1\| block/ plot) | | y~ VG+SR*S+  (1\| block/ plot) | | y~ VG+SR*L*S+  (1\| block/ plot) | | y~ VG+L*SR* S+  (1\| block/ plot) | |  |
|  |  | **X2** | **P** | **X2** | **P** | **X2** | **P** | **X2** | **P** | **X2** | **P** | **X2** | **P** |  |
| Metabolome richness Hill q0 negative binomial | Veg. height | NA | NA | NA | NA | NA | NA | 1.95 | 0.162 | 1.95 | 0.162 | 1.95 | 0.162 |  |
|  | Species richness (SR) | 0.19 | 0.662 | 0.19 | 0.662 | 0.24 | 0.623 | 2.62 | 0.106 | 2.62 | 0.106 | [2.96] | *0.085* |  |
|  | Legume (L) | NA | NA | 0.08 | 0.773 | 0.03 | 0.857 | NA | NA | 0.73 | 0.394 | 0.38 | 0.538 |  |
|  | Selection (S) | 0.00 | 1.000 | 0.00 | 0.978 | 0.00 | 0.978 | 0.00 | 0.957 | 0.00 | 0.994 | 0.00 | 0.994 |  |
|  | SR x L | NA | NA | [3.78] | *0.052* | [3.78] | *0.052* | NA | NA | 2.06 | 0.151 | 2.06 | 0.151 |  |
|  | SR x S | 1.39 | 0.238 | 1.38 | 0.240 | 1.27 | 0.260 | 1.43 | 0.231 | 1.46 | 0.226 | 1.36 | 0.243 |  |
|  | L x S | NA | NA | 0.03 | 0.868 | 0.14 | 0.710 | NA | NA | 0.02 | 0.901 | 0.12 | 0.733 |  |
|  | SR x L x S | NA | NA | 1.11 | 0.293 | 1.11 | 0.293 | NA | NA | 1.18 | 0.278 | 1.18 | 0.278 |  |
|  | R2 |  | |  | |  | |  | |  | |  | |  |
| Shannon metabolome diversity Hill q1 | Veg. height | NA | NA | NA | NA | NA | NA | 0.01 | 0.937 | 0.01 | 0.937 | 0.01 | 0.937 | Increase with increasing species richnes only in *selected* phytometers |
|  | Species richness (SR) | 2.00 | 0.158 | 2.00 | 0.158 | 2.13 | 0.145 | 2.53 | 0.112 | 2.53 | 0.112 | 2.26 | 0.132 |  |
|  | Legume (L) | NA | NA | 0.35 | 0.555 | 0.22 | 0.642 | NA | NA | 0.06 | 0.809 | 0.32 | 0.569 |  |
|  | Selection (S) | **4.04** | **0.044** | **4.25** | **0.039** | **4.25** | **0.039** | **4.14** | **0.042** | **4.23** | **0.040** | **4.23** | **0.040** |  |
|  | SR x L | NA | NA | 1.90 | 0.169 | 1.90 | 0.169 | NA | NA | 1.68 | 0.195 | 1.68 | 0.195 |  |
|  | SR x S | **6.35** | **0.012** | **6.71** | **0.010** | **5.80** | **0.016** | **6.39** | **0.011** | **6.70** | **0.010** | **5.78** | **0.016** |  |
|  | L x S | NA | NA | 0.71 | 0.401 | 1.62 | 0.204 | NA | NA | 0.71 | 0.400 | 1.62 | 0.203 |  |
|  | SR x L x S | NA | NA | 0.74 | 0.388 | 0.74 | 0.388 | NA | NA | 0.74 | 0.389 | 0.74 | 0.389 |  |
|  | R2 | 0.11/0.13 | | 0.15/0.16 | | 0.15/0.16 | | 0.12/NA | | 0.16/NA | | 0.16/NA | |  |
| Simpson metabolome diversity Hill q2 | Veg. height | NA | NA | NA | NA | NA | NA | 0.13 | 0.719 | 0.13 | 0.719 | 0.13 | 0.719 | Increase with increasing species richnes only in *selected* phytometers |
|  | Species richness (SR) | 0.34 | 0.559 | 0.34 | 0.559 | 0.39 | 0.530 | 0.79 | 0.373 | 0.79 | 0.373 | 0.62 | 0.431 |  |
|  | Legume (L) | NA | NA | 0.47 | 0.494 | 0.41 | 0.520 | NA | NA | 0.12 | 0.734 | 0.29 | 0.592 |  |
|  | Selection (S) | 1.57 | 0.210 | 1.72 | 0.189 | 1.72 | 0.189 | 1.64 | 0.200 | 1.72 | 0.190 | 1.72 | 0.190 |  |
|  | SR x L | NA | NA | 2.83 | *0.092* | 2.83 | *0.092* | NA | NA | 2.61 | 0.106 | 2.61 | 0.106 |  |
|  | SR x S | **7.90** | **0.005** | **8.38** | **0.004** | **7.47** | **0.006** | **7.94** | **0.005** | **8.38** | **0.004** | **7.47** | **0.006** |  |
|  | L x S | NA | NA | 0.48 | 0.489 | 1.39 | 0.239 | NA | NA | 0.48 | 0.490 | 1.39 | 0.239 |  |
|  | SR x L x S | NA | NA | 0.32 | 0.574 | 0.32 | 0.574 | NA | NA | 0.32 | 0.573 | 0.32 | 0.573 |  |
|  | R2 | 0.09/0.12 | | 0.13/NA | | 0.13/NA | | 0.1/NA | | 0.13/NA | | 0.13/NA | |  |

**Table S6** **Selection Experiment: Wald-chi-squared analysis of variance (ANOVA) results for the linear mixed models of *naïve* and *selected Plantago lanceolata* phytometers across a diversity gradient based on** **targeted defense metabolites.**

The effects of vegetation height, plant species richness, selection history (naïve or selected) and legumes (presence or absence) on untargeted metabolome diversity were tested using mixed-effects models. Six models were run to disentangle the confounding effects: Model 1 examined species richness, selection treatment, and their interaction. Models 2 and 3 assessed legume presence, either before or after species richness. Models 4-6 tested vegetation height, including it as a covariate. All models used plot nested within block as random effects. The table shows Chi-square (X²) and p-values for fixed effects, with significant effects in bold (P< 0.05) and tendencies within brackets (P < 0.1). Data were transformed as needed to meet assumptions.

| **Variable** | **Factor** | **Model 1** | | **Model 2** | | **Model 3** | | **Model 4** | | **Model 5** | | **Model 6** | | **Pattern** |
| --- | --- | --- | --- | --- | --- | --- | --- | --- | --- | --- | --- | --- | --- | --- |
|  |  | y~ SR* S+  (1\| block/ plot) | | y~ SR*L* S+  (1\| block/ plot) | | y~ L*SR*S+  (1\| block/ plot) | | y~ VG+SR*S+  (1\| block/ plot) | | y~ VG+SR*L*S+  (1\| block/ plot) | | y~ VG+L*SR* S+  (1\| block/ plot) | |  |
|  |  | **X2** | **P** | **X2** | **P** | **X2** | **P** | **X2** | **P** | **X2** | **P** | **X2** | **P** |  |
| **Defense hormones** | | | | | | | | | | | | | | |
| Jasmonic acid  (ng (gdw)-1) log10 N= 112 | Veg. height | NA | NA | NA | NA | NA | NA | 0.62 | 0.430 | 0.62 | 0.430 | 0.62 | 0.430 |  |
|  | Species richness (SR) | 0.00 | 0.979 | 0.00 | 0.979 | 0.06 | 0.811 | 0.56 | 0.454 | 0.56 | 0.454 | 0.53 | 0.466 |  |
|  | Legume (L) | NA | NA | 0.97 | 0.325 | 0.91 | 0.339 | NA | NA | 0.34 | 0.561 | 0.37 | 0.545 |  |
|  | Selection (S) | 1.79 | 0.181 | 1.65 | 0.199 | 1.65 | 0.199 | 1.62 | 0.203 | 1.57 | 0.210 | 1.57 | 0.210 |  |
|  | SR x L | NA | NA | 0.00 | 0.999 | 0.00 | 0.999 | NA | NA | 0.04 | 0.836 | 0.04 | 0.836 |  |
|  | SR x S | 0.02 | 0.881 | 0.03 | 0.857 | 0.05 | 0.819 | 0.03 | 0.865 | 0.03 | 0.852 | 0.05 | 0.819 |  |
|  | L x S | NA | NA | 0.07 | 0.786 | 0.05 | 0.817 | NA | NA | 0.06 | 0.812 | 0.04 | 0.843 |  |
|  | SR x L x S | NA | NA | 0.01 | 0.927 | 0.01 | 0.927 | NA | NA | 0.01 | 0.929 | 0.01 | 0.929 |  |
|  | R2 | 0.01/0.35 | | 0.04/0.35 | | 0.04/0.35 | | 0.09/NA | | 0.1/NA | | 0.1/NA | |  |
| Jasmonic acid isoleucine (JA-Ile) N= 112 | Veg. height | NA | NA | NA | NA | NA | NA | 0.22 | 0.642 | 0.22 | 0.642 | 0.22 | 0.642 |  |
|  | Species richness (SR) | 1.30 | 0.254 | 1.30 | 0.254 | 3.17 | *0.075* | 2.86 | *0.091* | 2.86 | *0.091* | 2.25 | 0.134 |  |
|  | Legume (L) | NA | NA | **4.58** | **0.032** | 2.71 | 0.100 | NA | NA | 2.96 | *0.086* | 3.56 | *0.059* |  |
|  | Selection (S) | [3.66] | *0.056* | [3.33] | *0.068* | [3.33] | *0.068* | [3.54] | *0.060* | [3.32] | *0.068* | 3.32 | *0.068* |  |
|  | SR x L | NA | NA | 0.13 | 0.718 | 0.13 | 0.718 | NA | NA | 0.19 | 0.663 | 0.19 | 0.663 |  |
|  | SR x S | 1.38 | 0.239 | 1.55 | 0.213 | 1.33 | 0.249 | 1.38 | 0.239 | 1.54 | 0.215 | 1.31 | 0.253 |  |
|  | L x S | NA | NA | 0.14 | 0.713 | 0.35 | 0.552 | NA | NA | 0.15 | 0.702 | 0.38 | 0.540 |  |
|  | SR x L x S | NA | NA | 0.86 | 0.355 | 0.86 | 0.355 | NA | NA | 0.87 | 0.351 | 0.87 | 0.351 |  |
|  | R2 | 0.07/0.26 | | 0.17/0.28 | | 0.17/0.28 | | 0.13/NA | | 0.2/NA | | 0.2/NA | |  |
| 12-hydroxy-jasmonic acid (OH-JA)  log10 N= 112 | Veg. height | NA | NA | NA | NA | NA | NA | [3.10] | *0.078* | [3.10] | *0.078* | [3.10] | *0.078* |  |
|  | Species richness (SR) | 1.35 | 0.246 | 1.35 | 0.246 | 1.01 | 0.316 | 0.11 | 0.738 | 0.11 | 0.738 | 0.11 | 0.746 |  |
|  | Legume (L) | NA | NA | 0.32 | 0.569 | 0.66 | 0.415 | NA | NA | 0.02 | 0.886 | 0.03 | 0.869 |  |
|  | Selection (S) | 0.15 | 0.695 | 0.17 | 0.676 | 0.17 | 0.676 | 0.17 | 0.685 | 0.16 | 0.689 | 0.16 | 0.689 |  |
|  | SR x L | NA | NA | **6.76** | **0.009** | **6.76** | **0.009** | NA | NA | **6.44** | **0.011** | **6.44** | **0.011** |  |
|  | SR x S | 0.31 | 0.575 | 0.35 | 0.556 | 0.75 | 0.387 | 0.34 | 0.559 | 0.41 | 0.521 | 0.83 | 0.361 |  |
|  | L x S | NA | NA | 2.06 | 0.151 | 1.66 | 0.198 | NA | NA | 2.02 | 0.155 | 1.60 | 0.206 |  |
|  | SR x L x S | NA | NA | 1.21 | 0.270 | 1.21 | 0.270 | NA | NA | 1.11 | 0.292 | 1.11 | 0.292 |  |
|  | R2 | 0.04/NA | | 0.17/NA | | 0.17/NA | | 0.07/0.19 | | 0.18/NA | | 0.18/NA | |  |
| 12-hydroxy-jasmonoyl-isoleucine  (12OH-JA-Ile) glmer N= 112 | Veg. height | NA | NA | NA | NA | NA | NA | 0.09 | 0.770 | 0.09 | 0.770 | 0.09 | 0.770 |  |
|  | Species richness (SR) | 1.86 | 0.173 | 1.86 | 0.173 | 2.89 | *0.089* | 1.99 | 0.158 | 1.99 | 0.158 | 0.99 | 0.320 |  |
|  | Legume (L) | NA | NA | 1.68 | 0.195 | 0.64 | 0.422 | NA | NA | 1.51 | 0.219 | 2.52 | 0.112 |  |
|  | Selection (S) | 1.11 | 0.292 | 1.01 | 0.316 | 1.01 | 0.316 | 1.09 | 0.296 | 1.00 | 0.316 | 1.00 | 0.316 |  |
|  | SR x L | NA | NA | 0.49 | 0.482 | 0.49 | 0.482 | NA | NA | 0.45 | 0.501 | 0.45 | 0.501 |  |
|  | SR x S | 0.38 | 0.536 | 0.43 | 0.514 | 0.48 | 0.490 | 0.38 | 0.539 | 0.43 | 0.512 | 0.48 | 0.488 |  |
|  | L x S | NA | NA | 0.07 | 0.797 | 0.02 | 0.899 | NA | NA | 0.07 | 0.796 | 0.02 | 0.899 |  |
|  | SR x L x S | NA | NA | [3.57] | *0.059* | [3.57] | *0.059* | NA | NA | [3.56] | *0.059* | [3.56] | *0.059* |  |
|  | R2 | 0.11/NA | | 0.21/NA | | 0.21/NA | | 0.11/NA | | 0.21/NA | | 0.21/NA | |  |
| 12-carboxy-jasmonoyl-L-isoleucine (COOH-JA-Ile) glmer N= 112 | Veg. height | NA | NA | NA | NA | NA | NA | 0.00 | 0.951 | 0.00 | 0.951 | 0.00 | 0.951 |  |
|  | Species richness (SR) | 0.68 | 0.410 | 0.68 | 0.410 | 0.70 | 0.401 | 0.86 | 0.353 | 0.86 | 0.353 | 0.85 | 0.355 |  |
|  | Legume (L) | NA | NA | 0.04 | 0.840 | 0.01 | 0.909 | NA | NA | 0.00 | 0.963 | 0.01 | 0.922 |  |
|  | Selection (S) | 0.17 | 0.682 | 0.16 | 0.692 | 0.16 | 0.692 | 0.14 | 0.709 | 0.14 | 0.708 | 0.14 | 0.708 |  |
|  | SR x L | NA | NA | 0.35 | 0.555 | 0.35 | 0.555 | NA | NA | 0.45 | 0.504 | 0.45 | 0.504 |  |
|  | SR x S | 0.84 | 0.358 | 0.70 | 0.403 | **6.25** | **0.012** | 0.89 | 0.345 | 0.75 | 0.388 | **6.76** | **0.009** |  |
|  | L x S | NA | NA | **20.72** | **0.000** | **15.17** | **0.000** | NA | NA | **21.26** | **0.000** | **15.25** | **0.000** |  |
|  | SR x L x S | NA | NA | 2.30 | 0.129 | 2.30 | 0.129 | NA | NA | 2.22 | 0.136 | 2.39 | 0.122 |  |
|  | R2 | 0.31/0.99 | | 0.74/1 | | 0.74/1 | | 0.97/NA | | 0.77/1 | | 0.79/1 | |  |
| Total jasmonates N= 112 | Veg. height | NA | NA | NA | NA | NA | NA | 0.86 | 0.355 | 0.86 | 0.355 | 0.86 | 0.355 |  |
|  | Species richness (SR) | 1.04 | 0.307 | 1.04 | 0.307 | 1.10 | 0.293 | 0.27 | 0.604 | 0.27 | 0.604 | 0.00 | 1.000 |  |
|  | Legume (L) | NA | NA | 0.06 | 0.805 | 0.00 | 0.985 | NA | NA | 0.22 | 0.642 | 0.55 | 0.460 |  |
|  | Selection (S) | 1.59 | 0.207 | 1.57 | 0.211 | 1.57 | 0.211 | 1.61 | 0.205 | 1.84 | 0.175 | 1.84 | 0.175 |  |
|  | SR x L | NA | NA | [2.86] | *0.091* | [2.86] | *0.091* | NA | NA | 2.39 | 0.123 | 2.39 | 0.123 |  |
|  | SR x S | 0.31 | 0.578 | 0.34 | 0.558 | 0.59 | 0.441 | 0.31 | 0.575 | 0.35 | 0.554 | 0.60 | 0.437 |  |
|  | L x S | NA | NA | 1.00 | 0.318 | 0.75 | 0.388 | NA | NA | 1.00 | 0.317 | 0.75 | 0.388 |  |
|  | SR x L x S | NA | NA | 1.02 | 0.314 | 1.02 | 0.314 | NA | NA | 1.00 | 0.316 | 1.00 | 0.316 |  |
|  | R2 | 0.05/NA | | 0.13/NA | | 0.13/NA | | 0.04/0.24 | | 0.14/NA | | 0.14/NA | |  |
| Abscisic acid log10 N =112 | Veg. height | NA | NA | NA | NA | NA | NA | **9.23** | **0.002** | **9.23** | **0.002** | **9.23** | **0.002** |  |
|  | Species richness (SR) | **3.44** | ***0.048*** | **3.44** | ***0.048*** | [3.11] | *0.078* | 0.12 | 0.733 | 0.12 | 0.733 | 0.05 | 0.825 |  |
|  | Legume (L) | NA | NA | 0.03 | 0.873 | 0.35 | 0.555 | NA | NA | 2.51 | 0.113 | 2.58 | 0.108 |  |
|  | Selection (S) | 0.03 | 0.864 | 0.03 | 0.858 | 0.03 | 0.858 | 0.11 | 0.736 | 0.07 | 0.790 | 0.07 | 0.790 |  |
|  | SR x L | NA | NA | [2.87] | *0.090* | 2.87 | *0.090* | NA | NA | 1.34 | 0.248 | 1.34 | 0.248 |  |
|  | SR x S | 1.68 | 0.195 | 1.70 | 0.192 | 1.22 | 0.269 | 1.56 | 0.211 | 1.88 | 0.171 | 1.38 | 0.240 |  |
|  | L x S | NA | NA | 0.85 | 0.357 | 1.33 | 0.249 | NA | NA | 0.77 | 0.380 | 1.26 | 0.261 |  |
|  | SR x L x S | NA | NA | 1.30 | 0.254 | 1.30 | 0.254 | NA | NA | 1.30 | 0.254 | 1.30 | 0.254 |  |
|  | R2 | 0.11/NA | | 0.15/0.25 | | 0.15/0.25 | | 0.29/NA | | 0.3/NA | | 0.3/NA | |  |
| Salicylic acid N=112 | Veg. height | NA | NA | NA | NA | NA | NA | **17.33** | **0.000** | **17.33** | **0.000** | **17.33** | **0.000** | Decreased with increasing diversity. |
|  | Species richness (SR) | **30.17** | **0.000** | **30.17** | **0.000** | **31.03** | **0.000** | **13.10** | **0.000** | **13.10** | **0.000** | **11.91** | **0.001** |  |
|  | Legume (L) | NA | NA | 0.87 | 0.352 | 0.00 | 0.953 | NA | NA | 2.58 | 0.108 | 3.77 | *0.052* |  |
|  | Selection (S) | 1.55 | 0.213 | 1.48 | 0.224 | 1.48 | 0.224 | 1.49 | 0.222 | 1.21 | 0.272 | 1.21 | 0.272 |  |
|  | SR x L | NA | NA | 0.47 | 0.492 | 0.47 | 0.492 | NA | NA | 1.27 | 0.260 | 1.27 | 0.260 |  |
|  | SR x S | [3.50] | *0.061* | [3.55] | *0.060* | [3.45] | *0.063* | [3.38] | *0.066* | [2.94] | *0.087* | [2.80] | *0.094* |  |
|  | L x S | NA | NA | 0.18 | 0.673 | 0.28 | 0.597 | NA | NA | 0.11 | 0.744 | 0.24 | 0.624 |  |
|  | SR x L x S | NA | NA | 0.18 | 0.672 | 0.18 | 0.672 | NA | NA | 0.03 | 0.867 | 0.03 | 0.867 |  |
|  | R2 | 0.03/NA | | 0.04/NA | | 0.04/NA | | 0.03/NA | | 0.03/NA | | 0.03/NA | |  |
| **Iridoid glycosides** | | | | | | | | | | | | | | |
| Aucubin N=107 | Veg. height | NA | NA | NA | NA | NA | NA | **8.02** | **0.005** | **8.02** | **0.005** | **8.02** | **0.005** | Considering the veg. height, concertation decreased with increasing SR. Veg. height increased concentration |
|  | Species richness (SR) | 0.16 | 0.693 | 0.16 | 0.693 | 0.37 | 0.543 | **9.45** | **0.002** | **9.45** | **0.002** | **10.36** | **0.001** |  |
|  | Legume (L) | NA | NA | 1.29 | 0.257 | 1.07 | 0.301 | NA | NA | 1.05 | 0.305 | 0.14 | 0.707 |  |
|  | Selection (S) | 0.79 | 0.374 | 0.94 | 0.333 | 0.94 | 0.333 | 1.18 | 0.277 | 1.12 | 0.291 | 1.12 | 0.291 |  |
|  | SR x L | NA | NA | 2.52 | 0.112 | 2.52 | 0.112 | NA | NA | 0.73 | 0.394 | 0.73 | 0.394 |  |
|  | SR x S | 0.99 | 0.319 | 0.91 | 0.341 | 0.83 | 0.361 | 1.09 | 0.297 | 0.97 | 0.325 | 0.88 | 0.348 |  |
|  | L x S | NA | NA | 0.02 | 0.885 | 0.09 | 0.761 | NA | NA | 0.03 | 0.861 | 0.12 | 0.730 |  |
|  | SR x L x S | NA | NA | [3.34] | *0.068* | [3.34] | *0.068* | NA | NA | [3.48] | *0.062* | [3.48] | *0.062* |  |
|  | R2 | 0.02/0.13 | | 0.09/0.15 | | 0.09/0.15 | | 0.19/NA | | 0.23/NA | | 0.23/NA | |  |
| Catalpol N=107 | Veg. height | NA | NA | NA | NA | NA | NA | 2.60 | 0.107 | 2.60 | 0.107 | 2.60 | 0.107 |  |
|  | Species richness (SR) | 0.31 | 0.579 | 0.31 | 0.579 | 0.34 | 0.560 | 0.12 | 0.726 | 0.12 | 0.726 | 0.41 | 0.523 |  |
|  | Legume (L) | NA | NA | 0.03 | 0.855 | 0.00 | 0.964 | NA | NA | 2.09 | 0.149 | 1.80 | 0.179 |  |
|  | Selection (S) | 0.00 | 0.993 | 0.00 | 0.996 | 0.00 | 0.996 | 0.01 | 0.935 | 0.00 | 1.000 | 0.00 | 1.000 |  |
|  | SR x L | NA | NA | [2.91] | *0.088* | [2.91] | *0.088* | NA | NA | [2.94] | *0.087* | [2.94] | *0.087* |  |
|  | SR x S | 0.97 | 0.326 | 0.84 | 0.361 | 0.60 | 0.440 | 0.96 | 0.327 | 0.73 | 0.392 | 0.54 | 0.462 |  |
|  | L x S | NA | NA | 0.60 | 0.440 | 0.84 | 0.361 | NA | NA | 0.39 | 0.530 | 0.59 | 0.443 |  |
|  | SR x L x S | NA | NA | 0.44 | 0.506 | 0.44 | 0.506 | NA | NA | 0.28 | 0.595 | 0.28 | 0.595 |  |
|  | R2 | 0.02/NA | | 0.07/NA | | 0.07/NA | | 0.05/NA | | 0.11/NA | | 0.11/NA | |  |
| **Phenylpropanoid glycosides** | | | | | | | | | | | | | | |
| Verbascoside N=107 | Veg. height | NA | NA | NA | NA | NA | NA | 0.15 | 0.701 | 0.15 | 0.701 | 0.15 | 0.701 | Decreased as SR increased. Naïve phytometers in communities without legumes did not vary across the diversity gradient |
|  | Species richness (SR) | 2.38 | 0.123 | 2.38 | 0.123 | 2.21 | 0.137 | **4.65** | **0.031** | **4.65** | **0.031** | **3.88** | **0.049** |  |
|  | Legume (L) | NA | NA | 0.75 | 0.386 | 0.92 | 0.338 | NA | NA | 0.01 | 0.925 | 0.77 | 0.379 |  |
|  | Selection (S) | 1.19 | 0.276 | 1.29 | 0.256 | 1.29 | 0.256 | 1.34 | 0.246 | 1.35 | 0.244 | 1.35 | 0.244 |  |
|  | SR x L | NA | NA | **4.92** | **0.027** | **4.92** | **0.027** | NA | NA | **4.39** | **0.036** | **4.39** | **0.036** |  |
|  | SR x S | 0.14 | 0.709 | 0.14 | 0.706 | 0.15 | 0.698 | 0.15 | 0.699 | 0.14 | 0.712 | 0.14 | 0.704 |  |
|  | L x S | NA | NA | 0.01 | 0.925 | 0.00 | 0.978 | NA | NA | 0.01 | 0.926 | 0.00 | 0.978 |  |
|  | SR x L x S | NA | NA | **4.49** | **0.034** | **4.49** | **0.034** | NA | NA | **4.41** | **0.036** | **4.41** | **0.036** |  |
|  | R2 | 0.1/NA | | 0.25/0.34 | | 0.25/0.34 | | 0.17/NA | | 0.28/NA | | 0.28/NA | |  |
| Plantamajoside N=112 | Veg. height | NA | NA | NA | NA | NA | NA | 2.35 | 0.125 | 2.35 | 0.125 | 2.35 | 0.125 | Decreased as SR increased. |
|  | Species richness (SR) | **7.14** | **0.008** | **7.14** | **0.008** | **7.08** | **0.008** | **4.96** | **0.026** | **4.96** | **0.026** | **5.67** | **0.017** |  |
|  | Legume (L) | NA | NA | 0.27 | 0.604 | 0.33 | 0.568 | NA | NA | 0.73 | 0.394 | 0.02 | 0.892 |  |
|  | Selection (S) | 0.24 | 0.625 | 0.28 | 0.599 | 0.28 | 0.599 | 0.22 | 0.640 | 0.26 | 0.609 | 0.26 | 0.609 |  |
|  | SR x L | NA | NA | 1.36 | 0.244 | 1.36 | 0.244 | NA | NA | 0.94 | 0.333 | 0.94 | 0.333 |  |
|  | SR x S | 0.09 | 0.761 | 0.13 | 0.723 | 0.32 | 0.573 | 0.09 | 0.764 | 0.13 | 0.720 | 0.33 | 0.568 |  |
|  | L x S | NA | NA | 1.18 | 0.277 | 0.99 | 0.320 | NA | NA | 1.21 | 0.271 | 1.01 | 0.314 |  |
|  | SR x L x S | NA | NA | 3.11 | *0.078* | 3.11 | *0.078* | NA | NA | [3.08] | *0.079* | [3.08] | *0.079* |  |
|  | R2 | 0.12/NA | | 0.2/NA | | 0.2/NA | | 0.13/NA | | 0.2/NA | | 0.2/NA | |  |

**Table S7 Community History Experiment: Wald-chi-squared analysis of variance (ANOVA) results for the linear mixed models of selected *Plantago lanceolata* phytometers across a diversity gradient in different community history environments based on** **volatile organic compounds profiles.**

The effects of vegetation height, species richness, experimental environment (*S+P+*, *S+P-*, *S-P-*) and legumes (presence or absence) on volatile organic compounds diversity were tested using mixed-effects models. *Community History Experiment* compared the metabolomic profiles of *selected* phytometers grown in different environment treatments based on the ΔBEF experiment established in 2016 (Vogel et al., 2019). Six models were run to disentangle the confounding effects: Model 1 examined species richness, environment treatment, and their interaction. Models 2 and 3 assessed legume presence, either before or after species richness. Models 4-6 tested vegetation height, including it as a covariate. All models used plot nested within block as random effects. The table shows Chi-square (X²) and p-values for fixed effects, with significant effects in bold (P< 0.05) and tendencies within brackets (P < 0.1). Data were transformed as needed to meet assumptions. N= 86

| **Variable** | **Factor** | **Model 1** | | **Model 2** | | **Model 3** | | **Model 4** | | **Model 5** | | **Model 6** | | **Pattern** |
| --- | --- | --- | --- | --- | --- | --- | --- | --- | --- | --- | --- | --- | --- | --- |
|  |  | y~ SR* E+  (1\| block/ plot) | | y~ SR*L* E+  (1\| block/ plot) | | y~ L*SR*E+  (1\| block/ plot) | | y~ VG+SR*E+  (1\| block/ plot) | | y~ VG+SR*L*E+  (1\| block/ plot) | | y~ VG+L*SR* E+  (1\| block/ plot) | |  |
|  |  | **X2** | **P** | **X2** | **P** | **X2** | **P** | **X2** | **P** | **X2** | **P** | **X2** | **P** |  |
| **Emission of volatile organic compounds** | | | | | | | | | | | | | | |
| aromatic | Veg. height | NA | NA | NA | NA | NA | NA | 0.03 | 0.852 | 0.03 | 0.852 | 0.03 | 0.852 | Decreased with increasing species richness |
|  | Species richness (SR) | 3.07 | *0.080* | 3.07 | *0.080* | 3.26 | *0.071* | **4.68** | **0.031** | **4.68** | **0.031** | **4.52** | **0.034** |  |
|  | Legume (L) | NA | NA | 0.24 | 0.628 | 0.05 | 0.817 | NA | NA | 0.02 | 0.885 | 0.18 | 0.670 |  |
|  | Environment (E) | 2.38 | 0.304 | 2.40 | 0.301 | 2.40 | 0.301 | 3.53 | 0.171 | 3.69 | 0.158 | 3.69 | 0.158 |  |
|  | SR x L | NA | NA | 1.36 | 0.244 | 1.36 | 0.244 | NA | NA | 1.39 | 0.239 | 1.39 | 0.239 |  |
|  | SR x E | 4.27 | 0.118 | 4.64 | *0.098* | 2.61 | 0.271 | 3.56 | 0.168 | 3.94 | 0.140 | 1.92 | 0.383 |  |
|  | L x E | NA | NA | 3.94 | 0.140 | 5.97 | *0.051* | NA | NA | 4.12 | 0.128 | **6.14** | **0.046** |  |
|  | SR x L x E | NA | NA | 2.78 | 0.249 | 2.78 | 0.249 | NA | NA | 3.08 | 0.214 | 3.08 | 0.214 |  |
|  | R2 | 0.16/0.29 | | 0.25/NA | | 0.25/NA | | 0.19/0.37 | | 0.24/0.38 | | 0.24/0.38 | |  |
| GLV | Veg. height | NA | NA | NA | NA | NA | NA | 1.04 | 0.308 | 1.04 | 0.308 | 1.04 | 0.308 |  |
|  | Species richness (SR) | 0.92 | 0.337 | 0.92 | 0.337 | 0.26 | 0.610 | 0.11 | 0.744 | 0.11 | 0.744 | 0.47 | 0.493 |  |
|  | Legume (L) | NA | NA | **4.12** | **0.042** | **4.78** | **0.029** | NA | NA | **4.11** | **0.043** | 3.75 | *0.053* |  |
|  | Environment (E) | 2.37 | 0.306 | 2.99 | 0.224 | 2.99 | 0.224 | 3.00 | 0.223 | 2.79 | 0.248 | 2.79 | 0.248 |  |
|  | SR x L | NA | NA | 3.51 | *0.061* | 3.51 | *0.061* | NA | NA | 3.72 | *0.054* | 3.72 | *0.054* |  |
|  | SR x E | 0.49 | 0.782 | 1.09 | 0.579 | 0.90 | 0.638 | 0.62 | 0.734 | 1.02 | 0.601 | 0.82 | 0.663 |  |
|  | L x E | NA | NA | 0.03 | 0.984 | 0.23 | 0.892 | NA | NA | 0.04 | 0.979 | 0.24 | 0.887 |  |
|  | SR x L x E | NA | NA | 1.08 | 0.584 | 1.08 | 0.584 | NA | NA | 0.94 | 0.625 | 0.94 | 0.625 |  |
|  | R2 | 0.05/NA | | 0.16/NA | | 0.16/NA | | 0.06/NA | | 0.16/NA | | 0.16/NA | |  |
| monoterpene | Veg. height | NA | NA | NA | NA | NA | NA | 0.58 | 0.448 | 0.58 | 0.448 | 0.58 | 0.448 | decreased with increasing species richness only in environments with soil history |
|  | Species richness (SR) | 2.08 | 0.149 | 2.08 | 0.149 | 2.08 | 0.149 | **5.11** | **0.024** | **5.11** | **0.024** | **6.13** | **0.013** |  |
|  | Legume (L) | NA | NA | 0.00 | 0.994 | 0.00 | 0.954 | NA | NA | 1.10 | 0.293 | 0.09 | 0.764 |  |
|  | Environment (E) | 0.26 | 0.879 | 0.26 | 0.879 | 0.26 | 0.879 | 0.02 | 0.991 | 0.00 | 0.998 | 0.00 | 0.998 |  |
|  | SR x L | NA | NA | 0.83 | 0.361 | 0.83 | 0.361 | NA | NA | 1.61 | 0.205 | 1.61 | 0.205 |  |
|  | SR x E | **8.53** | **0.014** | **9.25** | **0.010** | **7.23** | **0.027** | **7.31** | **0.026** | **8.38** | **0.015** | **6.26** | **0.044** |  |
|  | L x E | NA | NA | 2.69 | 0.260 | 4.71 | *0.095* | NA | NA | 2.25 | 0.325 | 4.37 | 0.113 |  |
|  | SR x L x E | NA | NA | 2.83 | 0.243 | 2.83 | 0.243 | NA | NA | 1.90 | 0.388 | 1.90 | 0.388 |  |
|  | R2 | 0.14/NA | | 0.19/NA | | 0.19/NA | | 0.14/0.22 | | 0.22/NA | | 0.22/NA | |  |
| sesquiterpene | Veg. height | NA | NA | NA | NA | NA | NA | 1.75 | 0.186 | 1.75 | 0.186 | 1.75 | 0.186 | decreased with species richness when considering the surrounding vegetation height |
|  | Species richness (SR) | 2.03 | 0.154 | 2.03 | 0.154 | 2.07 | 0.151 | **6.84** | **0.009** | **6.84** | **0.009** | **7.27** | **0.007** |  |
|  | Legume (L) | NA | NA | 0.25 | 0.615 | 0.21 | 0.644 | NA | NA | 0.47 | 0.493 | 0.04 | 0.837 |  |
|  | Environment (E) | 0.53 | 0.767 | 0.58 | 0.749 | 0.58 | 0.749 | 0.80 | 0.670 | 0.78 | 0.675 | 0.78 | 0.675 |  |
|  | SR x L | NA | NA | 0.62 | 0.430 | 0.62 | 0.430 | NA | NA | 1.95 | 0.163 | 1.95 | 0.163 |  |
|  | SR x E | 4.08 | 0.130 | 3.96 | 0.138 | 2.69 | 0.261 | 3.74 | 0.154 | 4.00 | 0.136 | 2.58 | 0.275 |  |
|  | L x E | NA | NA | 6.69 | **0.035** | 7.96 | **0.019** | NA | NA | 5.25 | *0.072* | **6.67** | **0.036** |  |
|  | SR x L x E | NA | NA | 4.79 | *0.091* | 4.79 | *0.091* | NA | NA | 2.61 | 0.272 | 2.61 | 0.272 |  |
|  | R2 | 0.09/0.17 | | 0.2/0.25 | | 0.2/0.25 | | 0.17/NA | | 0.25/NA | | 0.25/NA | |  |
| other | Veg. height | NA | NA | NA | NA | NA | NA | 0.33 | 0.567 | 0.33 | 0.567 | 0.33 | 0.567 |  |
|  | Species richness (SR) | **4.68** | **0.030** | **4.68** | **0.030** | **5.01** | **0.025** | **8.44** | **0.004** | **8.44** | **0.004** | **8.82** | **0.003** |  |
|  | Legume (L) | NA | NA | 0.33 | 0.563 | 0.01 | 0.941 | NA | NA | 0.46 | 0.499 | 0.07 | 0.788 |  |
|  | Environment (E) | 1.17 | 0.556 | 1.14 | 0.564 | 1.14 | 0.564 | 2.01 | 0.366 | 2.28 | 0.319 | 2.28 | 0.319 |  |
|  | SR x L | NA | NA | 0.18 | 0.672 | 0.18 | 0.672 | NA | NA | 0.00 | 0.987 | 0.00 | 0.987 |  |
|  | SR x E | 2.71 | 0.258 | 2.48 | 0.289 | 1.74 | 0.419 | 1.79 | 0.409 | 1.82 | 0.402 | 1.05 | 0.591 |  |
|  | L x E | NA | NA | 2.26 | 0.324 | 3.00 | 0.223 | NA | NA | 1.89 | 0.388 | 2.67 | 0.264 |  |
|  | SR x L x E | NA | NA | 1.12 | 0.571 | 1.12 | 0.571 | NA | NA | 1.33 | 0.514 | 1.33 | 0.514 |  |
|  | R2 | 0.15/NA | | 0.19/NA | | 0.19/NA | | 0.2/NA | | 0.23/NA | | 0.23/NA | |  |
| total | Veg. height | NA | NA | NA | NA | NA | NA | 0.89 | 0.345 | 0.89 | 0.345 | 0.89 | 0.345 |  |
|  | Species richness (SR) | 3.70 | *0.054* | 3.70 | *0.054* | 2.51 | 0.113 | 2.94 | *0.087* | 2.94 | *0.087* | **4.39** | **0.036** |  |
|  | Legume (L) | NA | NA | 1.58 | 0.209 | 2.77 | *0.096* | NA | NA | 3.33 | *0.068* | 1.88 | 0.170 |  |
|  | Environment (E) | 1.09 | 0.579 | 1.39 | 0.499 | 1.39 | 0.499 | 0.98 | 0.612 | 0.87 | 0.649 | 0.87 | 0.649 |  |
|  | SR x L | NA | NA | 0.51 | 0.475 | 0.51 | 0.475 | NA | NA | 0.93 | 0.335 | 0.93 | 0.335 |  |
|  | SR x E | 1.92 | 0.384 | 2.64 | 0.267 | 1.93 | 0.381 | 1.91 | 0.385 | 2.34 | 0.311 | 1.62 | 0.444 |  |
|  | L x E | NA | NA | 0.34 | 0.842 | 1.05 | 0.591 | NA | NA | 0.45 | 0.798 | 1.17 | 0.558 |  |
|  | SR x L x E | NA | NA | 1.56 | 0.459 | 1.56 | 0.459 | NA | NA | 1.00 | 0.607 | 1.00 | 0.607 |  |
|  | R2 | 0.08/NA | | 0.13/NA | | 0.13/NA | | 0.08/NA | | 0.14/NA | | 0.14/NA | |  |
| **Volatile compound diversity** | | | | | | | | | | | | | | |
| richness | Veg. height | NA | NA | NA | NA | NA | NA | 0.08 | 0.778 | 0.08 | 0.778 | 0.08 | 0.778 |  |
|  | Species richness (SR) | 1.57 | 0.210 | 1.57 | 0.210 | 2.17 | 0.141 | 3.50 | *0.061* | 3.50 | *0.061* | 3.33 | *0.068* |  |
|  | Legume (L) | NA | NA | 0.90 | 0.342 | 0.31 | 0.581 | NA | NA | 0.06 | 0.811 | 0.23 | 0.634 |  |
|  | Environment (E) | 0.50 | 0.780 | 0.54 | 0.763 | 0.54 | 0.763 | 0.34 | 0.842 | 0.37 | 0.829 | 0.37 | 0.829 |  |
|  | SR x L | NA | NA | 0.35 | 0.553 | 0.35 | 0.553 | NA | NA | 0.83 | 0.363 | 0.83 | 0.363 |  |
|  | SR x E | 1.98 | 0.371 | 1.84 | 0.398 | 2.25 | 0.324 | 1.84 | 0.398 | 1.75 | 0.417 | 2.05 | 0.359 |  |
|  | L x E | NA | NA | 1.68 | 0.432 | 1.27 | 0.531 | NA | NA | 1.21 | 0.547 | 0.90 | 0.637 |  |
|  | SR x L x E | NA | NA | 0.53 | 0.766 | 0.53 | 0.766 | NA | NA | 0.56 | 0.754 | 0.56 | 0.754 |  |
|  | R2 | 0.08/NA | | 0.1/NA | | 0.1/NA | | 0.09/NA | | 0.1/NA | | 0.1/NA | |  |
| shannon | Veg. height | NA | NA | NA | NA | NA | NA | 0.01 | 0.925 | 0.00 | 0.967 | 0.00 | 0.967 | decreased with increasing species richness only in phytometers that grew in environments with soil history (S+P+, S+P-), but remained similar across the species richness gradient in environments without history |
|  | Species richness (SR) | 0.26 | 0.612 | 0.18 | 0.669 | 0.41 | 0.520 | 0.46 | 0.495 | 0.26 | 0.612 | 0.23 | 0.633 |  |
|  | Legume (L) | NA | NA | 0.79 | 0.374 | 0.56 | 0.455 | NA | NA | 0.72 | 0.395 | 0.75 | 0.385 |  |
|  | Environment (E) | 2.50 | 0.287 | 1.63 | 0.442 | 1.63 | 0.442 | 3.24 | 0.198 | 1.69 | 0.429 | 1.69 | 0.429 |  |
|  | SR x L | NA | NA | 0.06 | 0.814 | 0.06 | 0.814 | NA | NA | 0.08 | 0.773 | 0.08 | 0.773 |  |
|  | SR x E | 3.13 | 0.209 | 2.47 | 0.290 | 2.21 | 0.332 | **5.55** | **0.048** | **5.37** | **0.049** | **2.08** | **0.354** |  |
|  | L x E | NA | NA | 0.03 | 0.983 | 0.30 | 0.861 | NA | NA | 0.03 | 0.983 | 0.33 | 0.849 |  |
|  | SR x L x E | NA | NA | 0.99 | 0.610 | 0.99 | 0.610 | NA | NA | 1.00 | 0.605 | 1.00 | 0.605 |  |
|  | R2 | 0.08/NA | | 0.1/NA | | 0.1/NA | | 0.09/NA | | 0.1/NA | | 0.1/NA | |  |
| simpson | Veg. height | NA | NA | NA | NA | NA | NA | 0.01 | 0.932 | 0.01 | 0.932 | 0.01 | 0.932 |  |
|  | Species richness (SR) | 0.07 | 0.786 | 0.07 | 0.786 | 0.27 | 0.601 | 0.07 | 0.786 | 0.07 | 0.786 | 0.08 | 0.773 |  |
|  | Legume (L) | NA | NA | 1.04 | 0.307 | 0.84 | 0.359 | NA | NA | 1.13 | 0.288 | 1.12 | 0.290 |  |
|  | Environment (E) | 3.28 | 0.194 | 3.38 | 0.184 | 3.38 | 0.184 | 3.70 | 0.157 | 3.36 | 0.186 | 3.36 | 0.186 |  |
|  | SR x L | NA | NA | 0.08 | 0.784 | 0.08 | 0.784 | NA | NA | 0.10 | 0.746 | 0.10 | 0.746 |  |
|  | SR x E | 2.17 | 0.338 | 2.02 | 0.364 | 1.40 | 0.498 | 1.82 | 0.402 | 1.92 | 0.383 | 1.30 | 0.521 |  |
|  | L x E | NA | NA | 0.40 | 0.818 | 1.03 | 0.599 | NA | NA | 0.40 | 0.818 | 1.02 | 0.601 |  |
|  | SR x L x E | NA | NA | 1.50 | 0.472 | 1.50 | 0.472 | NA | NA | 1.56 | 0.458 | 1.56 | 0.458 |  |
|  | R2 | 0.05/0.36 | | 0.14/NA | | 0.14/NA | | 0.05/0.34 | | 0.1/0.36 | | 0.1/0.36 | |  |

**Table S8. Community History Experiment: Wald-chi-squared analysis of variance (ANOVA) results for the linear mixed models of *selected* *Plantago lanceolata* phytometers across a diversity gradient in different community history environments based on** **untargeted metabolome diversity**.

The effects of vegetation height, species richness, environment treatment (*S+P+*, *S+P-*, *S-P-*) and legumes (presence or absence) on metabolome diversity were tested using mixed-effects models. *Community History Experiment* compared the metabolomic profiles of *selected* phytometers grown in different environment treatments based on the ΔBEF experiment established in 2016 (Vogel et al., 2019). Six models were run to disentangle the confounding effects: Model 1 examined species richness, environment treatment, and their interaction. Models 2 and 3 assessed legume presence, either before or after species richness. Models 4-6 tested vegetation height, including it as a covariate. All models used plot nested within block as random effects. The table shows Chi-square (X²) and p-values for fixed effects, with significant effects in bold (P< 0.05) and tendencies within brackets (P < 0.1). Data were transformed as needed to meet assumptions.

| **Variable** | **Factor** | **Model 1** | | **Model 2** | | **Model 3** | | **Model 4** | | **Model 5** | | **Model 6** | | **Pattern** |
| --- | --- | --- | --- | --- | --- | --- | --- | --- | --- | --- | --- | --- | --- | --- |
|  |  | y~ SR* E+  (1\| block/ plot) | | y~ SR*L* E+  (1\| block/ plot) | | y~ L*SR*E+  (1\| block/ plot) | | y~ VG+SR*E+  (1\| block/ plot) | | y~ VG+SR*L*E+  (1\| block/ plot) | | y~ VG+L*SR* E+  (1\| block/ plot) | |  |
|  |  | **X2** | **P** | **X2** | **P** | **X2** | **P** | **X2** | **P** | **X2** | **P** | **X2** | **P** |  |
| Metabolome richness Hill q0 negative binomial | Veg. height | NA | NA | NA | NA | NA | NA | 0.61 | 0.437 | 0.61 | 0.437 | 0.61 | 0.437 |  |
|  | Species richness (SR) | 0.02 | 0.888 | 0.02 | 0.888 | 0.02 | 0.884 | 0.14 | 0.713 | 0.14 | 0.713 | 0.15 | 0.703 |  |
|  | Legume (L) | NA | NA | 0.00 | 0.971 | 0.00 | 0.998 | NA | NA | 0.22 | 0.638 | 0.21 | 0.646 |  |
|  | Environment (E) | 0.42 | 0.809 | 0.42 | 0.810 | 0.42 | 0.810 | 0.69 | 0.707 | 0.72 | 0.696 | 0.72 | 0.696 |  |
|  | SR x L | NA | NA | 3.12 | 0.078 | 3.12 | 0.078 | NA | NA | 3.20 | 0.074 | 3.20 | 0.074 |  |
|  | SR x E | 0.38 | 0.827 | 0.34 | 0.842 | 0.40 | 0.819 | 0.50 | 0.777 | 0.45 | 0.800 | 0.48 | 0.787 |  |
|  | L x E | NA | NA | 0.56 | 0.756 | 0.50 | 0.778 | NA | NA | 0.44 | 0.803 | 0.41 | 0.817 |  |
|  | SR x L x E | NA | NA | 1.63 | 0.442 | 1.63 | 0.442 | NA | NA | 1.38 | 0.501 | 1.38 | 0.501 |  |
|  | R2 |  | |  | |  | |  | |  | |  | |  |
| Shannon metabolome diversity Hill q1 | Veg. height | NA | NA | NA | NA | NA | NA | 0.05 | 0.831 | 0.05 | 0.831 | 0.05 | 0.831 | Increased with increasing species richness only in selected phytometers that grew in the environment of origin |
|  | Species richness (SR) | 0.03 | 0.859 | 0.03 | 0.859 | 0.04 | 0.840 | 0.34 | 0.560 | 0.34 | 0.560 | 0.31 | 0.575 |  |
|  | Legume (L) | NA | NA | 0.06 | 0.801 | 0.05 | 0.816 | NA | NA | 0.00 | 0.998 | 0.03 | 0.872 |  |
|  | Environment (E) | 1.69 | 0.430 | 1.64 | 0.440 | 1.64 | 0.440 | 1.43 | 0.488 | 1.43 | 0.488 | 1.43 | 0.488 |  |
|  | SR x L | NA | NA | 1.60 | 0.207 | 1.60 | 0.207 | NA | NA | 1.54 | 0.214 | 1.54 | 0.214 |  |
|  | SR x E | **10.36** | **0.006** | **10.43** | **0.005** | **9.55** | **0.008** | **10.60** | **0.005** | **10.43** | **0.005** | **9.45** | **0.009** |  |
|  | L x E | NA | NA | 0.13 | 0.935 | 1.01 | 0.603 | NA | NA | 0.14 | 0.935 | 1.11 | 0.574 |  |
|  | SR x L x E | NA | NA | 0.88 | 0.645 | 0.88 | 0.645 | NA | NA | 0.87 | 0.647 | 0.87 | 0.647 |  |
|  | R2 | 0.08/0.16 | | 0.1/0.16 | | 0.1/0.16 | | 0.08/NA | | 0.1/0.16 | | 0.1/0.16 | |  |
| Simpson metabolome diversity Hill q2 | Veg. height | NA | NA | NA | NA | NA | NA | 0.00 | 0.971 | 0.00 | 0.971 | 0.00 | 0.971 | Increased with increasing species richness only in selected phytometers that grew in the environment of origin |
|  | Species richness (SR) | 0.22 | 0.638 | 0.22 | 0.638 | 0.21 | 0.646 | 0.63 | 0.428 | 0.63 | 0.428 | 0.65 | 0.419 |  |
|  | Legume (L) | NA | NA | 0.00 | 0.989 | 0.01 | 0.921 | NA | NA | 0.04 | 0.839 | 0.02 | 0.900 |  |
|  | Environment (E) | 1.62 | 0.446 | 1.62 | 0.446 | 1.62 | 0.446 | 1.64 | 0.440 | 1.68 | 0.433 | 1.68 | 0.433 |  |
|  | SR x L | NA | NA | 2.44 | 0.118 | 2.44 | 0.118 | NA | NA | 2.30 | 0.130 | 2.30 | 0.130 |  |
|  | SR x E | **8.23** | **0.016** | **8.39** | **0.015** | **7.78** | **0.020** | **7.96** | **0.019** | **8.16** | **0.017** | **7.49** | **0.024** |  |
|  | L x E | NA | NA | 0.14 | 0.933 | 0.74 | 0.689 | NA | NA | 0.12 | 0.942 | 0.79 | 0.673 |  |
|  | SR x L x E | NA | NA | 1.34 | 0.511 | 1.34 | 0.511 | NA | NA | 1.27 | 0.531 | 1.27 | 0.531 |  |
|  | R2 | 0.07/0.14 | | 0.1/0.14 | | 0.1/0.14 | | 0.07/NA | | 0.1/0.15 | | 0.1/0.15 | |  |

**Table S9. Community History Experiment: Wald-chi-squared analysis of variance (ANOVA) results for the linear mixed models of *selected Plantago lanceolata* phytometers across a diversity gradient in different community history environments based** **on targeted defense compounds**.

The effects of vegetation height, species richness, *experimental* environment *(S+P+, S+P-, S-P-)* and legumes (presence or absence) on targeted defense compounds were tested using mixed-effects models. *Community History Experiment* compared the metabolomic profiles of *selected* phytometers grown in different environment treatments based on the ΔBEF experiment established in 2016 (Vogel et al., 2019). Six models were run to disentangle the confounding effects: Model 1 examined species richness, environment treatment, and their interaction. Models 2 and 3 assessed legume presence, either before or after species richness. Models 4-6 tested vegetation height, including it as a covariate. All models used plot nested within block as random effects. The table shows Chi-square (X²) and p-values for fixed effects, with significant effects in bold (P< 0.05) and tendencies within brackets (P < 0.1). Data were transformed as needed to meet assumptions. N = 163.

| **Variable** | **Factor** | **Model 1** | | **Model 2** | | **Model 3** | | **Model 4** | | **Model 5** | | **Model 6** | | **Pattern** |
| --- | --- | --- | --- | --- | --- | --- | --- | --- | --- | --- | --- | --- | --- | --- |
|  |  | y~ SR* E+  (1\| block/ plot) | | y~ SR*L* E+  (1\| block/ plot) | | y~ L*SR*E+  (1\| block/ plot) | | y~ VG+SR*E+  (1\| block/ plot) | | y~ VG+SR*L*E+  (1\| block/ plot) | | y~ VG+L*SR* E+  (1\| block/ plot) | |  |
|  |  | **X2** | **P** | **X2** | **P** | **X2** | **P** | **X2** | **P** | **X2** | **P** | **X2** | **P** |  |
| **Defense hormones** | | | | | | | | | | | | | | |
| Jasmonic acid  (ng (gdw)-1) log10 | Veg. height | NA | NA | NA | NA | NA | NA | [3.38] | *0.066* | [3.38] | *0.066* | [3.38] | *0.066* |  |
|  | Species richness (SR) | 0.03 | 0.871 | 0.03 | 0.871 | 0.04 | 0.833 | 1.04 | 0.309 | 1.04 | 0.309 | 0.99 | 0.320 |  |
|  | Legume (L) | NA | NA | 0.05 | 0.820 | 0.03 | 0.854 | NA | NA | 0.00 | 0.955 | 0.05 | 0.827 |  |
|  | Environment (E) | 2.09 | 0.352 | 2.11 | 0.348 | 2.11 | 0.348 | 2.07 | 0.356 | 2.07 | 0.356 | 2.07 | 0.356 |  |
|  | SR x L | NA | NA | 0.45 | 0.503 | 0.45 | 0.503 | NA | NA | 0.44 | 0.508 | 0.44 | 0.508 |  |
|  | SR x E | [5.19] | *0.075* | [5.19] | *0.075* | [5.19] | *0.075* | 4.30 | 0.116 | 4.30 | 0.117 | 4.46 | 0.107 |  |
|  | L x E | NA | NA | 1.47 | 0.480 | 1.19 | 0.553 | NA | NA | 0.74 | 0.690 | 0.58 | 0.750 |  |
|  | SR x L x E | NA | NA | 2.12 | 0.346 | 2.12 | 0.346 | NA | NA | 1.52 | 0.467 | 1.52 | 0.467 |  |
|  | R2 | 0.03/0.32 | | 0.06/0.33 | | 0.06/0.33 | | 0.12/NA | | 0.14/NA | | 0.14/NA | |  |
| Jasmonic acid isoleucine (JA-Ile) | Veg. height | NA | NA | NA | NA | NA | NA | 0.95 | 0.330 | 0.95 | 0.330 | 0.95 | 0.330 |  |
|  | Species richness (SR) | 1.75 | 0.185 | 1.75 | 0.185 | 1.88 | 0.171 | [3.36] | *0.067* | [3.36] | *0.067* | 3.36 | *0.067* |  |
|  | Legume (L) | NA | NA | 0.15 | 0.699 | 0.03 | 0.872 | NA | NA | 0.02 | 0.889 | 0.01 | 0.910 |  |
|  | Environment (E) | 0.67 | 0.717 | 0.69 | 0.708 | 0.69 | 0.708 | 0.49 | 0.782 | 0.50 | 0.779 | 0.50 | 0.779 |  |
|  | SR x L | NA | NA | 1.08 | 0.299 | 1.08 | 0.299 | NA | NA | 0.88 | 0.348 | 0.88 | 0.348 |  |
|  | SR x E | **6.29** | **0.043** | **6.23** | **0.044** | **10.03** | **0.007** | [5.53] | *0.063* | [5.56] | *0.062* | **9.11** | **0.011** |  |
|  | L x E | NA | NA | **6.49** | **0.039** | 2.70 | 0.260 | NA | NA | [5.88] | *0.053* | 2.33 | 0.312 |  |
|  | SR x L x E | NA | NA | 3.68 | 0.159 | 3.68 | 0.159 | NA | NA | 3.18 | 0.204 | 3.18 | 0.204 |  |
|  | R2 | 0.07/0.22 | | 0.12/0.25 | | 0.12/0.25 | | 0.11/NA | | 0.14/0.26 | | 0.14/0.26 | |  |
| 12-hydroxy-jasmonic acid (OH-JA) | Veg. height | NA | NA | NA | NA | NA | NA | 0.14 | 0.708 | 0.14 | 0.708 | 0.14 | 0.708 | decresed with increasing species richness in communities without legumes |
|  | Species richness (SR) | 2.05 | 0.152 | 2.05 | 0.152 | [3.27] | *0.071* | 2.52 | 0.113 | 2.52 | 0.113 | [2.97] | *0.085* |  |
|  | Legume (L) | NA | NA | 1.89 | 0.169 | 0.67 | 0.413 | NA | NA | 1.30 | 0.254 | 0.85 | 0.356 |  |
|  | Environment (E) | 1.58 | 0.455 | 1.65 | 0.437 | 1.65 | 0.437 | 1.27 | 0.531 | 1.69 | 0.430 | 1.69 | 0.430 |  |
|  | SR x L | NA | NA | **7.96** | **0.005** | **7.96** | **0.005** | NA | NA | **7.98** | **0.005** | **7.98** | **0.005** |  |
|  | SR x E | 3.05 | 0.217 | 3.25 | 0.197 | 3.13 | 0.209 | 2.86 | 0.239 | 3.20 | 0.202 | 3.10 | 0.212 |  |
|  | L x E | NA | NA | 0.96 | 0.618 | 1.08 | 0.583 | NA | NA | 0.94 | 0.624 | 1.05 | 0.593 |  |
|  | SR x L x E | NA | NA | 0.87 | 0.647 | 0.87 | 0.647 | NA | NA | 0.87 | 0.648 | 0.87 | 0.648 |  |
|  | R2 | 0.07/0.2 | | 0.15/NA | | 0.15/NA | | 0.06/0.19 | | 0.15/NA | | 0.15/NA | |  |
| 12-hydroxy-jasmonoyl-isoleucine  (12OH-JA-Ile) | Veg. height | NA | NA | NA | NA | NA | NA | 1.17 | 0.279 | 1.17 | 0.279 | 1.17 | 0.279 | decresed with increasing species richness only in S+P- |
|  | Species richness (SR) | [3.65] | *0.056* | [3.65] | *0.056* | **3.87** | **0.049** | **5.94** | **0.015** | **5.94** | **0.015** | **5.86** | **0.015** |  |
|  | Legume (L) | NA | NA | 0.22 | 0.636 | 0.00 | 0.964 | NA | NA | 0.01 | 0.926 | 0.09 | 0.767 |  |
|  | Environment (E) | 0.46 | 0.793 | 0.46 | 0.795 | 0.46 | 0.795 | 0.15 | 0.926 | 0.15 | 0.927 | 0.15 | 0.927 |  |
|  | SR x L | NA | NA | 0.22 | 0.637 | 0.22 | 0.637 | NA | NA | 0.21 | 0.650 | 0.21 | 0.650 |  |
|  | SR x E | **7.25** | **0.027** | **7.16** | **0.028** | **7.92** | **0.019** | **6.53** | **0.038** | **6.52** | **0.038** | **7.21** | **0.027** |  |
|  | L x E | NA | NA | 2.78 | 0.249 | 2.03 | 0.363 | NA | NA | 2.16 | 0.340 | 1.46 | 0.481 |  |
|  | SR x L x E | NA | NA | 1.42 | 0.492 | 1.42 | 0.492 | NA | NA | 1.17 | 0.558 | 1.17 | 0.558 |  |
|  | R2 | 0.12/0.29 | | 0.14/0.3 | | 0.14/0.3 | | 0.18/NA | | 0.19/NA | | 0.19/NA | |  |
| 12-carboxy-jasmonoyl-L-isoleucine (COOH-JA-Ile) glmer | Veg. height | NA | NA | NA | NA | NA | NA | **5.67** | **0.017** | **5.67** | **0.017** | **5.67** | **0.017** | decrease with increasing vegetation height |
|  | Species richness (SR) | 0.78 | 0.378 | 0.78 | 0.378 | 0.77 | 0.380 | 2.47 | 0.116 | 2.47 | 0.116 | 2.07 | 0.150 |  |
|  | Legume (L) | NA | NA | 0.01 | 0.908 | 0.02 | 0.893 | NA | NA | 0.63 | 0.426 | 1.03 | 0.309 |  |
|  | Environment (E) | 2.51 | 0.284 | 2.51 | 0.285 | 2.51 | 0.285 | 0.30 | 0.861 | 0.20 | 0.904 | 0.20 | 0.904 |  |
|  | SR x L | NA | NA | 0.01 | 0.940 | 0.01 | 0.940 | NA | NA | 0.00 | 0.966 | 0.00 | 0.966 |  |
|  | SR x E | 0.53 | 0.767 | 0.54 | 0.763 | 0.50 | 0.778 | 0.26 | 0.878 | 0.27 | 0.875 | 0.36 | 0.834 |  |
|  | L x E | NA | NA | **11.42** | **0.003** | **11.46** | **0.003** | NA | NA | **7.63** | **0.022** | **7.54** | **0.023** |  |
|  | SR x L x E | NA | NA | 1.64 | 0.44 | 1.64 | 0.44 | NA | NA | 1.82 | 0.402 | 1.82 | 0.402 |  |
|  | R2 | 0.32/0.99 | | 0.48/0.99 | | 0.48/0.99 | | 0.47/0.99 | | 0.5/0.99 | | 0.98/NA | |  |
| Total jasmonates | Veg. height | NA | NA | NA | NA | NA | NA | 2.27 | 0.132 | 2.27 | 0.132 | 2.27 | 0.132 |  |
|  | Species richness (SR) | 0.83 | 0.361 | 0.83 | 0.361 | 0.82 | 0.366 | 2.32 | 0.128 | 2.32 | 0.128 | 2.02 | 0.155 |  |
|  | Legume (L) | NA | NA | 0.11 | 0.744 | 0.12 | 0.725 | NA | NA | 0.37 | 0.541 | 0.67 | 0.412 |  |
|  | Environment (E) | 1.87 | 0.393 | 1.84 | 0.398 | 1.84 | 0.398 | 1.01 | 0.603 | 0.93 | 0.628 | 0.93 | 0.628 |  |
|  | SR x L | NA | NA | **4.45** | **0.035** | **4.45** | **0.035** | NA | NA | [3.71] | *0.054* | [3.71] | *0.054* |  |
|  | SR x E | 4.00 | 0.136 | 4.03 | 0.134 | 4.31 | 0.116 | 3.37 | 0.186 | 3.45 | 0.178 | 3.65 | 0.161 |  |
|  | L x E | NA | NA | 1.75 | 0.416 | 1.47 | 0.479 | NA | NA | 1.18 | 0.555 | 0.97 | 0.615 |  |
|  | SR x L x E | NA | NA | 2.09 | 0.352 | 2.09 | 0.352 | NA | NA | 2.36 | 0.307 | 2.36 | 0.307 |  |
|  | R2 | 0.05/0.28 | | 0.14/0.27 | | 0.14/0.27 | | 0.1/NA | | 0.19/NA | | 0.19/NA | |  |
| Abscisic acid (ABA) | Veg. height | NA | NA | NA | NA | NA | NA | 0.58 | 0.445 | 0.58 | 0.445 | 0.58 | 0.445 | decreased with increasing species richness only in P. lanceolata phytometers in environments with soil and plant history when considering the surrounding vegetation height |
|  | Species richness (SR) | 1.36 | 0.243 | 1.36 | 0.243 | 1.18 | 0.277 | 0.85 | 0.356 | 0.85 | 0.356 | 0.83 | 0.361 |  |
|  | Legume (L) | NA | NA | 0.03 | 0.852 | 0.22 | 0.641 | NA | NA | 0.01 | 0.919 | 0.03 | 0.862 |  |
|  | Environment (E) | 1.12 | 0.571 | 1.12 | 0.573 | 1.12 | 0.573 | 1.29 | 0.524 | 1.28 | 0.526 | 1.28 | 0.526 |  |
|  | SR x L | NA | NA | 2.20 | 0.138 | 2.20 | 0.138 | NA | NA | 2.44 | 0.119 | 2.44 | 0.119 |  |
|  | SR x E | **5.94** | **0.048** | **5.94** | **0.048** | [4.84] | *0.089* | **6.26** | **0.044** | **6.38** | **0.041** | [5.26] | *0.072* |  |
|  | L x E | NA | NA | 0.38 | 0.828 | 1.52 | 0.467 | NA | NA | 0.35 | 0.839 | 1.47 | 0.479 |  |
|  | SR x L x E | NA | NA | 0.27 | 0.873 | 0.27 | 0.873 | NA | NA | 0.53 | 0.768 | 0.53 | 0.768 |  |
|  | R2 | 0.07/NA | | 0.1/NA | | 0.1/NA | | 0.07/NA | | 0.11/NA | | 0.11/NA | |  |
| Salicylic acid (SA) | Veg. height | NA | NA | NA | NA | NA | NA | [3.43] | *0.064* | [3.43] | *0.064* | 1.00 | 0.317 | Decreased with increasing species richness in non-legume communities. In plots with legumes, only increased with sp. richness in the environment with no history environment whereas in other environments there was no effect of species richness |
|  | Species richness (SR) | **5.59** | **0.018** | **5.59** | **0.018** | **5.35** | **0.021** | [3.56] | *0.059* | [3.56] | *0.059* | 13.41 | **< 0.001** |  |
|  | Legume (L) | NA | NA | 0.01 | 0.934 | 0.24 | 0.623 | NA | NA | 0.23 | 0.629 | 0.14 | 0.707 |  |
|  | Environment (E) | 1.81 | 0.405 | 1.82 | 0.402 | 1.82 | 0.402 | 1.31 | 0.519 | 1.30 | 0.522 | 2.89 | 0.236 |  |
|  | SR x L | NA | NA | **8.82** | **0.003** | **8.82** | **0.003** | NA | NA | **10.46** | **0.001** | **11.96** | **0.001** |  |
|  | SR x E | 0.79 | 0.673 | 0.91 | 0.633 | 0.56 | 0.756 | 1.02 | 0.601 | 1.17 | 0.557 | 1.77 | 0.413 |  |
|  | L x E | NA | NA | 1.21 | 0.545 | 1.57 | 0.457 | NA | NA | 1.46 | 0.482 | 3.07 | 0.216 |  |
|  | SR x L x E | NA | NA | **10.62** | **0.005** | **10.62** | **0.005** | NA | NA | **10.89** | **0.004** | **6.69** | **0.035** |  |
|  | R2 | 0.16/0.33 | | 0.29/0.35 | | 0.29/0.35 | | 0.18/0.33 | | 0.31/0.34 | | 0.02/NA | |  |
| **Irioid glycosides** | | | | | | | | | | | | | | |
| Aucubin | Veg. height | NA | NA | NA | NA | NA | NA | [3.56] | *0.059* | [3.56] | *0.059* | [3.56] | *0.059* | decreased as plant species richness increased when vegetation height in the surrounding was taken into consideration |
|  | Species richness (SR) | 0.94 | 0.333 | 0.94 | 0.333 | 1.29 | 0.256 | **6.32** | **0.012** | **6.32** | **0.012** | **5.97** | **0.015** |  |
|  | Legume (L) | NA | NA | 1.98 | 0.160 | 1.63 | 0.202 | NA | NA | 0.11 | 0.735 | 0.47 | 0.493 |  |
|  | Environment (E) | 1.94 | 0.379 | 1.84 | 0.399 | 1.84 | 0.399 | 2.71 | 0.258 | 2.63 | 0.268 | 2.63 | 0.268 |  |
|  | SR x L | NA | NA | 0.04 | 0.840 | 0.04 | 0.840 | NA | NA | 0.00 | 0.946 | 0.00 | 0.946 |  |
|  | SR x E | 0.55 | 0.760 | 0.50 | 0.777 | 0.42 | 0.812 | 0.30 | 0.859 | 0.31 | 0.855 | 0.21 | 0.902 |  |
|  | L x E | NA | NA | 0.12 | 0.940 | 0.21 | 0.900 | NA | NA | 0.30 | 0.859 | 0.41 | 0.814 |  |
|  | SR x L x E | NA | NA | 3.9 | 0.142 | 3.9 | 0.142 | NA | NA | 3.75 | 0.153 | 3.75 | 0.153 |  |
|  | R2 | 0.03/0.12 | | 0.06/0.14 | | 0.06/0.14 | | 0.1/NA | | 0.13/NA | | 0.13/NA | |  |
| Catalpol | Veg. height | NA | NA | NA | NA | NA | NA | **9.77** | **0.002** | **9.77** | **0.002** | **9.77** | **0.002** | Increased with increasing vegetation height |
|  | Species richness (SR) | 1.19 | 0.276 | 1.19 | 0.276 | 1.59 | 0.207 | 0.58 | 0.445 | 0.58 | 0.445 | 0.84 | 0.360 |  |
|  | Legume (L) | NA | NA | 0.43 | 0.510 | 0.03 | 0.866 | NA | NA | **4.14** | **0.042** | **3.89** | **0.049** |  |
|  | Environment (E) | **6.27** | **0.043** | **6.26** | **0.044** | **6.26** | **0.044** | 4.68 | *0.097* | 4.84 | *0.089* | 4.84 | *0.089* |  |
|  | SR x L | NA | NA | 0.54 | 0.462 | 0.54 | 0.462 | NA | NA | 0.60 | 0.440 | 0.60 | 0.440 |  |
|  | SR x E | 2.54 | 0.280 | 2.51 | 0.285 | 2.13 | 0.344 | 1.77 | 0.412 | 1.82 | 0.402 | 1.50 | 0.472 |  |
|  | L x E | NA | NA | 0.46 | 0.795 | 0.84 | 0.657 | NA | NA | 0.77 | 0.680 | 1.09 | 0.579 |  |
|  | SR x L x E | NA | NA | 4.07 | 0.131 | 4.07 | 0.131 | NA | NA | 2.76 | 0.251 | 2.76 | 0.251 |  |
|  | R2 | 0.07/0.12 | | 0.1/0.16 | | 0.1/0.16 | | 0.12/NA | | 0.17/NA | | 0.17/NA | |  |
| **Phenylpropanoid glycosides** | | | | | | | | | | | | | | |
| Plantamajoside | Veg. height | NA | NA | NA | NA | NA | NA | 1.11 | 0.292 | 1.11 | 0.292 | 1.11 | 0.292 | decreased with species richness only in communities with legumes |
|  | Species richness (SR) | 0.67 | 0.412 | 0.67 | 0.412 | 0.67 | 0.412 | 0.17 | 0.679 | 0.17 | 0.679 | 0.12 | 0.733 |  |
|  | Legume (L) | NA | NA | 0.00 | 0.977 | 0.00 | 0.992 | NA | NA | 0.08 | 0.777 | 0.14 | 0.713 |  |
|  | Environment (E) | 2.99 | 0.225 | 2.99 | 0.224 | 2.99 | 0.224 | 2.63 | 0.268 | 2.58 | 0.276 | 2.58 | 0.276 |  |
|  | SR x L | NA | NA | **9.07** | **0.003** | **9.07** | **0.003** | NA | NA | **9.07** | **0.003** | **9.07** | **0.003** |  |
|  | SR x E | 3.43 | 0.180 | 3.54 | 0.170 | 2.07 | 0.356 | 3.58 | 0.167 | 3.69 | 0.158 | 2.15 | 0.341 |  |
|  | L x E | NA | NA | 1.91 | 0.386 | 3.38 | 0.184 | NA | NA | 1.89 | 0.388 | 3.43 | 0.180 |  |
|  | SR x L x E | NA | NA | **11.04** | **0.004** | **11.04** | **0.004** | NA | NA | **10.65** | **0.005** | **10.65** | **0.005** |  |
|  | R2 | 0.05/NA | | 0.28/NA | | 0.28/NA | | 0.04/0.26 | | 0.28/NA | | 0.28/NA | |  |
| Verbascoside | Veg. height | NA | NA | NA | NA | NA | NA | 0.05 | 0.817 | 0.05 | 0.817 | 0.05 | 0.817 | Considering the presence of legumes, concentration increased with species richness in the presence of legumes but decreased in communities without legumes, regardless of the experimental environment |
|  | Species richness (SR) | 0.61 | 0.436 | 0.61 | 0.436 | 0.89 | 0.346 | 0.97 | 0.324 | 0.97 | 0.324 | 1.26 | 0.262 |  |
|  | Legume (L) | NA | NA | 0.47 | 0.492 | 0.19 | 0.662 | NA | NA | 0.28 | 0.595 | 0.00 | 1.000 |  |
|  | Environment (E) | 2.77 | 0.250 | 2.73 | 0.255 | 2.73 | 0.255 | 3.41 | 0.182 | 2.00 | 0.367 | 2.00 | 0.367 |  |
|  | SR x L | NA | NA | **8.22** | **0.004** | **8.22** | **0.004** | NA | NA | **10.04** | **0.002** | **10.04** | **0.002** |  |
|  | SR x E | 3.85 | 0.146 | 3.72 | 0.156 | 3.09 | 0.213 | 3.39 | 0.183 | 3.30 | 0.192 | 2.66 | 0.264 |  |
|  | L x E | NA | NA | 1.03 | 0.597 | 1.66 | 0.436 | NA | NA | 0.90 | 0.637 | 1.54 | 0.464 |  |
|  | SR x L x E | NA | NA | [5.24] | *0.073* | [5.24] | *0.073* | NA | NA | **6.47** | **0.039** | **6.47** | **0.039** |  |
|  | R2 | 0.06/NA | | 0.25/0.33 | | 0.25/0.33 | | 0.07/NA | | 0.28/NA | | 0.28/NA | |  |
